# Toward a first principles understanding of the activation and deactivation mechanisms of class A G-protein coupled receptors and voltage-gated cation channels

**DOI:** 10.1101/2022.03.29.486149

**Authors:** Hongbin Wan, Robert A. Pearlstein

## Abstract

We previously reported a first principles multi-scale theory called Biodynamics that attributes cellular functions to sets of coupled molecular and ionic fluxes operating in the non-equilibrium/non-linear dynamic regime. Fluxes build and decay over time and undergo dynamic non-covalent intra- and intermolecular state transitions powered principally by the storage and release of free energy to/from the H-bond networks of external and internal solvation (that we refer to as solvation dynamics) at rates governed by the desolvation and resolvation costs incurred during their entry and exit, respectively. We have thus far examined the functional state transitions of cereblon and COVID M^pro^ in this context, and now turn to the agonist-induced activating and deactivating state transitions of class A G-protein coupled receptors (GPCRs) and membrane potential-/dipole potential-induced activating and deactivating state transitions of voltage-gated cation channels (VGCCs). We analyzed crystal structures of the activated and deactivated forms of the human β_2_-adrenergic receptor (β_2_-AR) and cryo-EM structures of the activated and deactivated forms of Na_v_1.7 channels. We postulate that activation and deactivation of the β_2_-AR is conveyed by switchable changes in transmembrane helix (TMH) orientations relative to extracellular loop 2 (ECL2) and curvature of TMH6 and TMH7, all of which are powered by solvation free energy and kickstarted by agonist binding. The known activation and deactivation mechanisms of Na_v_1.7 consist of S4 translations toward and away from the extracellular membrane surface, respectively, resulting in S4-S5 linker repositioning, followed by rearrangements of the S5 and S6 helices. The latter TMH conveys channel opening and closing by respectively curving away from and toward the central pore axis. We postulate that all of these rearrangements are likewise powered by solvation free energy and kickstarted by changes in the membrane and dipole potentials. The results of our study may facilitate structure-based design of GPCR agonists/antagonists and mitigation of drug-induced ion channel blockade.

## Introduction

G-protein coupled receptors (GPCRs) transmit chemical signals (agonists) between their extracellular (EC) orthosteric and intracellular (IC) allosteric binding sites to which agonists and heterotrimeric GTP-binding proteins (G-proteins) respectively bind. Voltage-gated cation channels (VGCCs) transmit electrical signals (ion currents) between the extracellular (EC) and intracellular (IC) compartments via pores that open and close at channel-specific membrane voltages (Δψ_*m*_(*t*)). Many GPCRs and some VGCCs are constitutively active to varying degrees due to low barriers between their deactivated and activated states. The structure-function relationships of GPCRs and VGCCs have been investigated intensively for more than three decades, beginning with the pioneering work of Schertler,^1^ Baldwin,^2^ Weinstein,^3^ Ballesteros,^4^ and Kobilka^5^ on GPCRs and that of Guy^6^ and MacKinnon^7–9^ on VGCCs. However, the detailed activation and deactivation mechanisms, including the means by which they are powered, remain poorly understood.

Crystal structures of a growing number of class A agonist and antagonist-bound GPCRs have been solved, beginning with bovine rhodopsin in 2000^10,11^ All class A GPCRs are comprised of a single seven trans-membrane helical (7-TMH) bundle interconnected by three N-terminal EC loops (ECL1-3) and three C-terminal IC loops (ICL1-3) of varying length. GPCRs are activated by a wide variety of endogenous peptide, lipid, and small molecule agonists that bind within the 7-TMH bundle (buried deep within the bundle in some cases) or to separate EC N-terminal domains, case-by-case. Crystal and cryo-EM structures have likewise been solved for a growing number of VGCCs, beginning with the bacterial potassium (K_v_) channels KcsA in 1998^7^, MthK in 2002,^12^ and K_v_AP in 2005,^13^ and more recently, the human cardiac K_v_11.1 channel (hERG),^14^ the human cardiac sodium (Na_v_) channel Na_v_1.5,^15^ and the human neuronal channels Na_v_1.1,^16^ Na_v_1.4,^17^ and Na_v_1.7.^18^

We previously reported a first principles multi-scale theory called “Biodynamics” that addresses the general means by which cellular functions are conveyed by dynamic/non-equilibrium multi-species systems (the concentrations and state distributions of which build and decay over time), and by which such systems are powered by covalent and non-covalent free energy forms.^19–21^ As is well appreciated, the populations of non-covalent intra- and intermolecular states at static equilibrium are distributed according to their free energy <u>differences</u> (ΔG). However, under non-equilibrium conditions, molecular states are distributed according to their entry and exit free energy <u>barriers</u> 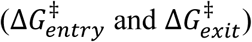 attributed by our theory principally to solvation free energy contributions, as follows:

1. ΔG^‡^ consists principally of the total cost of H-bond disruption (water-water and water-solute).
2. Under <u>aqueous</u> conditions, the free energy barrier to entering a given non-covalent state j from state i 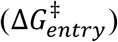 consists principally of the cost of <u>desolvating</u> H-bond enriched solvation in state i relative to bulk solvent, and the free energy barrier to exiting from state i back to state j 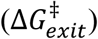 consists principally of the cost of <u>resolvating</u> state j at H-bond depleted solvation positions.
3. The rates of entry to and exit from each non-covalent state are proportional to 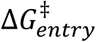 and 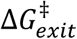, respectively.

GPCRs and VGCCs necessarily reside at the intersection between structural integrity/global stability and dynamic rearrangeability/local instability governed energetically by:

1. The balance between H-bond enriched/stabilizing and H-bond depleted/de-stabilizing solvation residing within:

a. Binding sites containing both H-bond enriched and depleted solvation.
b. Transient internal channels/cavities containing trapped or impeded H-bond depleted solvation that serve as free volumes into which intramolecular rearrangements proceed (noting that channels connect to the external surface whereas cavities do not). Internal solvation is present within both globular and membrane-bound proteins, including GPCRs (Figure 1) and VGCCs. Such solvation necessarily resides within a Goldilocks zone of positive free energy (too high and folding is precluded; too low and rearrangeability is precluded). Functional intramolecular rearrangements are powered principally by the release of this free energy via desolvation in cis (typically initiated by <u>intermolecular</u> rearrangements in trans).
2. Disruption of intra-helical H-bonds (IHHBs) within certain pairs of helical turns, which are energetically strong in the absence of bulk solvent.

**Figure 1.**
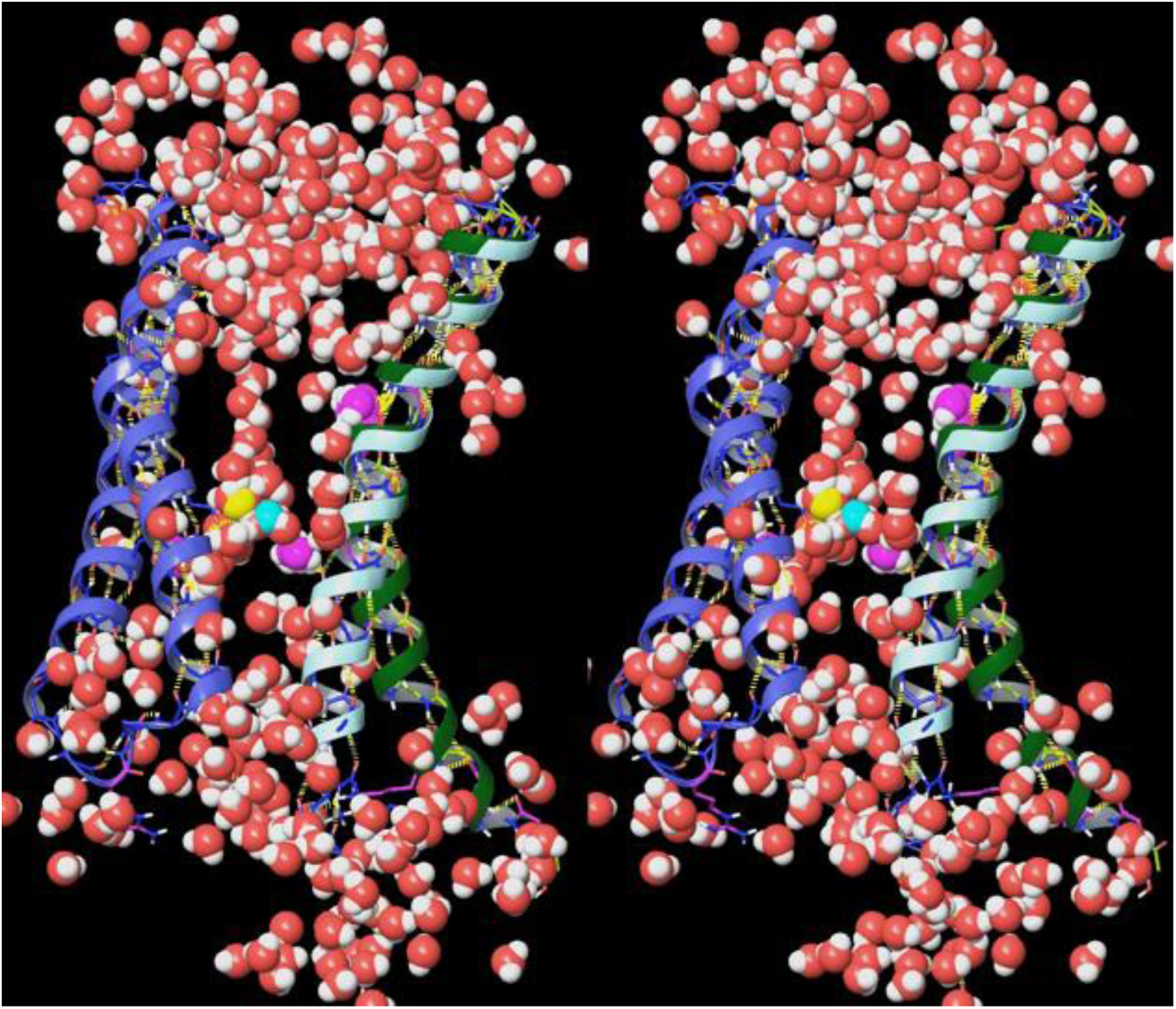
Stereo view of the external and internal solvation of several GPCR structures overlaid on the β_2_ adrenergic receptor (PDB code = 3SN6). The solvation content varies case-by-case among the individual receptor structures, but internal cavities detected using PyMol (see Materials and methods) are necessarily solvated by H-bond depleted water.

In our previous work, we studied the structure-function of cereblon and COVID M^pro^ guided by Biodynamics theory.^21,22^ Here, we apply our theory to the study of TMH rearrangements that convey activation and deactivation of GPCRs (exemplified by the β_2_ adrenergic receptor) and VGCCs (exemplified by Na_v_1.7), together with the means by which such rearrangements are powered by internal solvation and unilateral disruption of IHHBs within helical dyads that we refer to as “tropimers”.

## Materials and methods

We studied the canonical activation and deactivation mechanisms of class A GPCRs using the agonist- and antagonist-bound structures listed in Table 1 (ligands listed in Figure 2). Those of VGCCs using the structures listed in Table 2. All visualizations and calculations were performed using Maestro 2019-1 and PyMol 2.4.1 (Schrodinger, LLC, Portland, OR). The structures were prepared using the protein preparation tool in Maestro and overlaid using the protein structure alignment tool in PyMol. Sequence alignments were performed using the multiple sequence viewer in Maestro. Qualitative solvation field analysis of the open allosteric binding site of the β_2_-AR (3SN6) was performed using SiteMap (default settings). Topological maps of the β_2_-AR TMHs were generated by visual inspection. No structural modeling beyond that listed above was performed in this work.

**Figure 2.**
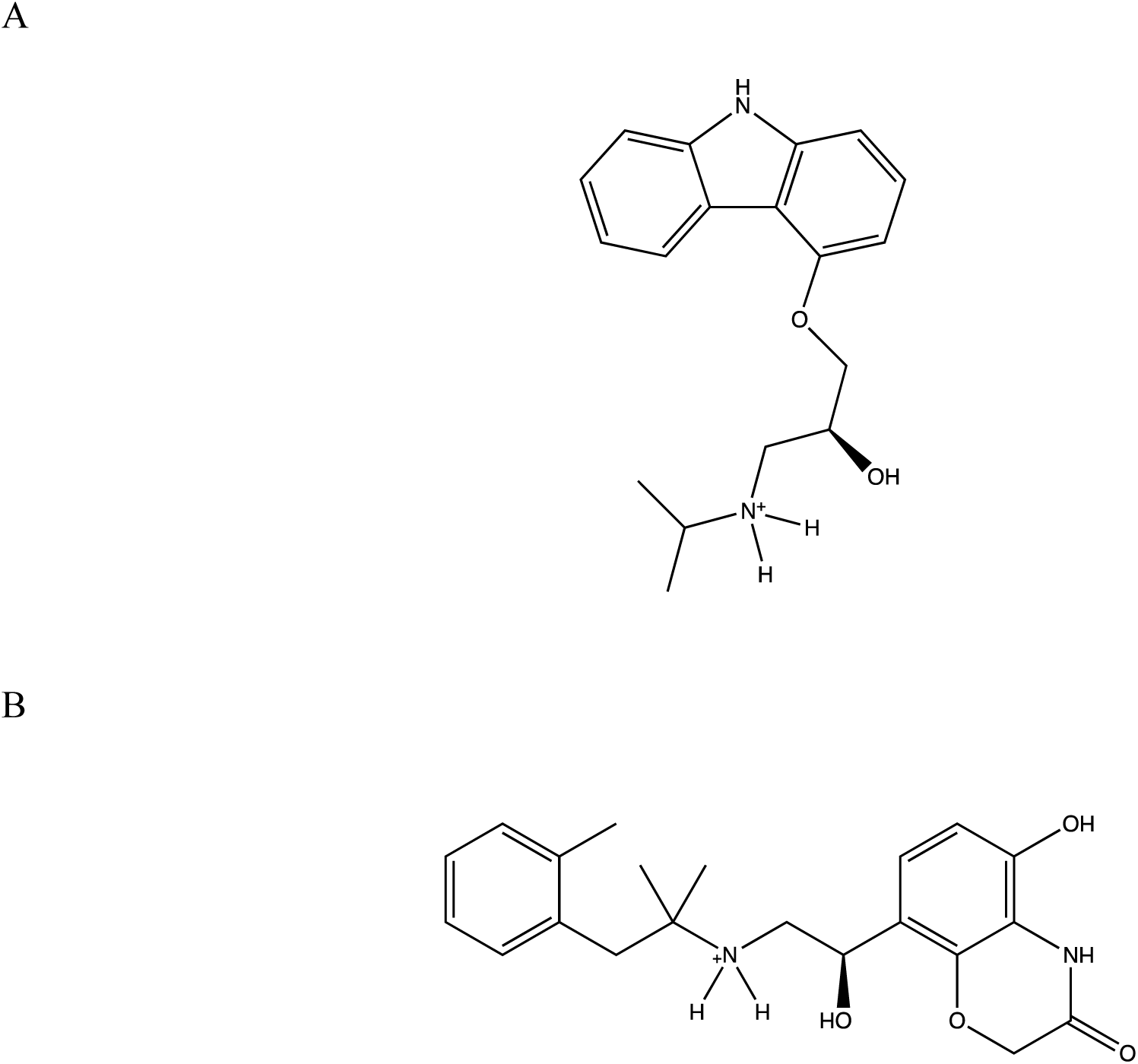
The b_2_-AR ligands studied in this work. (A) Antagonist (2S)-1-(9H-Carbazol-4-yloxy)- 3-(isopropylamino)propan-2-ol (CAU) (2RH1).^24^ (B) Agonist 8-[(1R)-2-{[1,1-dimethyl-2- (2methylphenyl)ethyl]amino}-1-hydroxyethyl]-5-hydroxy-2H-1,4-benzoxazin-3(4H)-one (P0G) (3SN6).^25^

**Table 1.** The set of GPCR receptors studied in this work.

| PDB code | State | GPCR family |
| --- | --- | --- |
| 3VW7 | Deactivated | Protease-activated receptor 1<sup>23</sup> |
| 2RH1 | Deactivated | $\beta_2$-adrenergic receptor<sup>24</sup> |
| 3SN6 | Activated | $\beta_2$-adrenergic receptor<sup>25</sup> |
| 6MXT | Intermediate | $\beta_2$-adrenergic receptor<sup>26</sup> |

**Table 2.** The set of VGCCs studied in this work.

| PDB code | Ion channel family |
| --- | --- |
| 6N4R | Cryo-EM structure of deactivated human Na<sub>v</sub>1.7<sup>18</sup> |
| 6N4Q | Cryo-EM structure of activated human Na<sub>v</sub>1.7<sup>18</sup> |
| 2A79 | Crystal structure of activated rat K<sub>v</sub>1.2<sup>27</sup> |
| 5VMS | Cryo-EM structure of deactivated KCNQ1<sup>28</sup> |

At the time of this work, the β_2_-AR is the only structurally known GPCR in both the native deactivated and fully activated agonist/G-protein-bound states and Na_v_1.7 is the only structurally known VGCC in both the activated and deactivated states (although solved in the presence of a toxin). We make the following assumptions:

1. Despite the presence of the toxin, the Na_v_1.7 structures capture the approximate native states of the channel.
2. The activation/deactivation mechanisms deduced from the β_2_-AR and Na_v_1.7 structures are paradigmatically similar across all class A GPCRs and VGCCs, respectively.
3. The deactivated state of GPCRs is similar to the antagonist-bound state. The orthosteric pocket is fully blocked in 3SN6 and partially blocked in 2RH1, which necessarily differs from the empty deactivated pocket.
4. The Na^+^ coordination site captured in the crystal structure of PAR1 is present in the deactivated state of all class A GPCRs, including the β_2_-AR receptor.

## Results

### Putative GPCR structure-mechanism relationships

In this work, we set about to study the following questions:

1. The interplay between the structural integrity vis-à-vis rearrangeability of the 7-TMH bundle in GPCRs based on the aforementioned crystal structures, spatial adjacency among the TMHs (Table 3), and qualitative topological maps of the deactivated (Figure 3A) and activated (Figure 3B) receptor states.
2. The putative mechanisms by which the TMHs rearrange during receptor activation and deactivation, including the means by which they are powered and governed by agonist association and dissociation.
3. The putative structure-kinetics relationships of GPCR-agonist, antagonist, and G-protein binding vis-à-vis our proposed activation/deactivation mechanism.\

**Figure 3.**
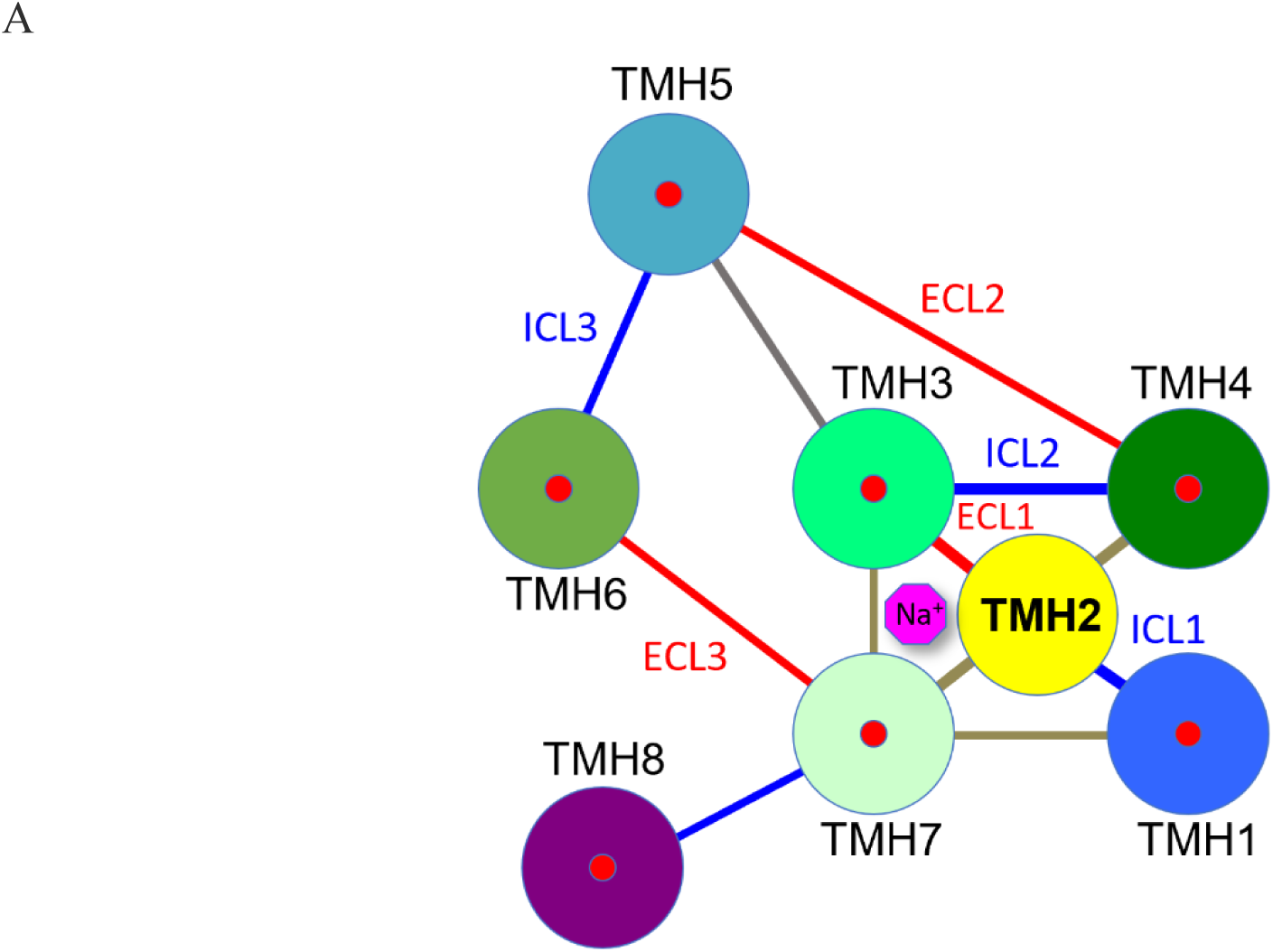
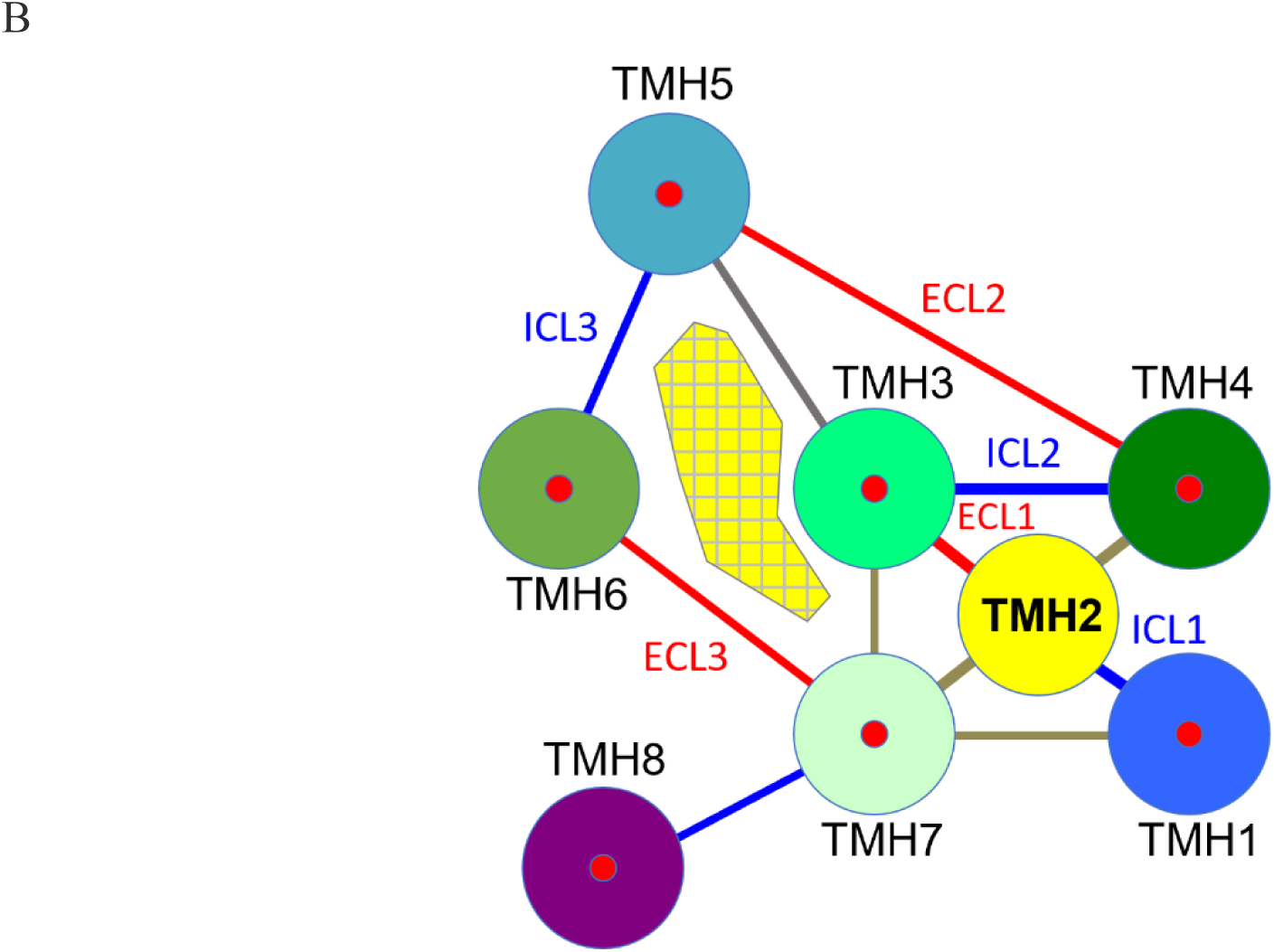
(A) Topology map of the deactivated and (B) activated 7-TMH bundle of the b_2_-AR based on the observed spatial adjacencies of the TMHs in 3SN6 and 2RH1. The allosteric binding site is denoted by the yellow crosshatched region between TMH3, TMH5, TMH6, and TMH7. The TMHs are denoted by filled circles, the ICLs and ECLs are labeled in blue and red, respectively, and the edges of disconnected TMHs are shown in gray. The 7-TM bundle can be partitioned into four components consisting of TMH1, TMH2-TMH3, TMH4-TMH5, and TMH6-TMH7 based on the ICL and ECL connectivity.

**Table 3.**
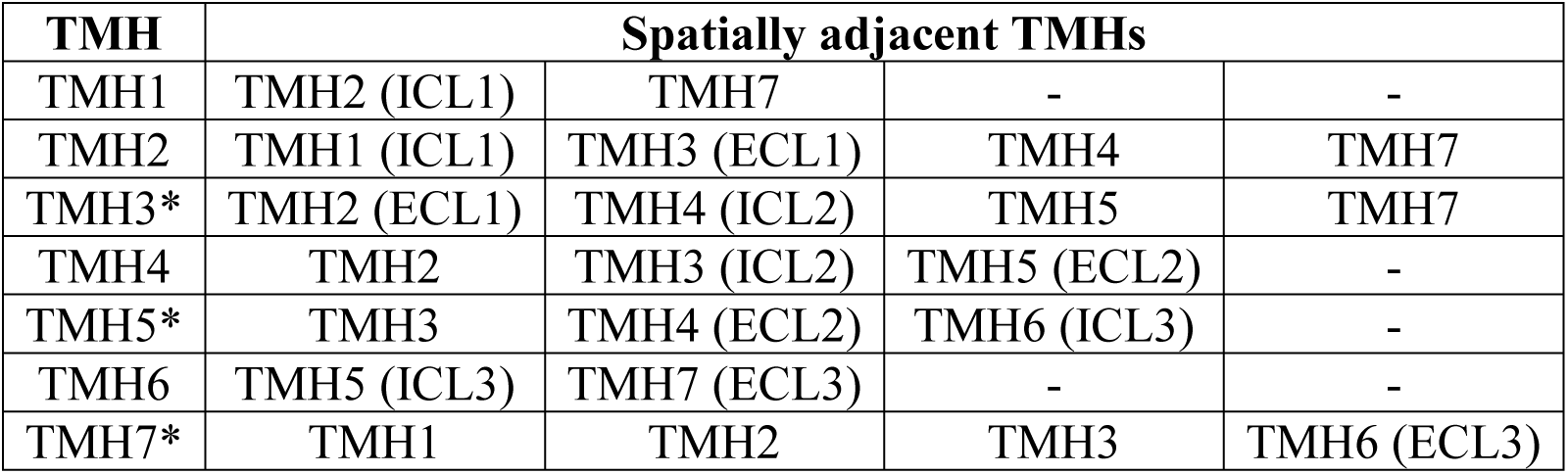
TMH adjacency matrix of the β_2_-AR. TMH3, TMH5, and TMH7 undergo first order rigid-body pivoting in direct response to agonist binding. TMH1, TMH2, TMH4, and TMH6 undergo second order pivoting and translation in response to the these rearrangements.

TMHs that form activating H-bonds with agonists are starred.

#### The putative role of the ECLs in receptor activation

We postulate that significant intra- and interhelical rearrangements occur during activation and deactivation of the β_2_-AR relative to a stationary anchor region, and we set about to deduce (but not simulate) the rearreangement pathway based on the correct best-fit overlay of the corresponding residues in the activated and deactivated states captured in 3SN6 and 2RH1, respectively (i.e., the two endpoints of the rearrangements). We attempted best-fit overlays between corresponding residues in the two structures, as follows:

1. About all such residues in the entire sequence (referred to as the “global overlay”) (Figure 4).
2. About the residues in each individual TMH and those in subsets of the TMHs (referred to as “local TMH overlays”).
3. About residues in ECL2, the backbone conformations of which are nearly identical in the two structures.

**Figure 4.**
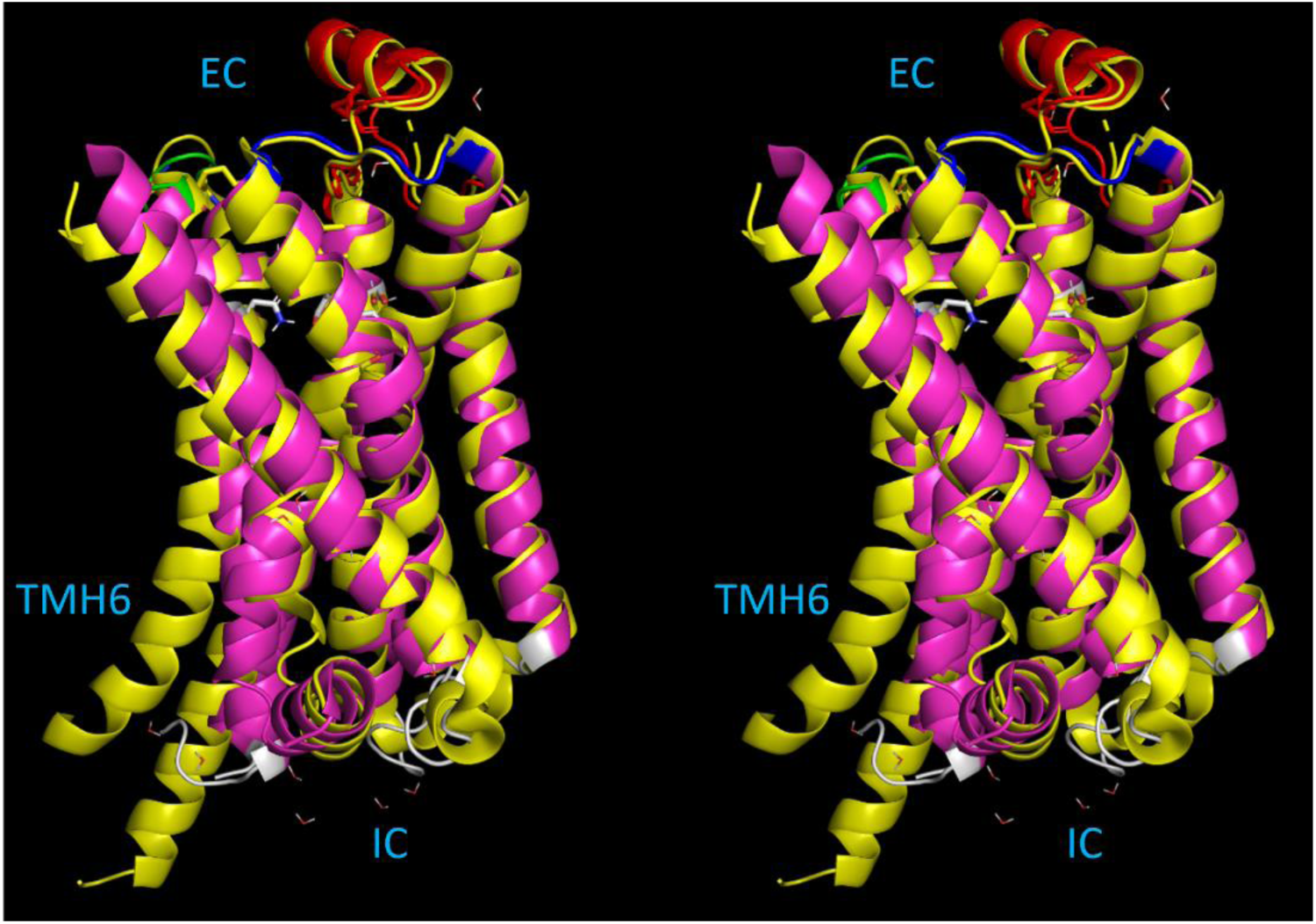
Stereo view of the <u>global</u> structural alignment of the activated and deactivated states of the β_2_-AR captured respectively in 3SN6 (magenta) and 2RH1 (yellow). The structures are oriented with the intracellular (IC) side at the bottom of the figure and the extracellular (EC) side at the top of the figure, in which ECL1 (blue), ECL2 (red), and ECL3 (green) are visible. TMH6 is labeled for reference. Structural differences reside mainly in the TMH6 and the ICL conformations. However, activation/deactivation relevant differences are averaged out in this alignment.

We assessed each of these overlays based on their implied intra- and interhelical rearrangement pathways. The global overlay is limited largely to intra-helical rearrangements in TMH6 and TMH7 (suggesting that the receptor converges to similar inter-helical relationships among the two states), whereas the local TMH overlays are inconsistent with meaningful inter-helical rearrangements. On the other hand, the ECL2 overlay is consistent with meaningful inter- and intra-helical rearrangements of all TMHs, suggesting that ECL2 serves as the stationary anchor region of the bundle (i.e., that the TMHs move in relation to this loop). This hypothesis is further supported by the following observations:

1. ECL2 is semi-rigid (containing an α-helix and a disulfide bond) (Figure 5A).
2. ECL2 and ECL3 are well-packed in the activated and deactivated structures, suggesting that they behave as an integral unit (Figures 5B-C).
3. Highly meaningful correspondences exist between the TMH positions/orientations/conformations and internal solvation cavities (described at length below).
4. ECL2 is known to play a key role in receptor activation,^24,29^ destabilization of which leads to a 1,000-fold decrease in agonist activity.^30,31^

**Figure 5.**
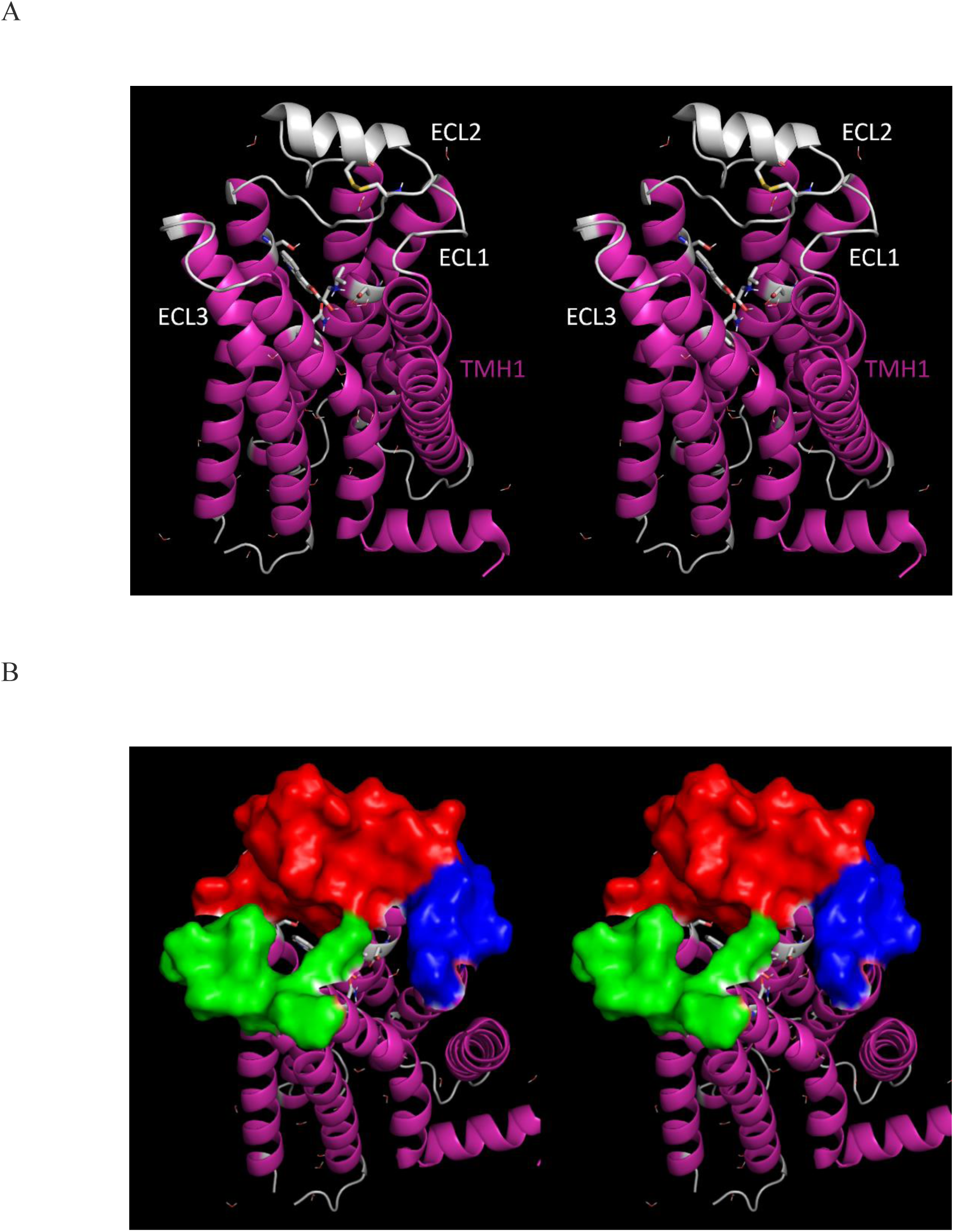
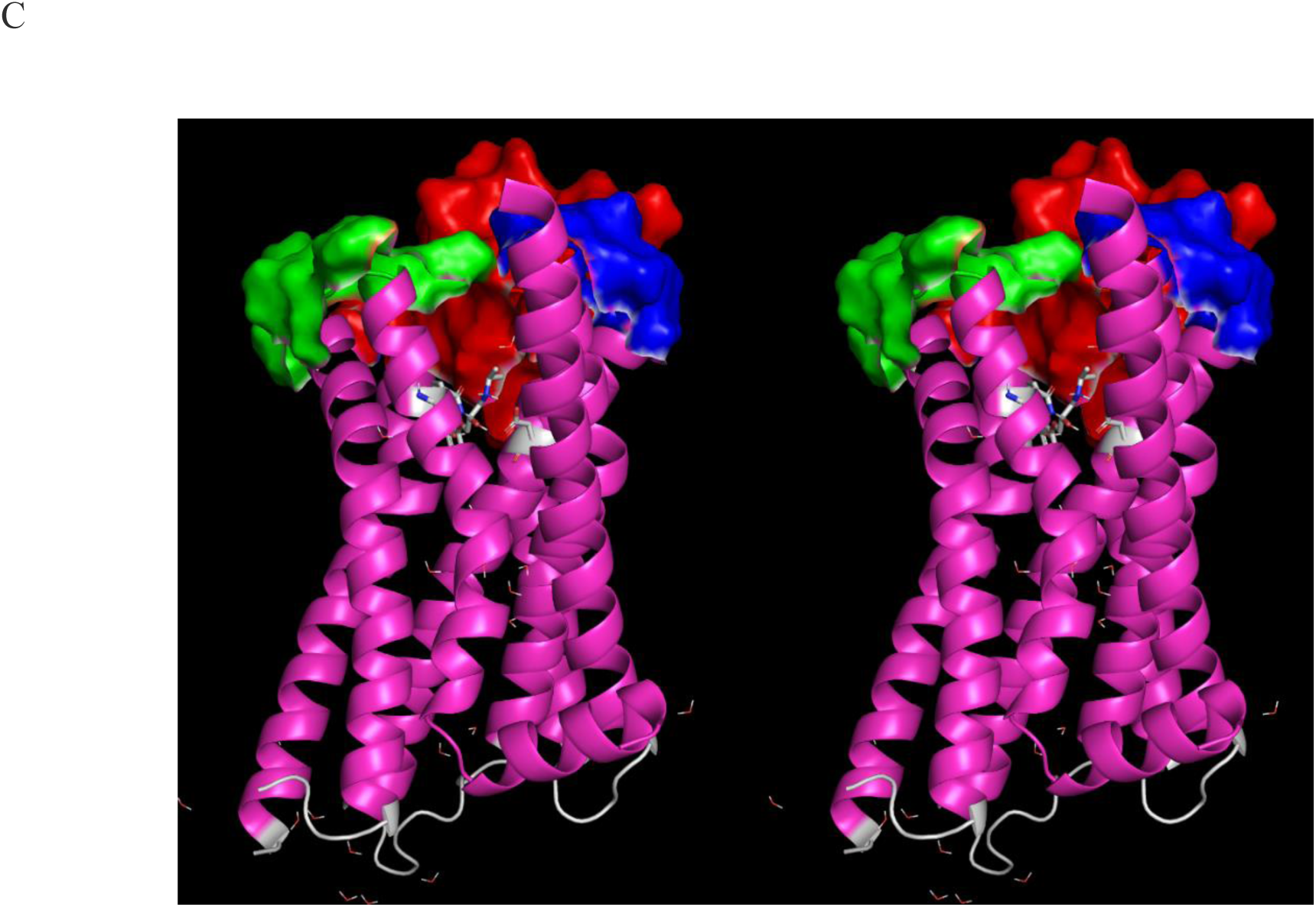
Stereo views of the deactivated β_2_-AR (2RH1). (A) The ECL loop configuration (white regions at the top of the figure) in the deactivated state viewed in the approximate EC → IC direction. (B) Surfaces of the three ECLs shown in A. ECL1 (blue) and ECL2 (red) pack into an integral substructure that putatively anchors TMH2, TMH3, TMH4, and TMH5 (analogous to the cross-brace of a marionette). ECL3 (green) anchors TMH6 and TMH7 (noting that TMH1 is unanchored by an ECL). (C) Same as B, except viewed approximately perpendicular to the 7-TMH bundle axis. ECL2 projects deep into the bundle (red surface), packing with both the antagonist and agonist in 2RH1 (white) and 3SN6 (not shown), which are transiently trapped under the loop in the activated state.

#### Activating TMH rearrangements are putatively jumpstarted by agonist binding

We postulate that receptors undergo <u>non-equilibrium</u> binding with agonists and antagonists due to transient agonist/antagonist buildup and decay, receptor inactivation by beta-arrestins, receptor internalization and degradation, and other mechanisms. As such, agonist and antagonist binding is best described in terms of dynamic occupancy (governed by association and dissociation barriers and rate constants), rather than affinity per se, and the effects thereof on receptor activation and deactivation are best described in terms of barriers that are modulated by agonist and antagonist occupancy. We further postulate that:

1. The degree of pocket opening is not all-or-none (noting that the fully open form is unobserved experimentally, the partially open form is observed in 2RH1 (Figure 6A), and the closed form in 3SN6 (Figure 6B)). The closed form, in which agonists are transiently trapped, is driven by agonist-induced TMH rearrangements that are described in the following section.
2. Agonist and antagonist association barriers 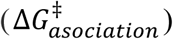 and the corresponding k_on_s depend on the degree of mutual replacement of the H-bonds of H-bond enriched binding partner solvation (i.e., the desolvation costs of the ligand and pocket), which is typically a zero sum game at best (where fully optimal replacement results in no net desolvation loss or gain, translating to diffusion-lmited k_on_). However, the high surface curvature and high non-polar content of the orthosteric pocket in the β_2_-AR suggest that the polar groups of the receptor participating in agonist H-bonding, (Asp113/TMH3, Ser203 and Ser207/TMH5, and Asn312/TMH7^32^ (Figure 7)), are solvated by <u>atypically higher free energy</u> (unfavorable) H-bond enriched water, such that replacement of the H-bonds thereof by polar agonist H-bond groups results in the net <u>gain</u> of desolvation free energy (i.e., ΔG_net_ = ΔG_desolvation_ − ΔG_H-bond_ < 0) needed to surmount the barrier to TMH3, TMH5, and TMH7 rearrangements (i.e., ΔG_TM3-TM5-TM7 rearrangements_ ≈ ΔG_net_) (noting that this barrier is necessarily lower in constitutively active receptors).
3. The agonist dissociation barrier 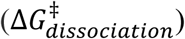 and corresponding k_off_s depend principally on the resolvation costs of the dissociating binding partners, equating to the total free energy of the H-bond <u>depleted</u> solvation expelled during association. The agonist k_off_ in the pre-trapped state is necessarily optimized to the rate of pocket closure and trapping, and is necessarily zero in the trapped state. Although we did not calculate the solvation field within the orthosteric pocket, H-bond depleted solvation underlying 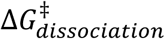 is typically present on non-polar molecular surfaces that are either highly concave or convex^22^ (noting that binding is enhanced by aromatic ligand groups positioned between Val114 and Phe290^32–34)^.
4. GPCR operation depends on the delicate balance between the following free energy contributions:

a. <u>Internal</u> desolvation and resolvation costs governing the rates of activating and deactivating TMH rearrangements.
b. <u>External</u> desolvation and resolvation costs of the agonist and orthosteric pockets governing agonist k_on_ and k_off_.
c. <u>External</u> desolvation and resolvation costs of the G-protein and allosteric pocket governing G-protein k_on_ and k_off_.
d. IHHB disruption costs governing the rates of helical curvature (described below).
5. Receptor activation is slowed/sped by the following:

a. Increased/decreased desolvation costs governing the rates of activating TMH rearrangements.
b. Increased/decreased desolvation costs of Asp113, Ser203, Ser207, and/or Asn312, resulting in diminished/increased net H-bond free energy gains and mistmatching between this energy and the receptor activation barrier (i.e., ΔG_TM3-TM5-TM7 rearrangements_ > |ΔG_net_|).
c. Decreased resolvation costs governing agonist k_off_, resulting in agonist dissociation prior to receptor activation and G-protein binding.
6. Receptor deactivation would be respectively slowed/sped by:

a. Decreased/increased resolvation cost governing the rates of deactivating TMH rearrangements.
b. Increased/decreased resolvation cost of receptor-G-protein dissociation governing G-protein k_off_ (noting that receptor deactivation depends on G-protein dissociation).

**Figure 6.**
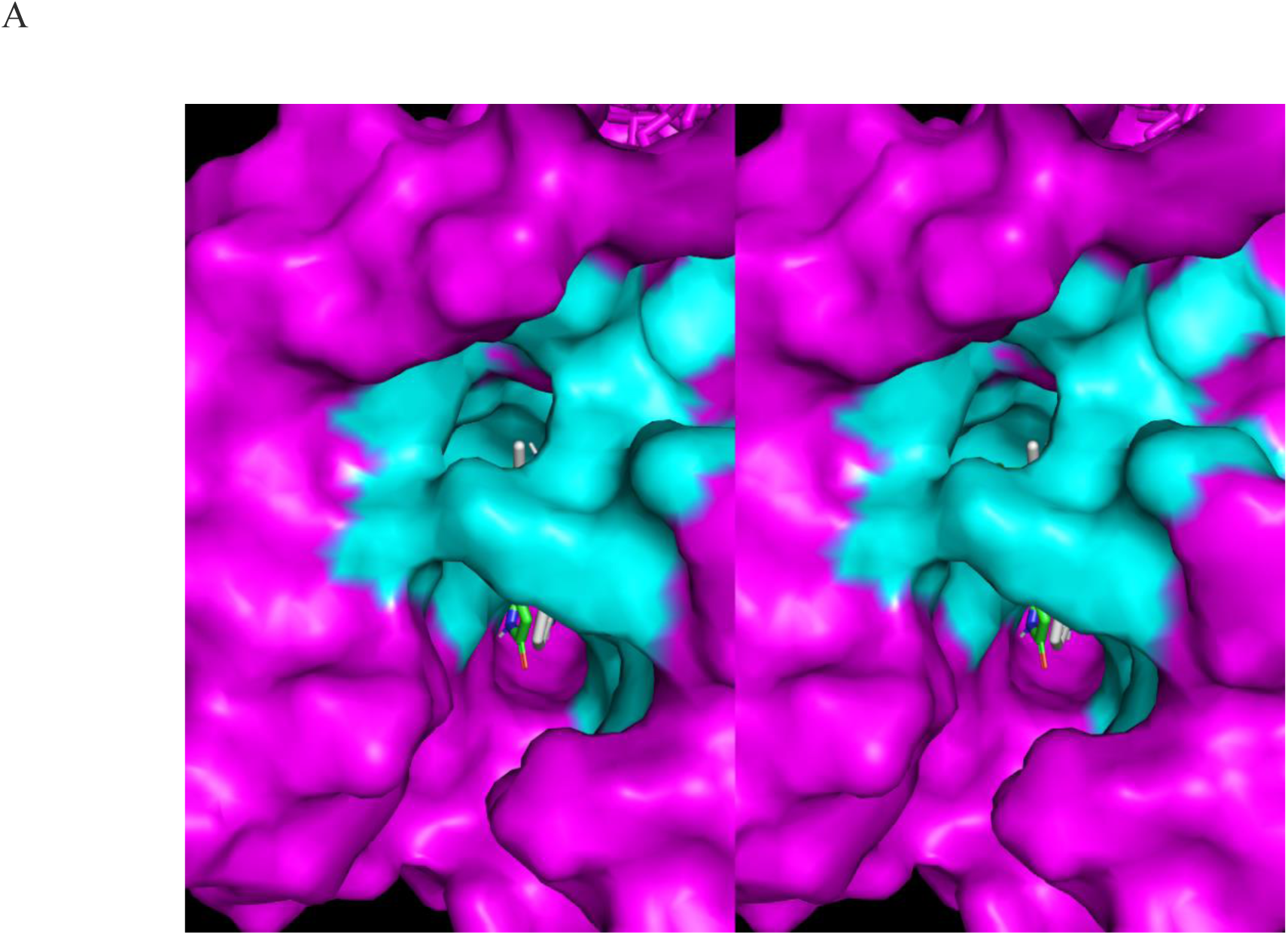
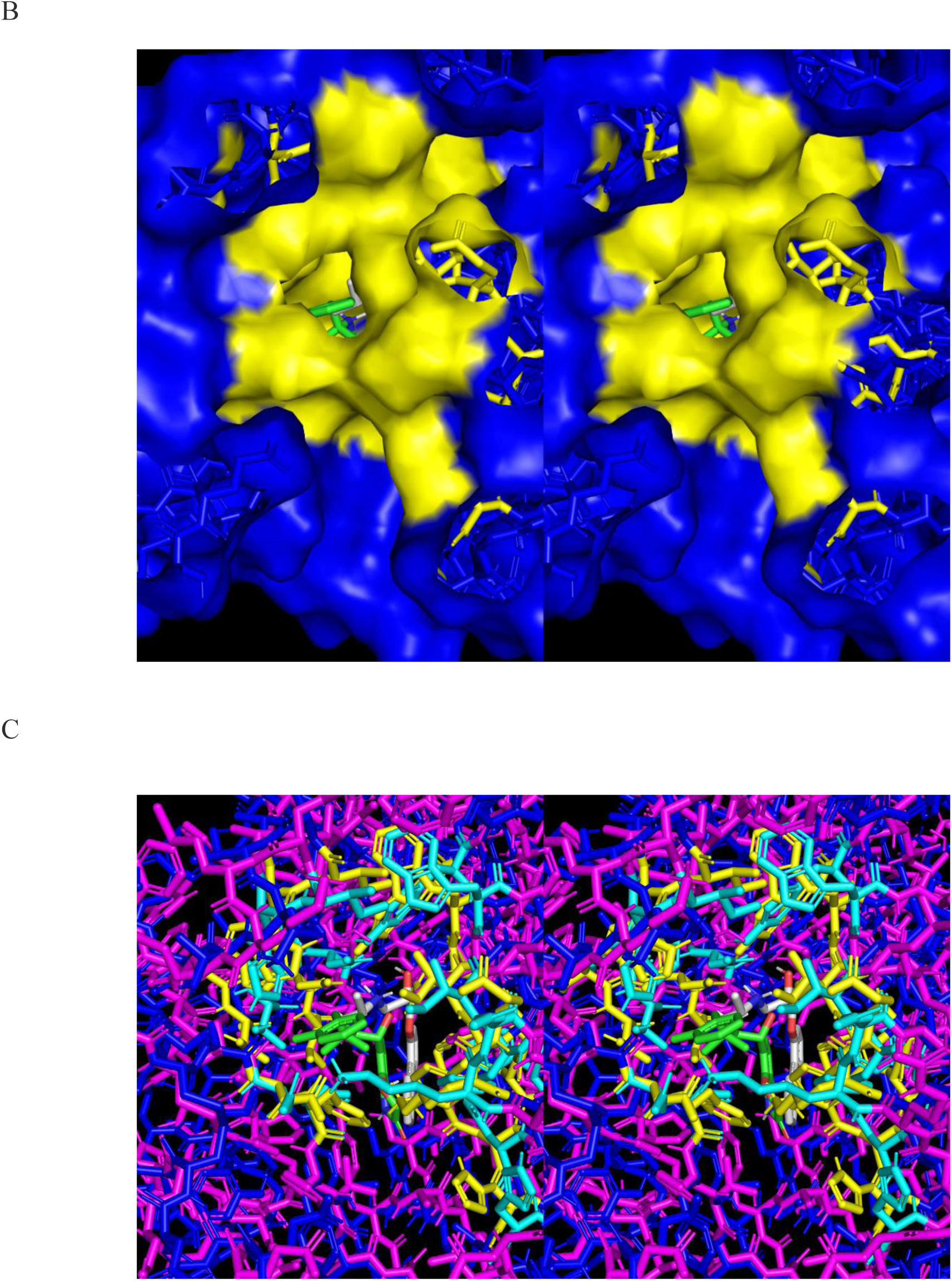
Stereo views of the agonist- (green sticks) and antagonist- (white sticks) bound orthosteric pocket in the β_2_-AR, looking in the EC to IC direction. (A) The deactivated state, in which the pocket is largely open for agonist and antagonist association. The degree of closure is determined by a specific set of side chains (yellow) the spatial positions of which are governed principally by the empty versus agonist/antagonist-induced positions/orientations of the TMHs on which they are located (noting that side chain conformational differences may additionally govern the degree of pocket opening). (B) Same as A, except for the activated agonist-bound state in which the agonist is transiently trapped. The side chains governing the degree of pocket opening are shown in cyan. (C) Overlay of the structures of the deactivated (magenta) and activated (blue) states (2RH1 and 3SN6, respectively), showing the main residues governing the open (cyan sticks) and closed (yellow sticks) state of the pocket.

**Figure 7.**
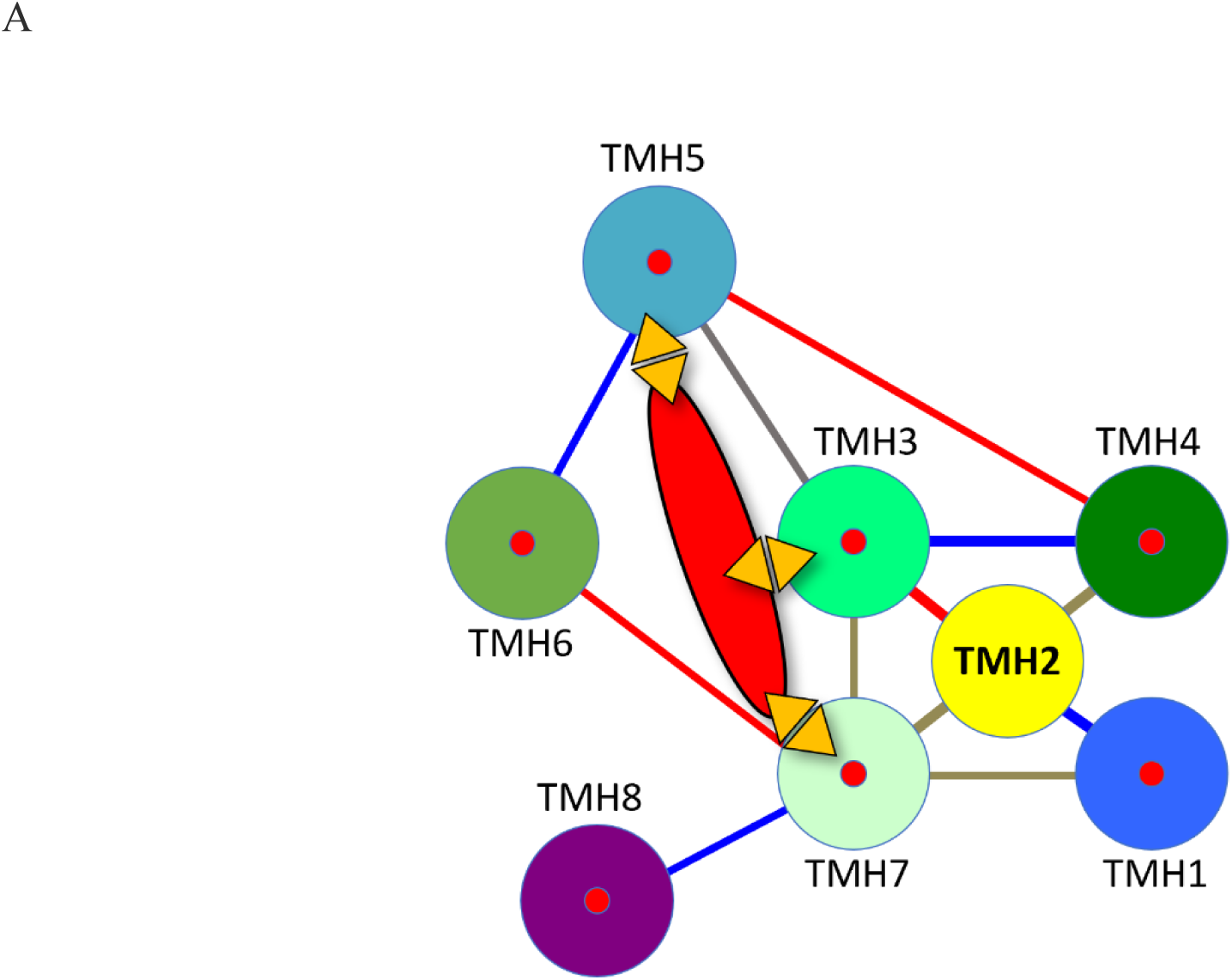
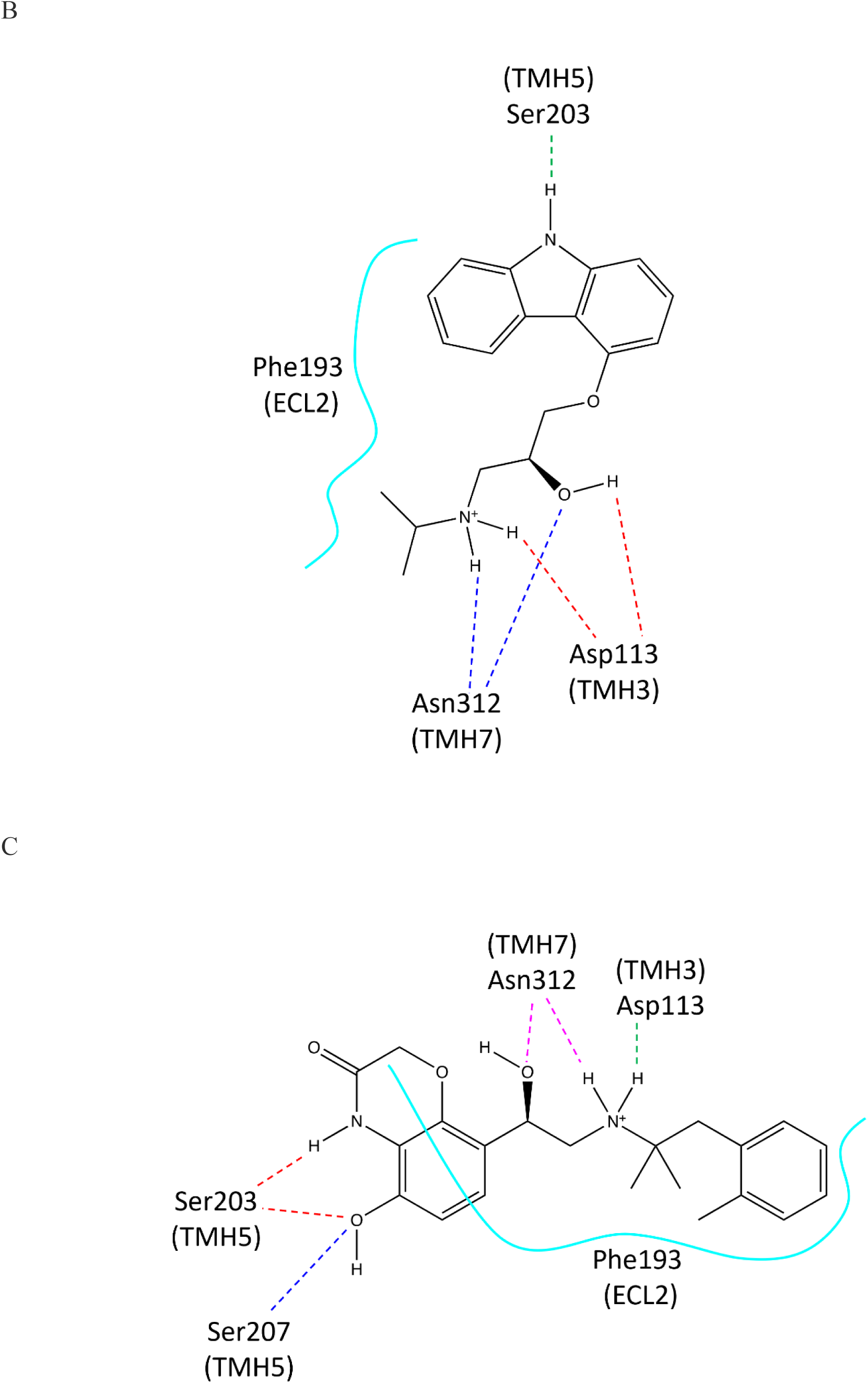
(A) The topological H-bond network (orange triangles) linking TMH3, TMH5, and TMH7 to the agonist (red oval). (B) Comparison of the antagonist and (C) agonist H-bond networks within the orthosteric pocket of the β_2_-AR.

#### Agonist-induced first and second order rigid-body pivoting of the TMHs relative to ECL2

Alignment of 3SN6 and 2RH1 on ECL2 reveals the postulated intra- and inter-helical TMH activation and deactivation pathways, consisting of rotation/pivoting and translational rearrangements. The TMH conformations are highly similar in all cases, except those containing switchable tropimers (Figure 8).

**Figure 8.**
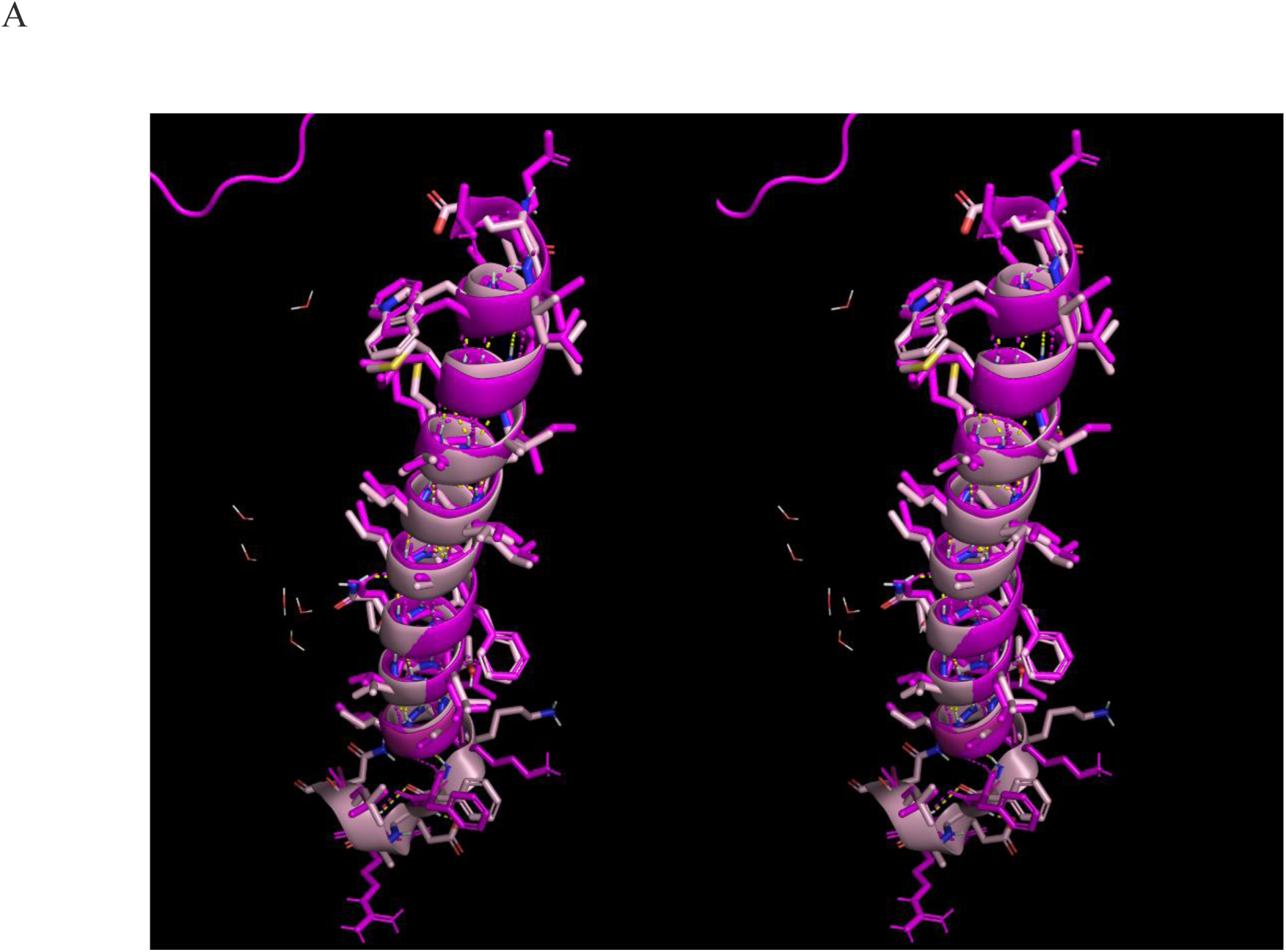
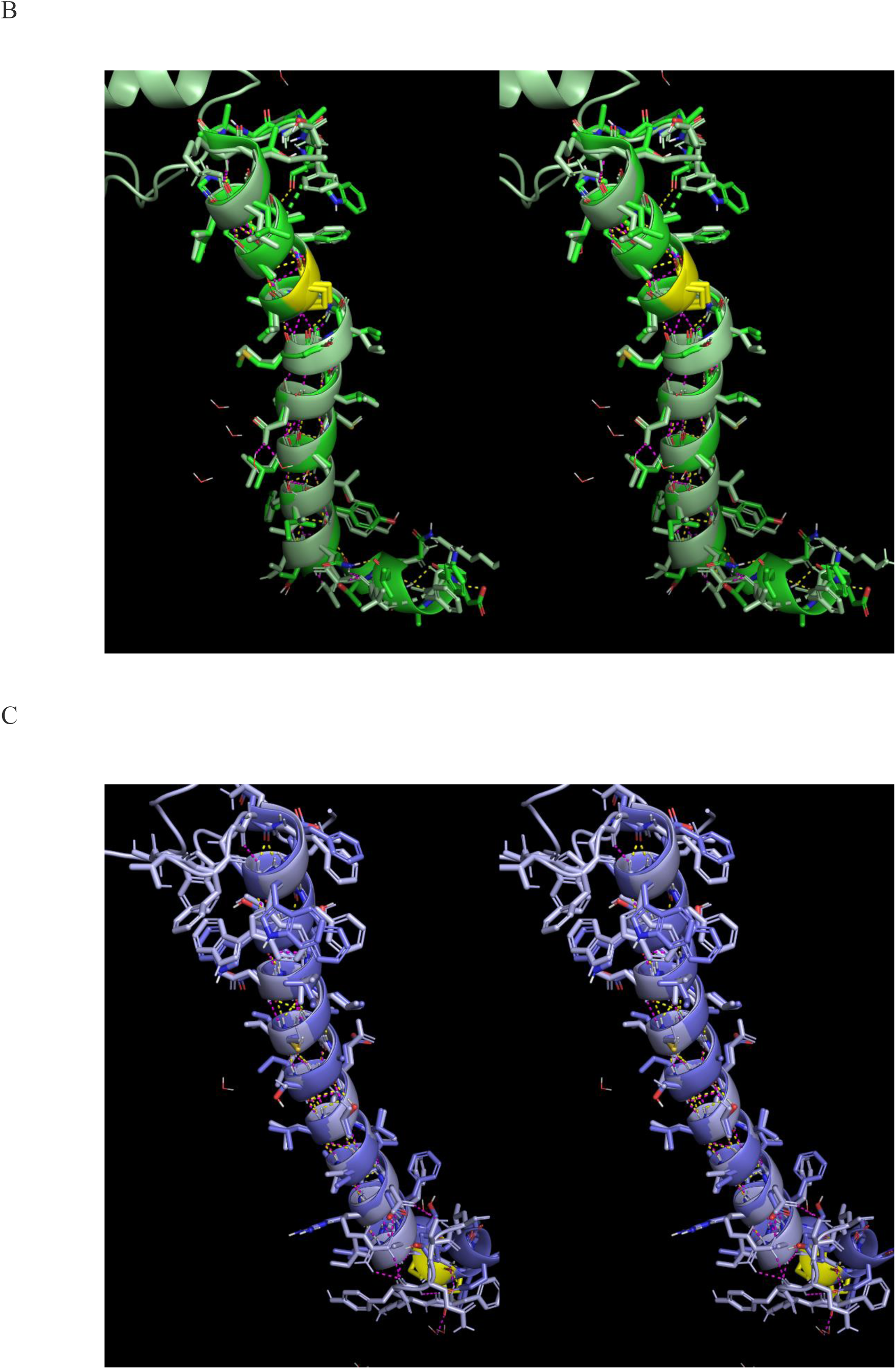
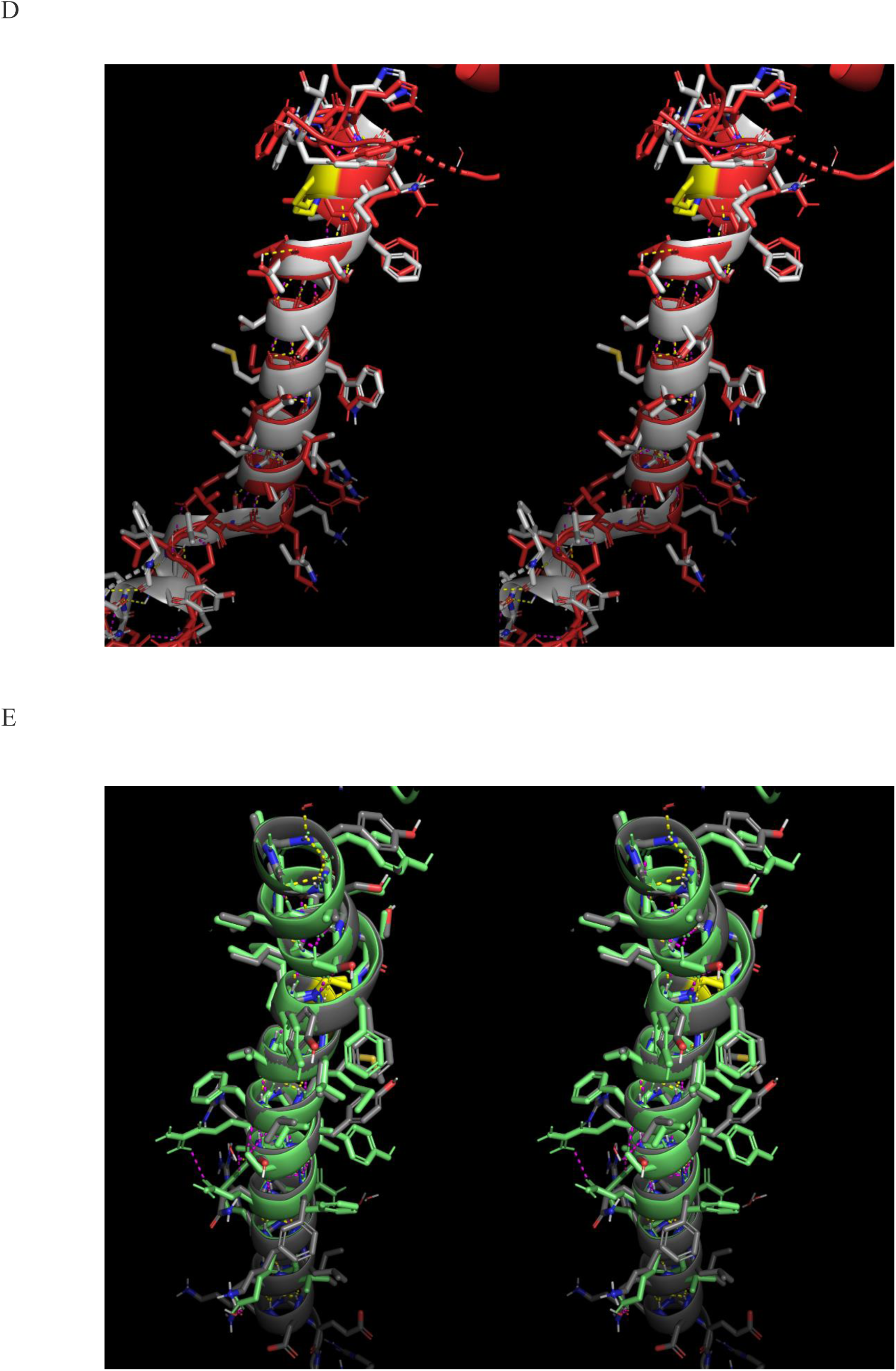
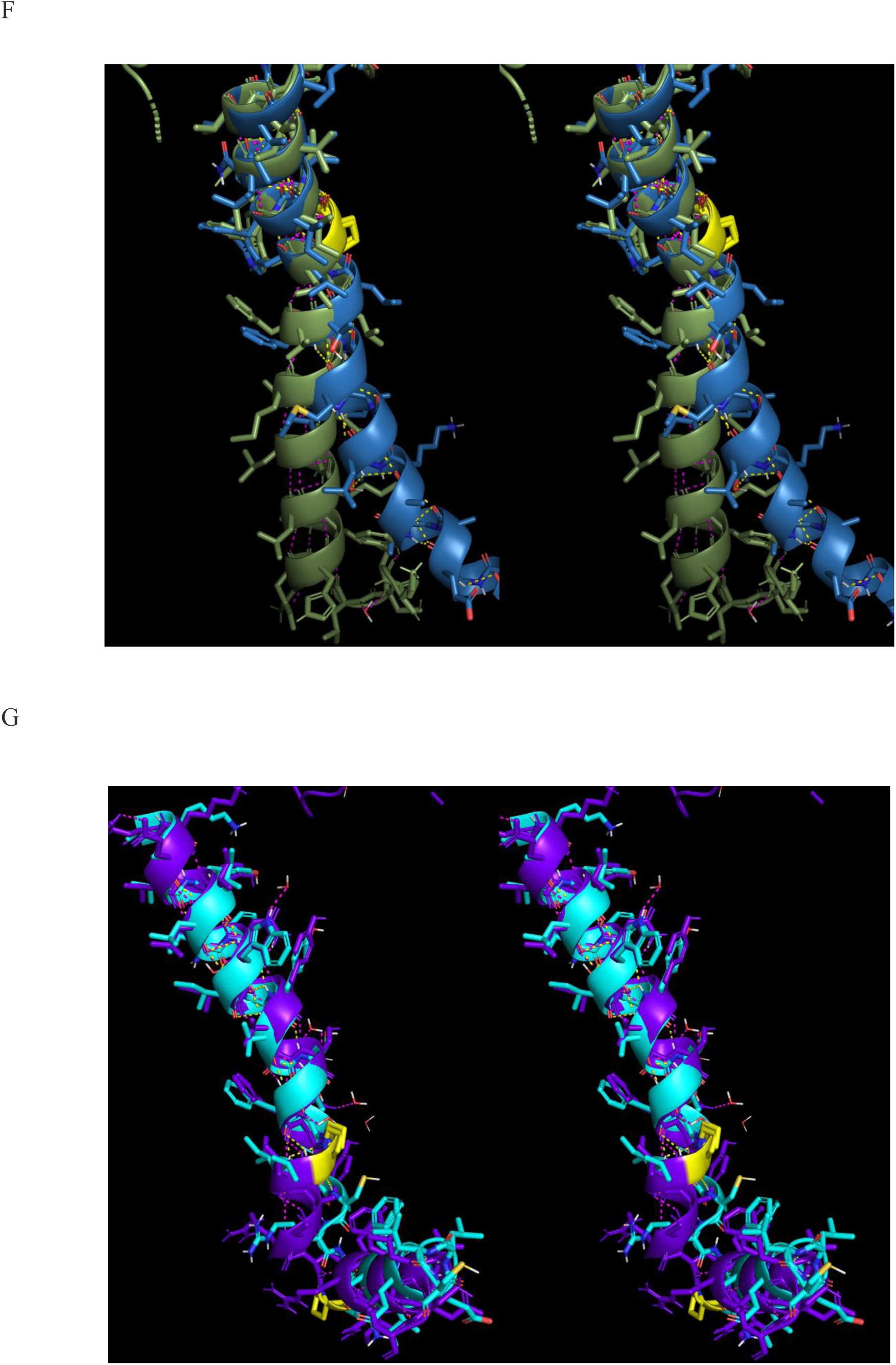
Stereo views of one-to-one overlays of each TMH of the activated (3SN6) and deactivated (2RH1) receptor structures. Proline positions are highlighted in yellow. The TMHs are nearly identical in both states, except for switchable tropimers in TMH6 and TMH7. (A) Activated (pink) versus deactivated (magenta) TMH1. (B) Activated (green) versus deactivated (light green) TMH2. (C) Activated (blue) versus deactivated (light blue) TMH3. (D) Activated (white) versus deactivated (red) TMH4. (E) Activated (gray) versus+ deactivated (light green) TMH5. (F) Activated (blue) versus deactivated (olive green) TMH6. (G) Activated (cyan) versus deactivated (purple) TMH7.

The TMHs undergo pivoting (i.e., scissors motion) on the EC side of the bundle, and as such sweep out small arcs on that side and larger arcs on the IC side (commonly referred to as “lever arm effects”) (Figure 9). Additionally, small translations are observed between the two states by which inter-helical packing is likely optimized.

**Figure 9.**
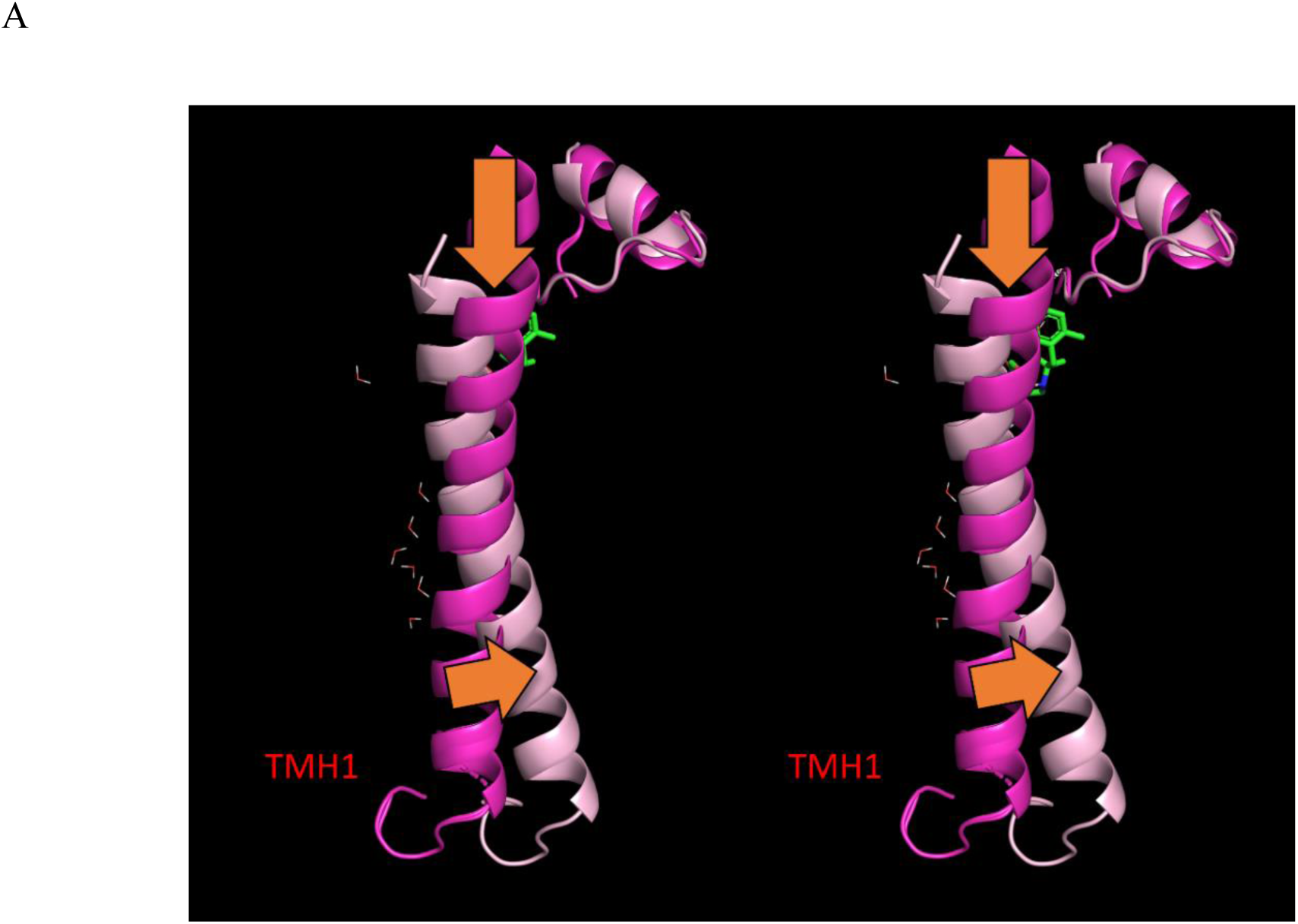
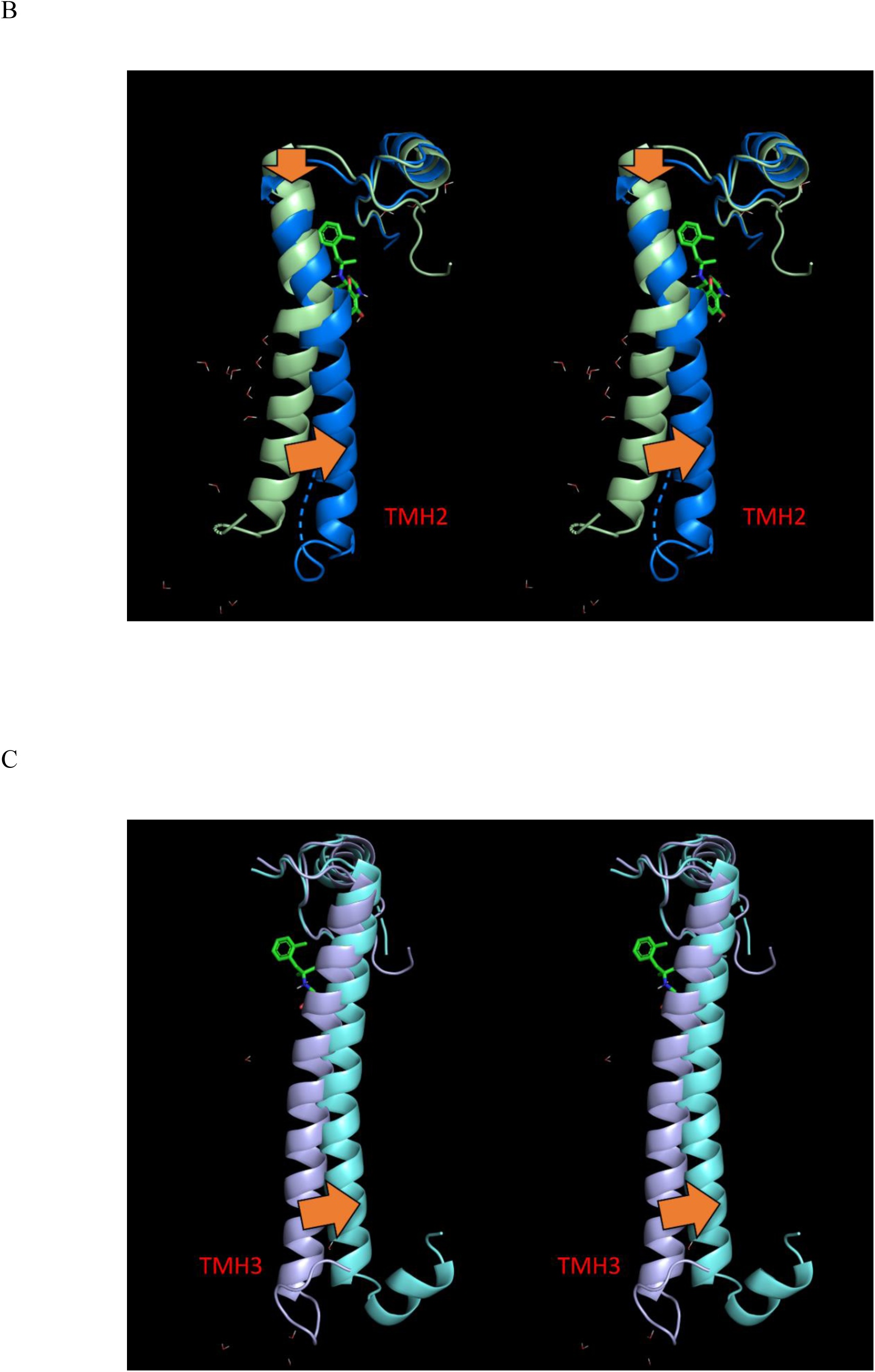
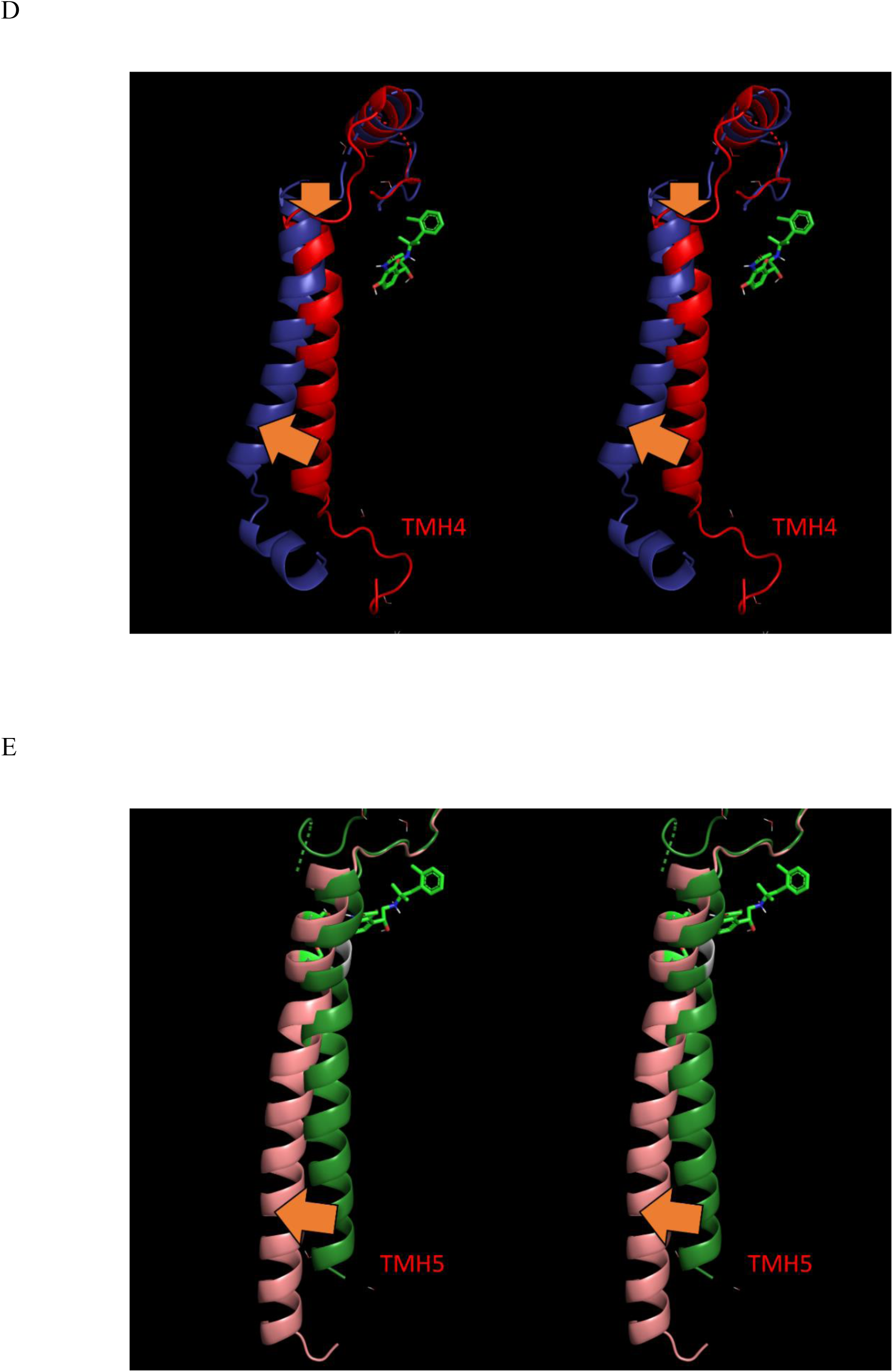
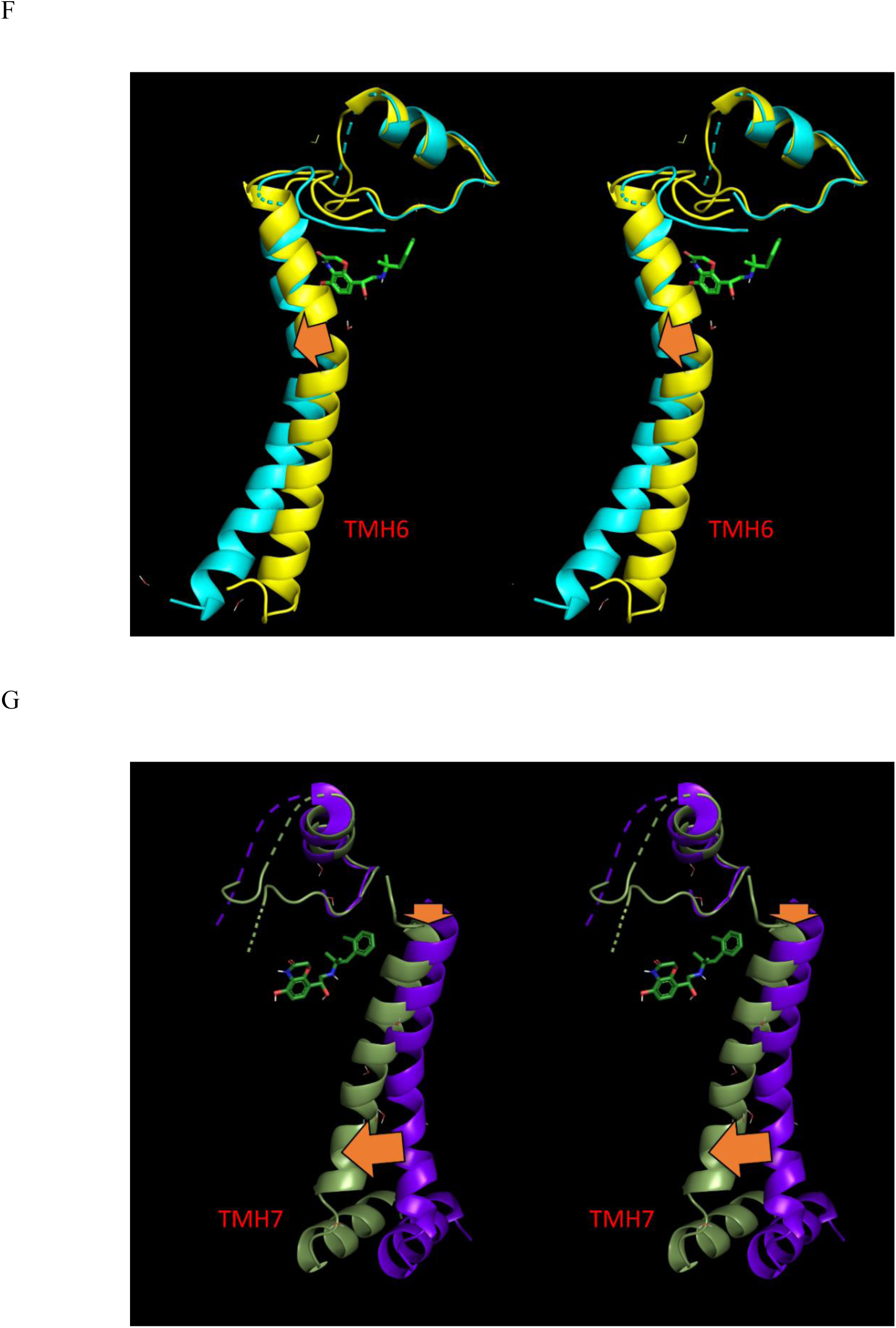
Stereo views of the activated receptor (3SN6) aligned about ECL2 in the deactivated state (2RH1), showing agonist-mediated first order rigid-body pivoting of TMH3, TMH5, and TMH7 and second order rearrangements of the adjacent TMHs. The rearrangement directions are annotated with orange arrows. (A) TMH1 deactivated (magenta) → activated (pink). (B) TMH2 deactivated (light green) → activated (blue). (C) TMH3 deactivated (violet) → activated (teal). (D) TMH4 deactivated (red) → activated (dark blue). (E) TMH5 deactivated (dark green) → activated (salmon). (F) TMH6 deactivated (yellow) → activated (cyan). (E) TMH7 deactivated (purple) → activated (olive green).

<u>Agonists</u> bind to the deactivated state. However, complementarity between agonist and TMH3, TMH5, and TMH7 partner positions exists solely in the <u>activated</u> state (Figure 10). Optimal complementarity between the H-bond partners of TMH3, TMH5, and TMH7 and <u>agonist</u> partners is achieved via the rigid-body pivoting and translation of those TMHs described above (Figure 11). Conversely, optimal complementarity between <u>antagonist</u> and H-bond partner positions within the 7-TMH bundle exists soley in the deactivated state, and as such, the TMH positions remain more or less stationary. The rotational orientations among the TMHs in both states are consistent with the classical ridge-groove α-helix packing arrangement.^35^

**Figure 10.**
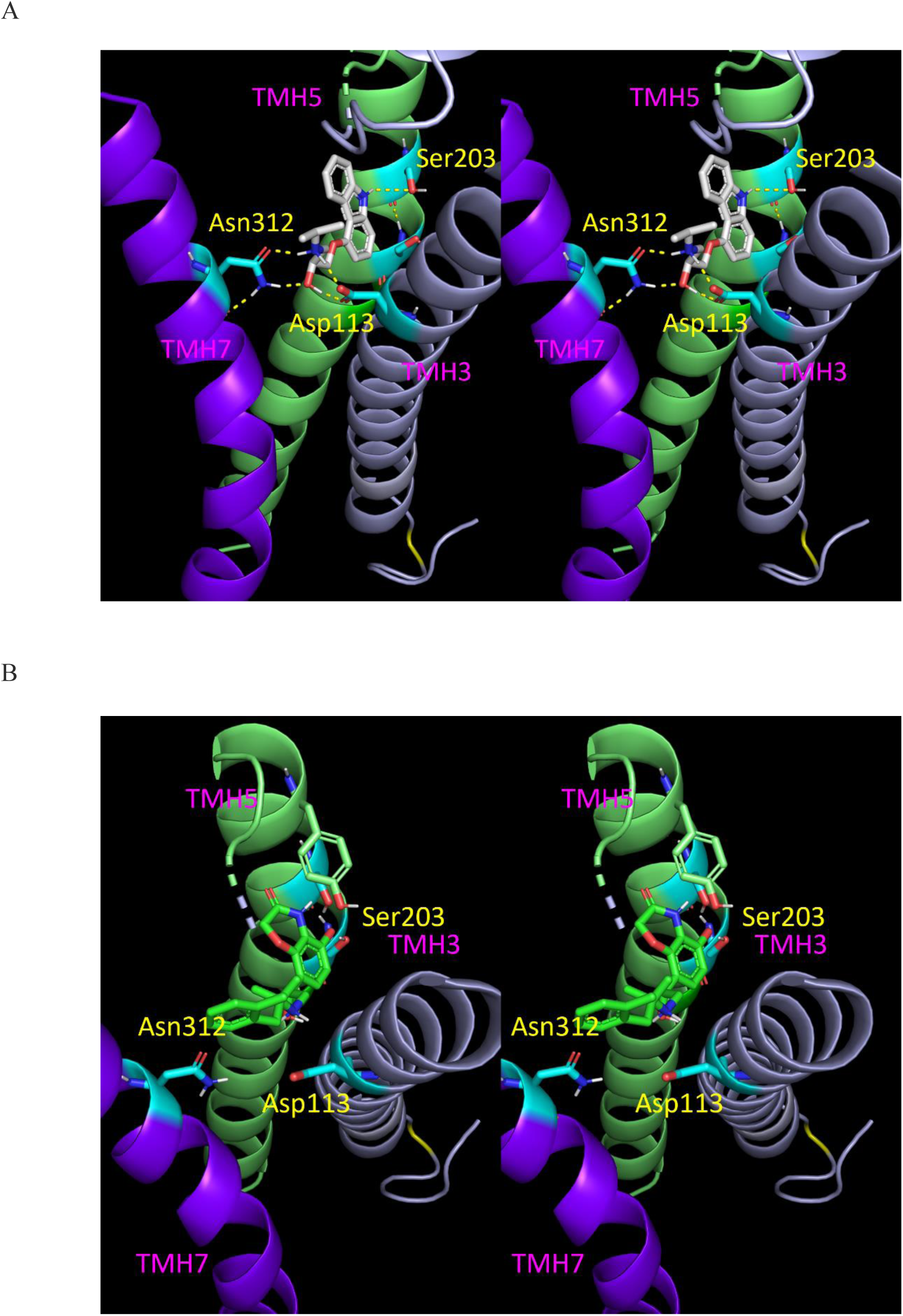
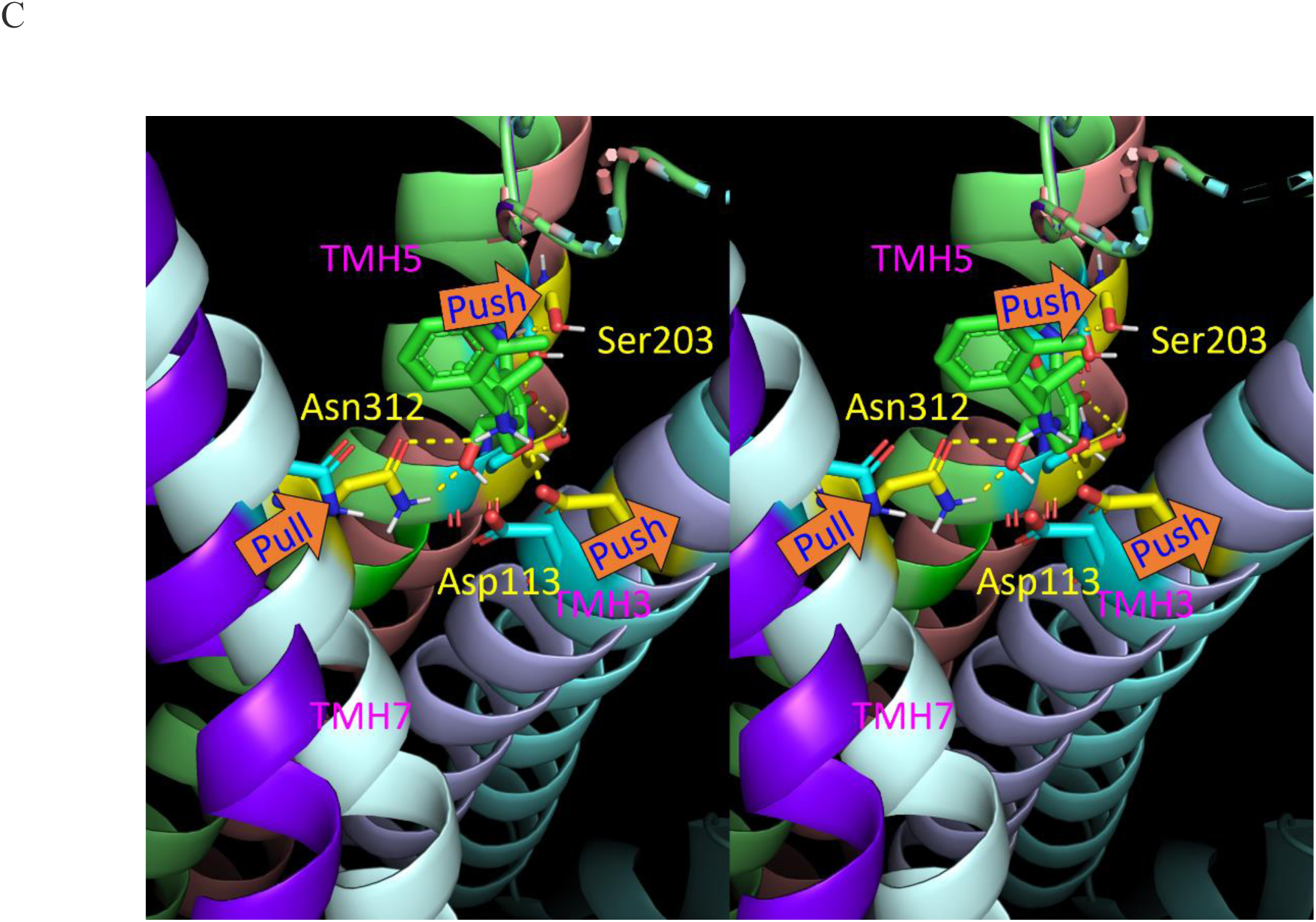
(A) Stereo view of the <u>antagonist</u> H-bond network in 2RH1 (H-bonds shown as yellow dotted lines). The antagonist is well-matched to the H-bond partner positions in the deactivated non-pivoted configuration of TMH3, TMH5, and TMH7. (B) Same as A, except for the agonist, which is mismatched to the deactivated TMHs. (C) Same as A, except for the agonist H-bond network in 3SN6 overlaid on the equivalent TMHs in 2RH1. Agonist-receptor H-bonds depend on pivoting of TMH3, TMH5, and TMH7 in the directions shown by the solid orange arrows. Pivoting is constrained to the free volumes (corresponding to internal solvation, as described below) within the bundle, together with the optimal H-bond geometries.

**Figure 11.**
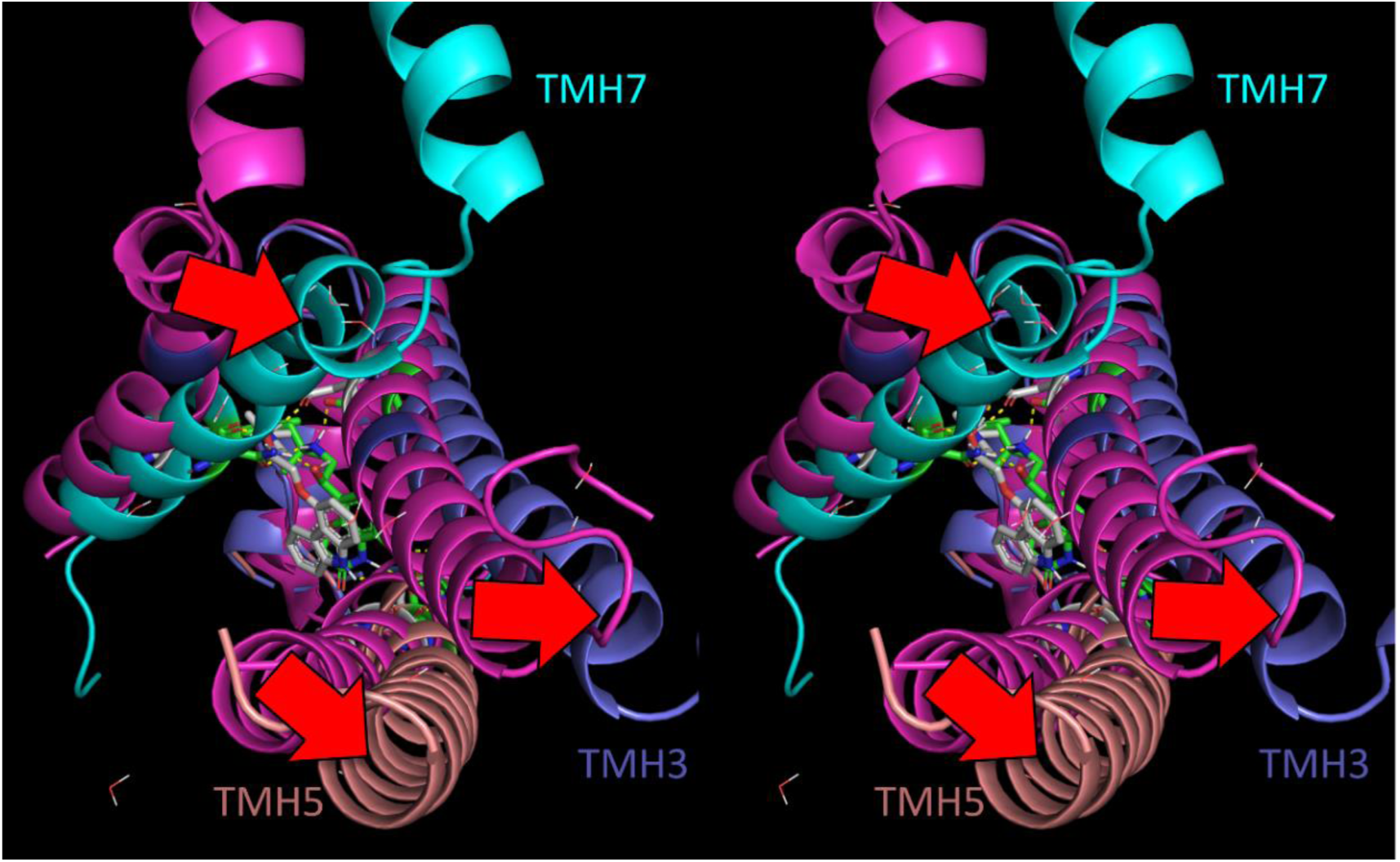
Stereo view of the deactivated (magenta) and activated (blue) states of TMH3, TMH5, and TMH7 of the β_2_-AR (2RH1 and 3SN6, respectively) overlaid on ECL2. These TMHs undergo rigid-body pivoting in direct response to H-bond formation with agonists, but not antagonists.

Opening of the allosteric pocket on the IC side of the 7-TMH bundle is achieved largely via outward splaying of the TMHs relative to the centroid of the bundle, accompanied by ICL adjustments needed to span the altered inter-TMH distances (Figure 12).

**Figure 12.**
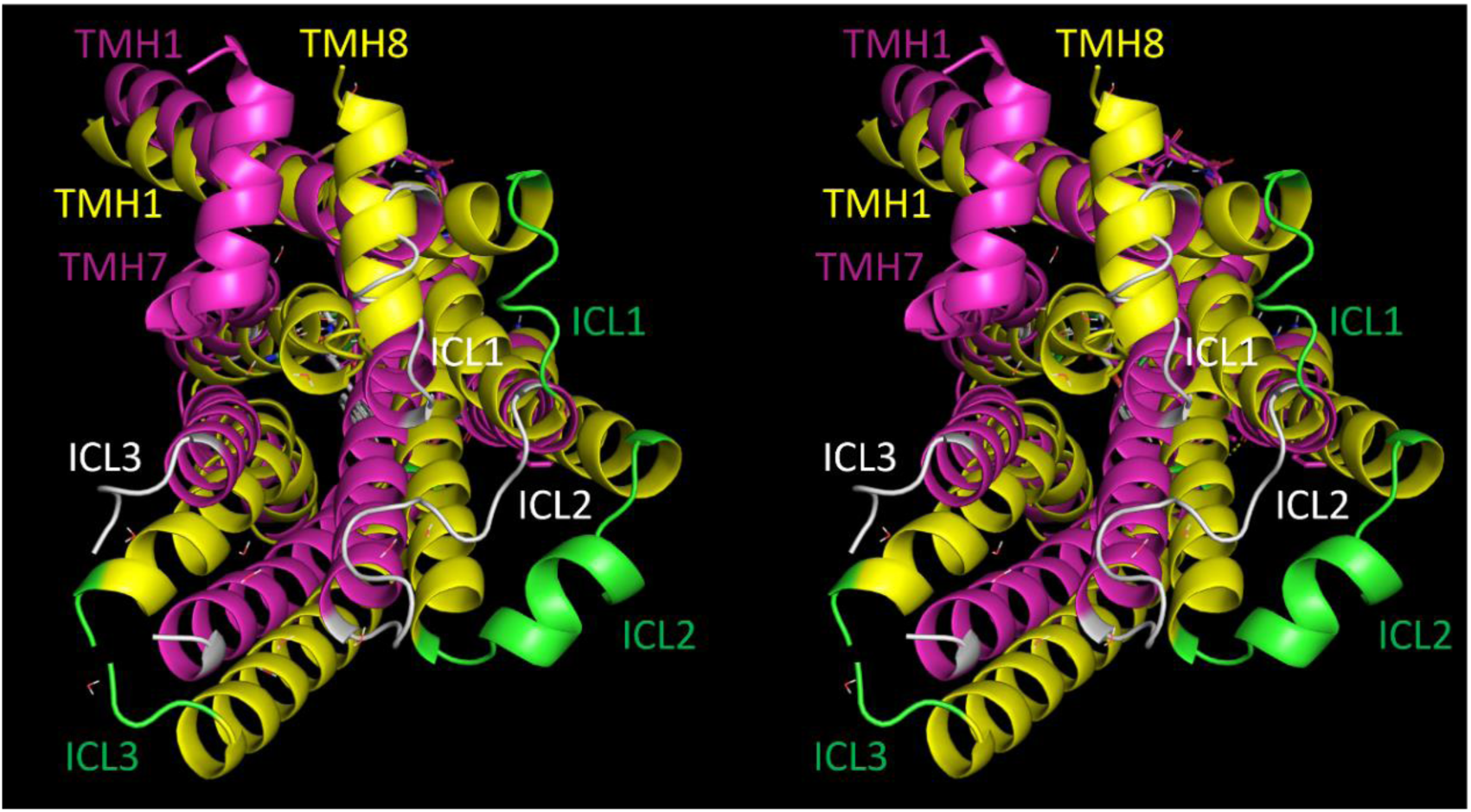
Stereo view of the β_2_-AR, showing the ICLs in the activated (yellow TMHs and green loops) and deactivated states (magenta TMHs and white loops). The ICL conformations adjust in each state to the relative positions of the TMHs to which they interconnect (noting that ICL2 contains an α-helix in the activated, but not the deactivated state, and ICL1 in the deactivated state coincides with the position of TMH8 in the activated state).

We set about to explore activation of the β_2_-AR, which is conveyed by a cascade of rigid-body TMH pivoting/translations and tropimeric rearrangements within the 7-TMH bundle (Figures 13 and 14), beginning with first order H-bonding between a bound agonist and TMH3, TMH5, and TMH7. TMH3 and TMH5 are pushed away from the agonist, while THM7 is pulled toward it into an internal solvation cavity (noting that such rearrangements are precluded by high complementarity between antagonists and the deactivated orthosteric pocket). This is followed by second order (knock-on) effects, largely consisting of rigid-body pivoting of TMH1, TMH2, TMH4, and TMH6. The cascade flows largely along the ECL and ICL loop directions. These rearrangements are many-bodied, and therefore, likely highly non-linear (i.e., non-sequential).

**Figure 13.**
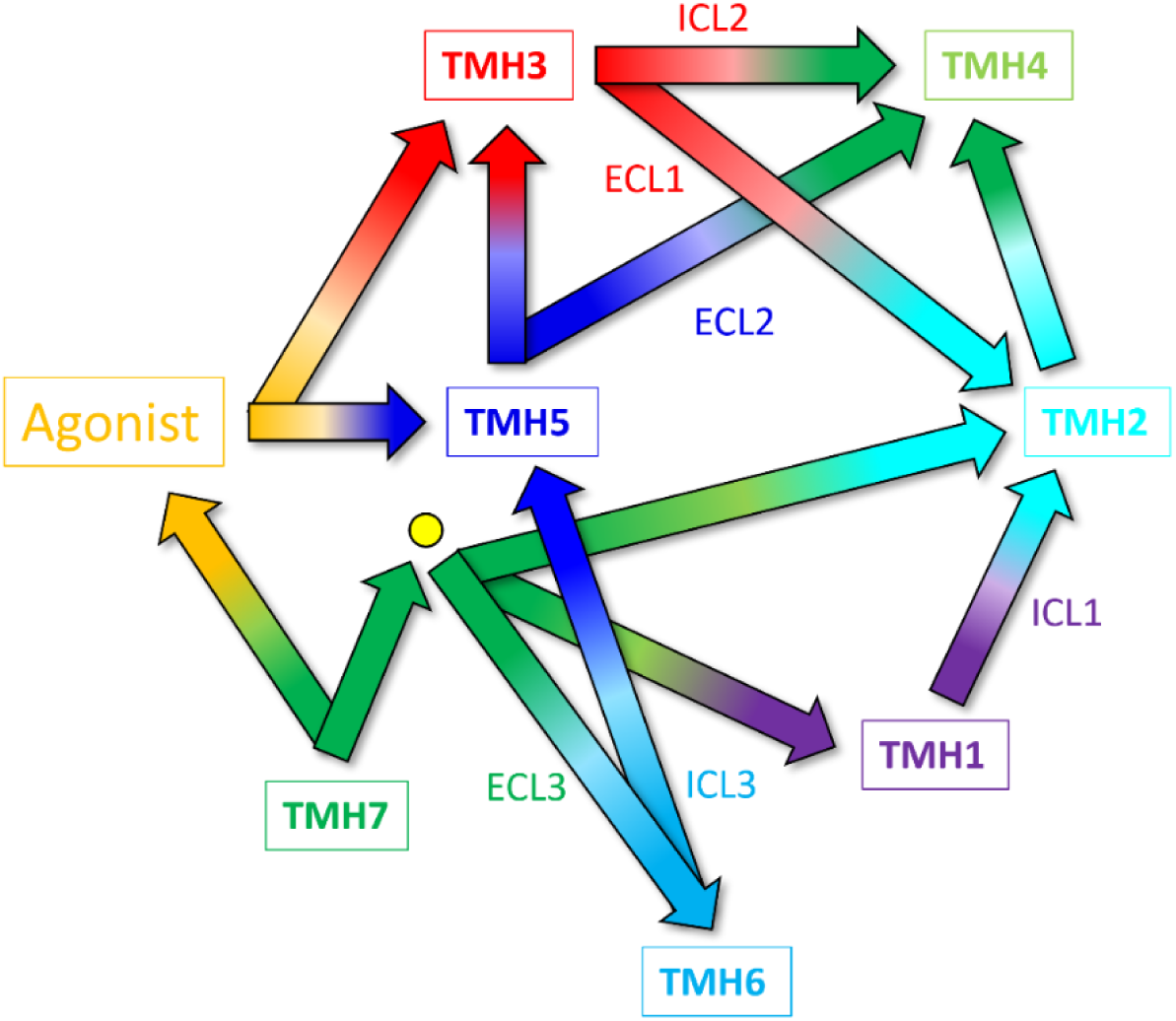
Activation of the β_2_-AR (and likely all class A GPCRs) is conveyed by a cascade of rigid-body and tropimeric rearrangements within the 7-TMH bundle, beginning with agonist H-bonding to partners on TMH3, TMH5, and TMH7 (first order effects annotated by solid orange arrows). Second order knock-on effects, consisting largely of rigid-body pivoting of TMH1, TMH2, TMH4, and TMH6, are annotated by solid arrows color-coded to the penultimate TMH in the cascade. A Na^+^ ion that is putatively present in the deactivated state is depicted as a yellow circle.

**Figure 14.**
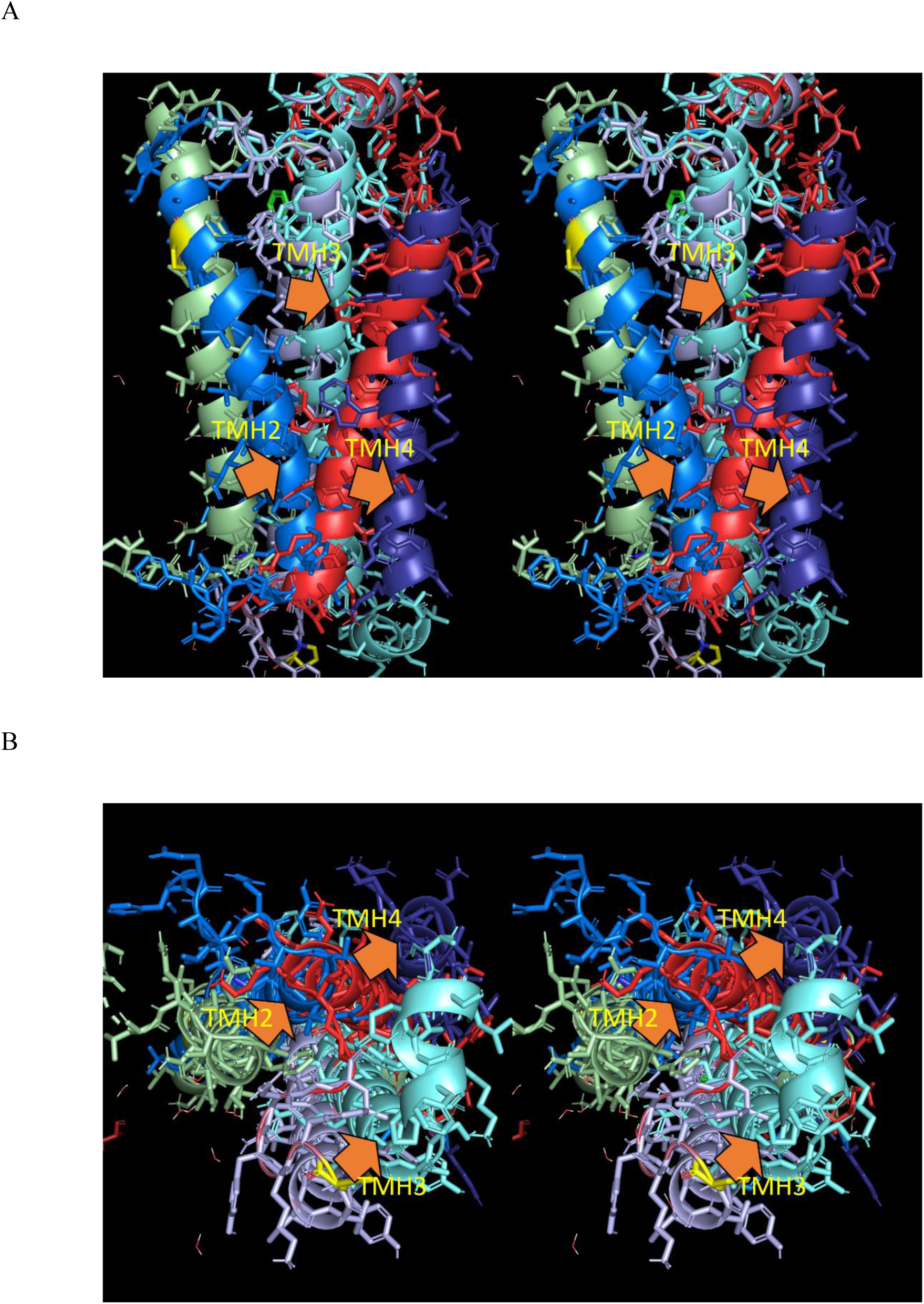
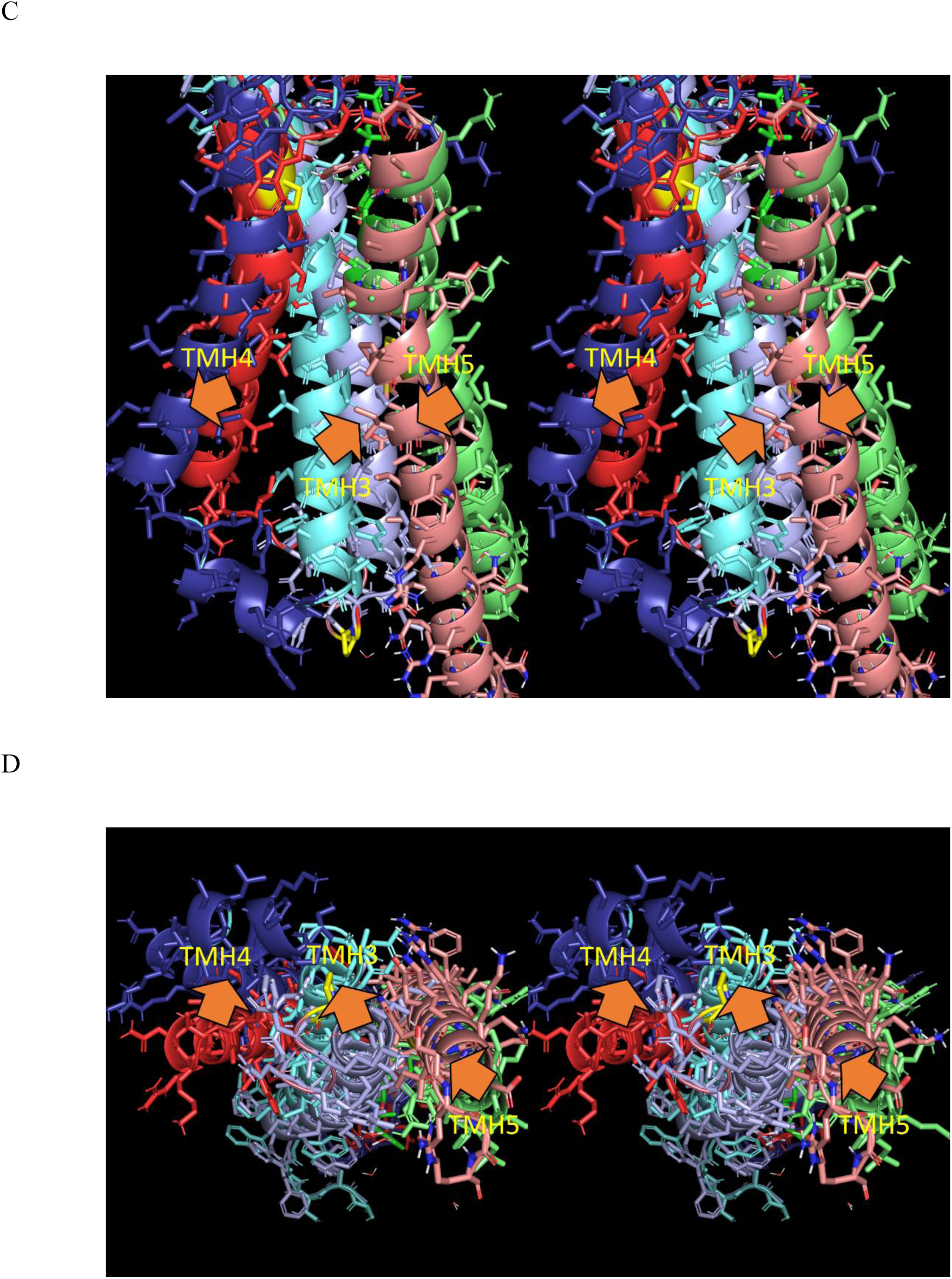
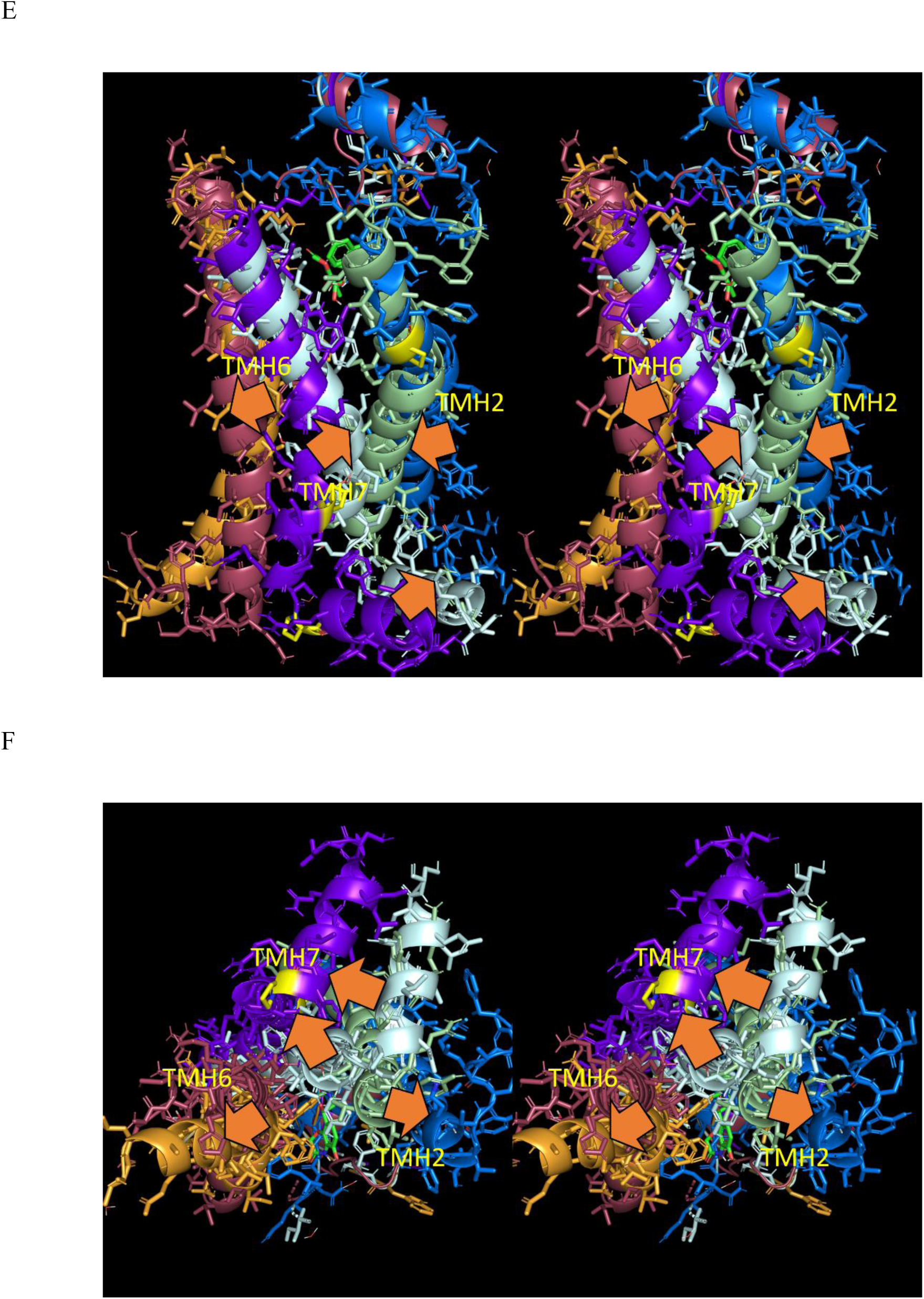
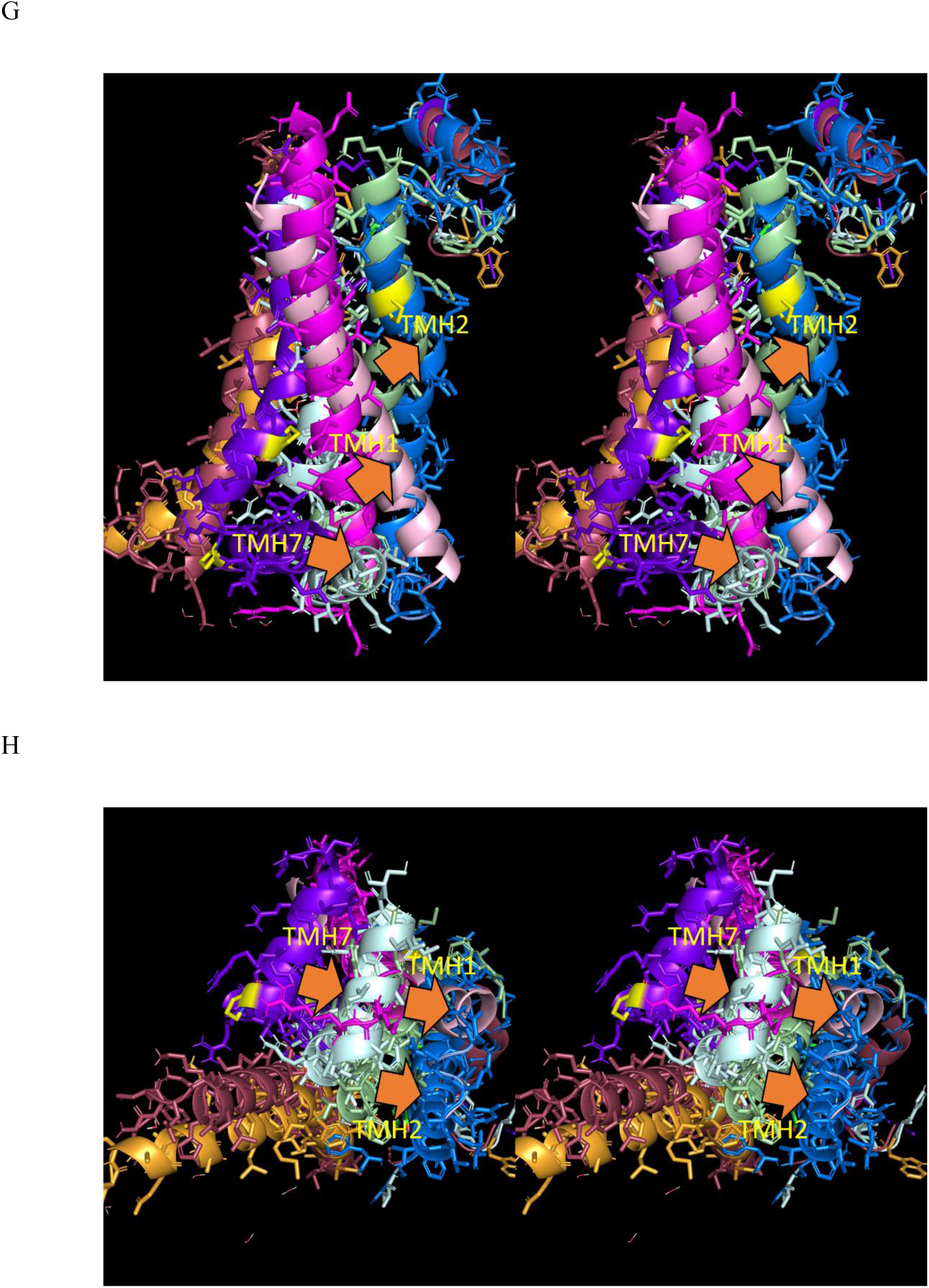
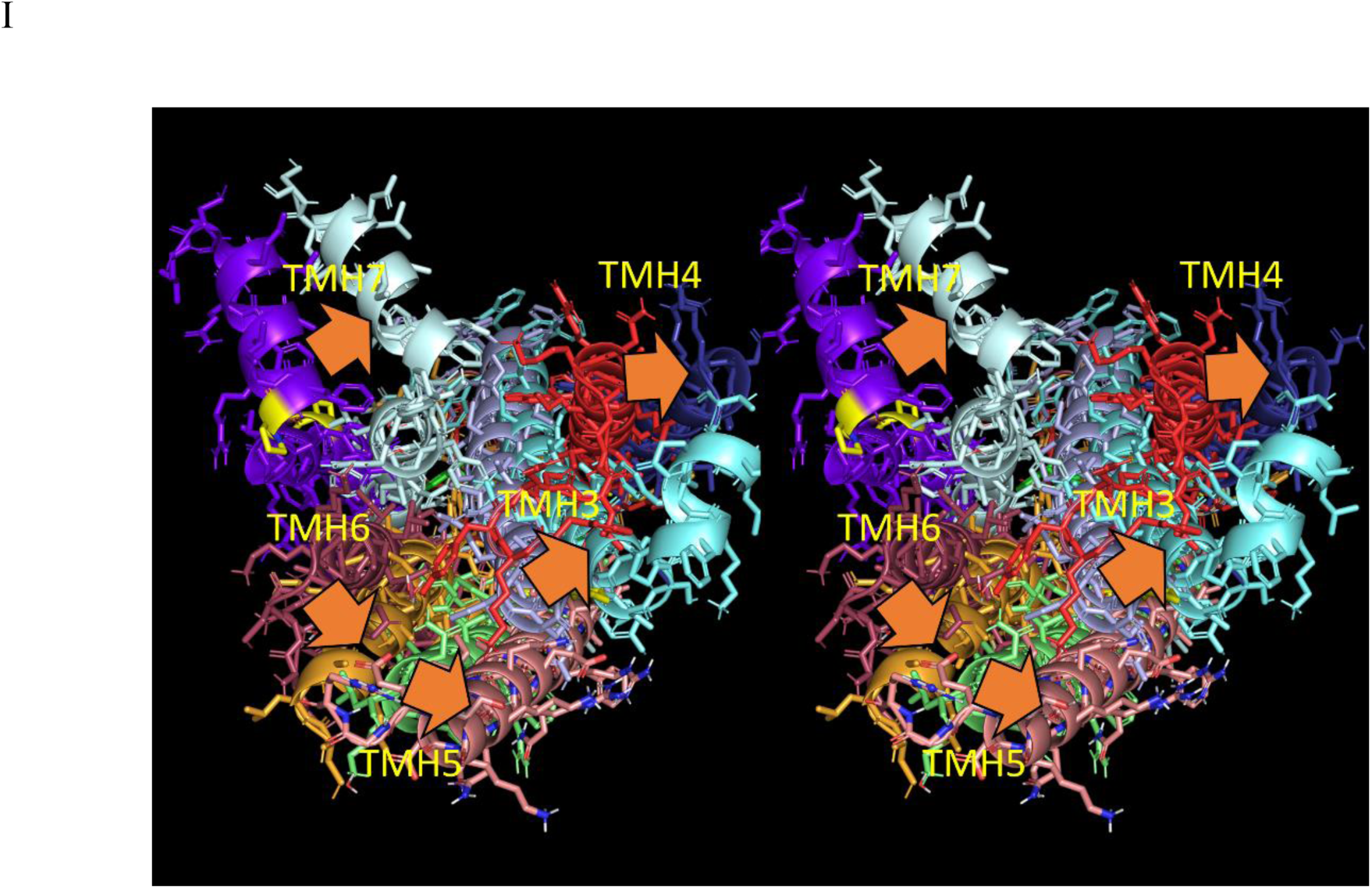
Stereo views of the second order rearrangements of the TMHs in the β_2_-AR, color-coded as follows: Deactivated TMH1 (magenta), TMH2 (light green), TMH3 (cyan), TMH4 (red), TMH5 (green), TMH6 (brown), TMH7 (purple). Activated TMH1 (pink), TMH2 (blue), TMH3 (violet), TMH4 (dark blue), TMH5 (salmon), TMH6 (tan), TMH7 (aqua). (A) Deactivated TMH3/TMH2/TMH4 → activated transitions viewed approximately perpendicular to the helix axes. (B) Same as A, except viewed from the IC side parallel to the helix axes. (C) Deactivated TMH3/TMH5/TMH4 → activated transitions (perpendicular view). (D) Same as C (parallel view). (E) Deactivated TMH7/TMH2/TMH6 → activated transitions (perpendicular view). (F) Same as E (parallel view). (G) Deactivated TMH7/TMH1/TMH2 → activated transitions (perpendicular view). (H) Same as G (parallel view). (I) Deactivated TMH3/TMH5/TMH7/TM4/TMH6 → activated transitions (parallel view only for the sake of clarity).

#### β_2_-AR activation/deactivation and agonist and G-protein binding are powered principally by the expulsion of H-bond depleted solvation from external and internal surfaces, respectively

In our previous work on COVID M^pro^ ^6^ and cereblon,^7^ we proposed the following general mechanistic attributes of dynamic molecular systems operating in the native cellular setting (summarized in Figure 15):

1. Molecules operate in the non-equilibrium regime, cycling between their functional intra- and intermolecular states over time.
2. Cycling depends on internal cavities and channels that:

a. Serve as free volume for intramolecular rearrangements.
b. Contain H-bond depleted solvation that is trapped within, or that exhibits impeded exchange with bulk solvent (noting that such solvation exists within a Goldilocks zone of unfavorable free energy). Intramolecular rearrangements are powered by the auto-desolvation of such water.
3. Non-equilibrium conditions are maintained throughout the cycle via conservation of the total positive solvation free energy of the system (i.e., new channels/cavities are generated as others are desolvated/destroyed).
4. The deactivated and activated states are separated by a free energy barrier preventing spontaneous transitioning to and from (noting that spontaneous transitioning occurs in constitutively active proteins). This barrier is surmounted by the input of favorable free energy to the system, such as agonist binding in the case of GPCRs and reversal of Δψ_*m*_(*t*) in the case of VGCCs.

**Figure 15.**
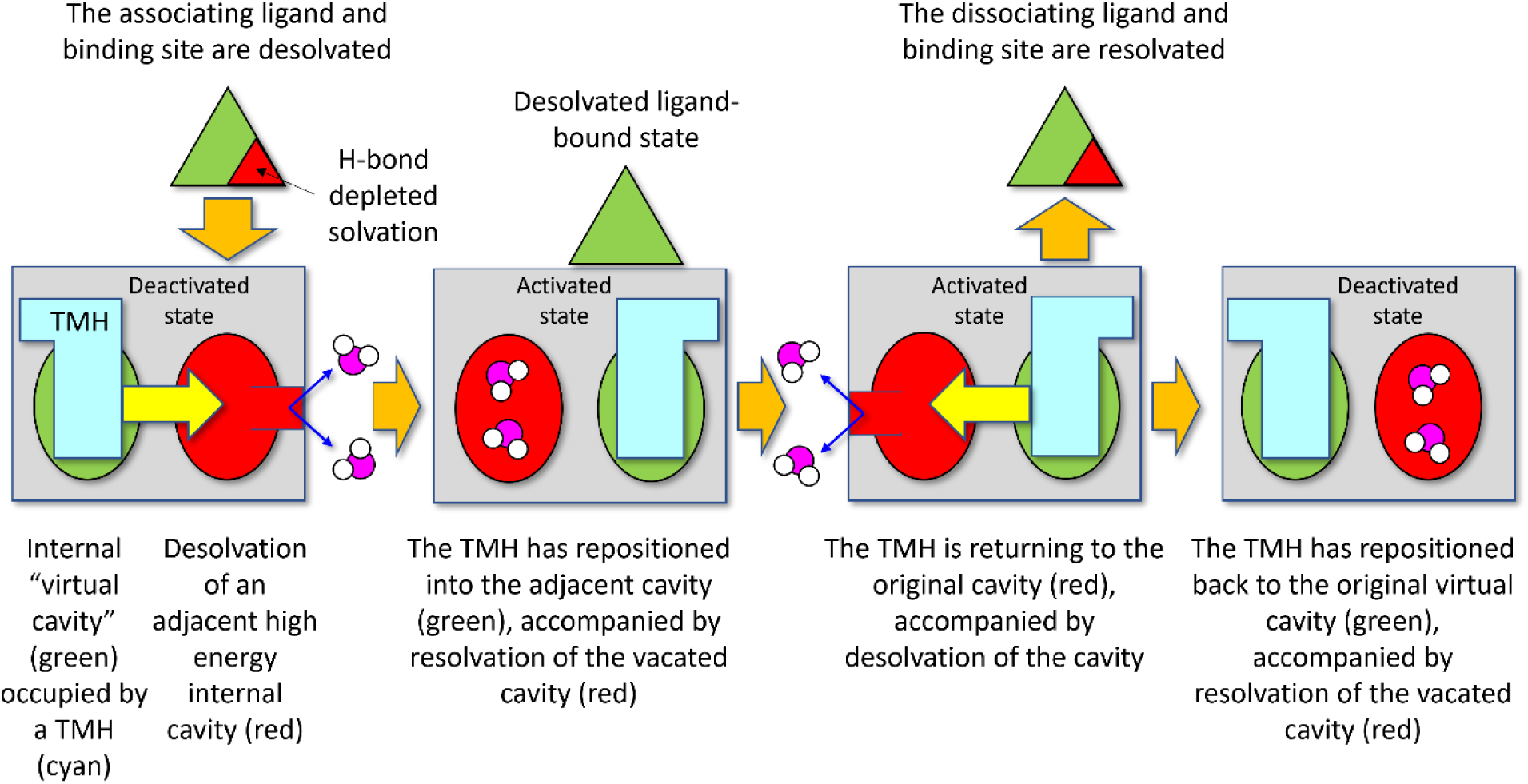
Cartoon depicting a hypothetical cycle in which an intramolecular rearrangement from the deactivated to activated state is powered by desolvation of high energy H-bond depleted water from an internal cavity initiatied by a binding event (a perturbation). Such cavities constitute the free volumes into which the protein rearranges, leaving behind the vacated free volume (solvated by H-bond depleted water) in their wake. Such cavities (the extent of which may be underestimated in crystal structures) necessarily communicate with the protein surface from which the displaced water escapes.

We postulate that internal water channels and cavities exist within GPCRs (and VGCCs), and play similar roles in their structure-function as described above (summarized in Figure 16):

1. The unbound orthosteric pocket contains excess H-bond depleted solvation deduced from the non-polar nature of the side chains lining the pocket (a typical property of binding sites), which is accompanied by impeded or trapped H-bond depleted solvation within the bundle.
2. Agonist/antagonist binding to the orthosteric pocket is powered by mutual desolvation of H-bond depleted ligand and binding site solvation. Additional H-bond depleted solvation is likely expelled during closure of the pocket via activating TMH rearrangements.
3. TMH3, TMH5, and TMH7 undergo <u>first order</u> pivoting into solvation-displaceable internal cavities in response to agonist H-bonding, followed by <u>second order</u> (knock-on) pivoting of the additional TMHs and tropimer curvature that is likewise powered by the desolvation of internal cavities.
4. The allosteric pocket opens via TMH pivoting and tropimer curvature in TMH6 and TMH7, which fills with H-bond depleted solvation needed to slow the G-protein k_off_ (Figure 17) (noting that the cost of generating H-bond depleted solvation in the allosteric pocket is necessarily surmounted by free energy gains from the TMH rearrangements, and as for the orthosteric pocket, resides within a Goldilocks zone of free energy). The G-protein k_off_ is necessarily ≤ the G-protein activation rate. The open allosteric pocket likely persists for multiple G-protein activation cycles due to agonist trapping within the activated receptor.

**Figure 16.**
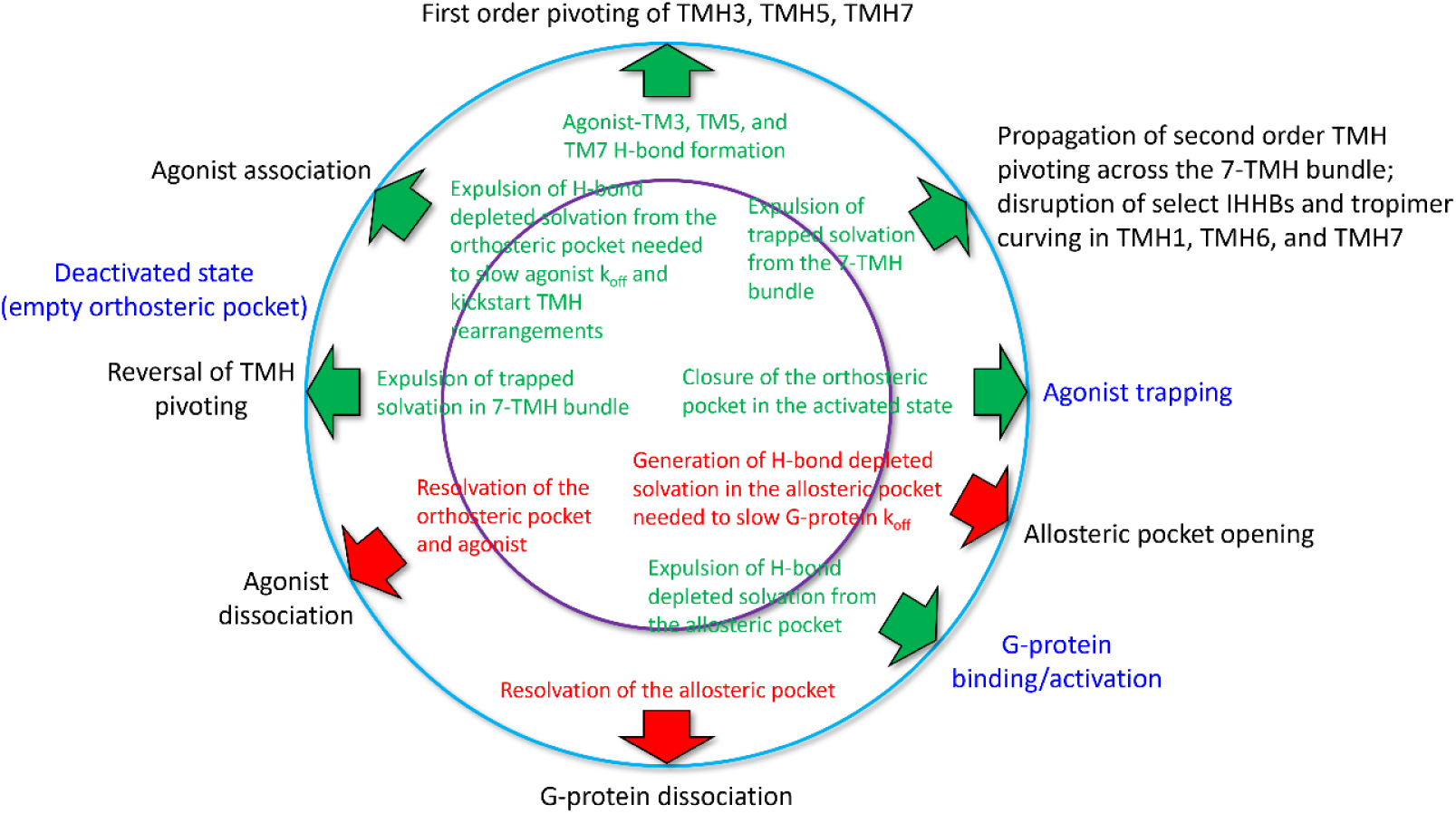
The putative free energy cycle powering the non-equilibrium state transitions of GPCRs deduced from Biodynamics theory. Activating rearrangements of the 7-TMH bundle are kickstarted by agonist binding, which lowers the free energy barrier to the downstream rearrangements that are powered by the expulsion of internal H-bond depleted solvation (see text).

**Figure 17.**
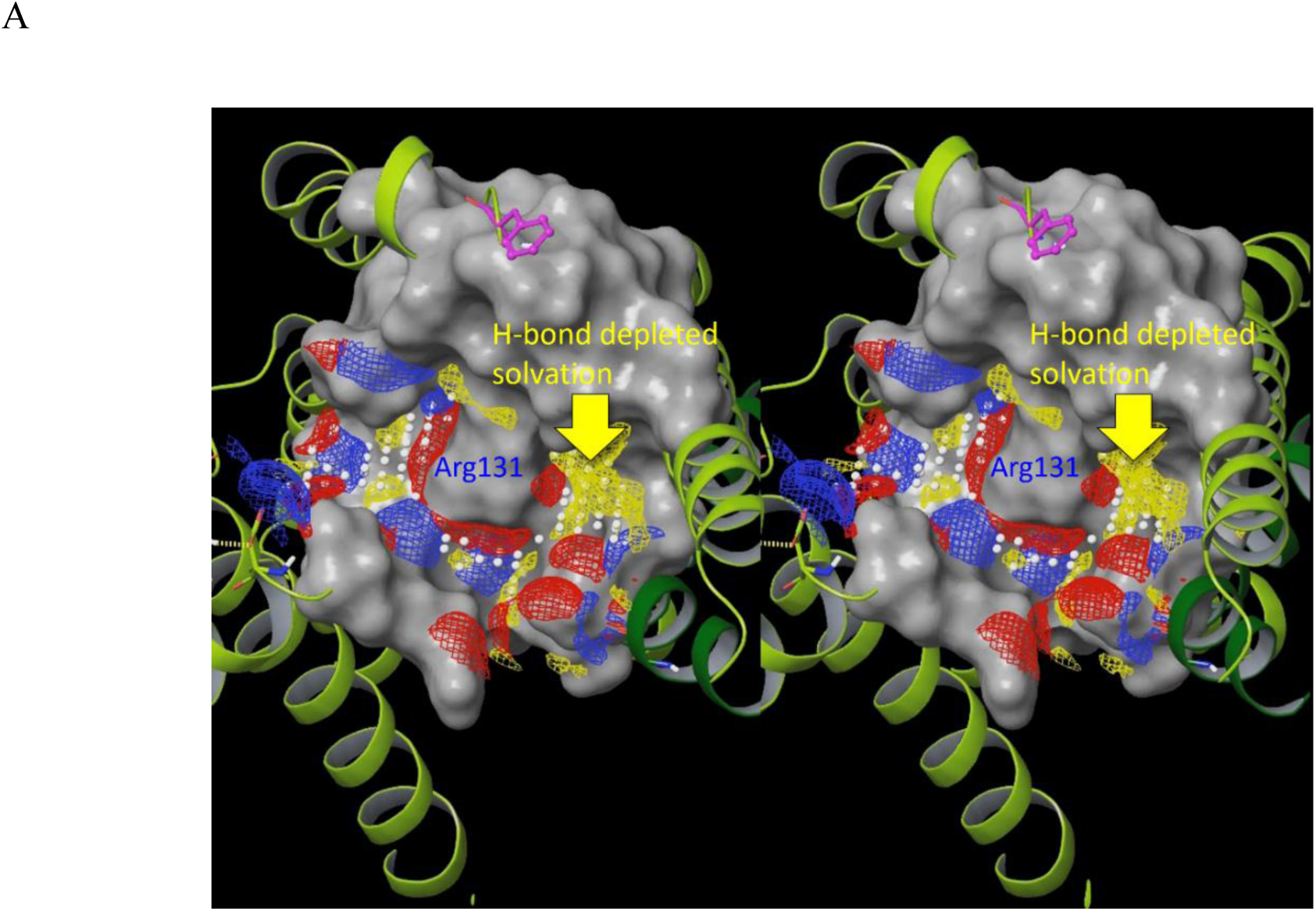
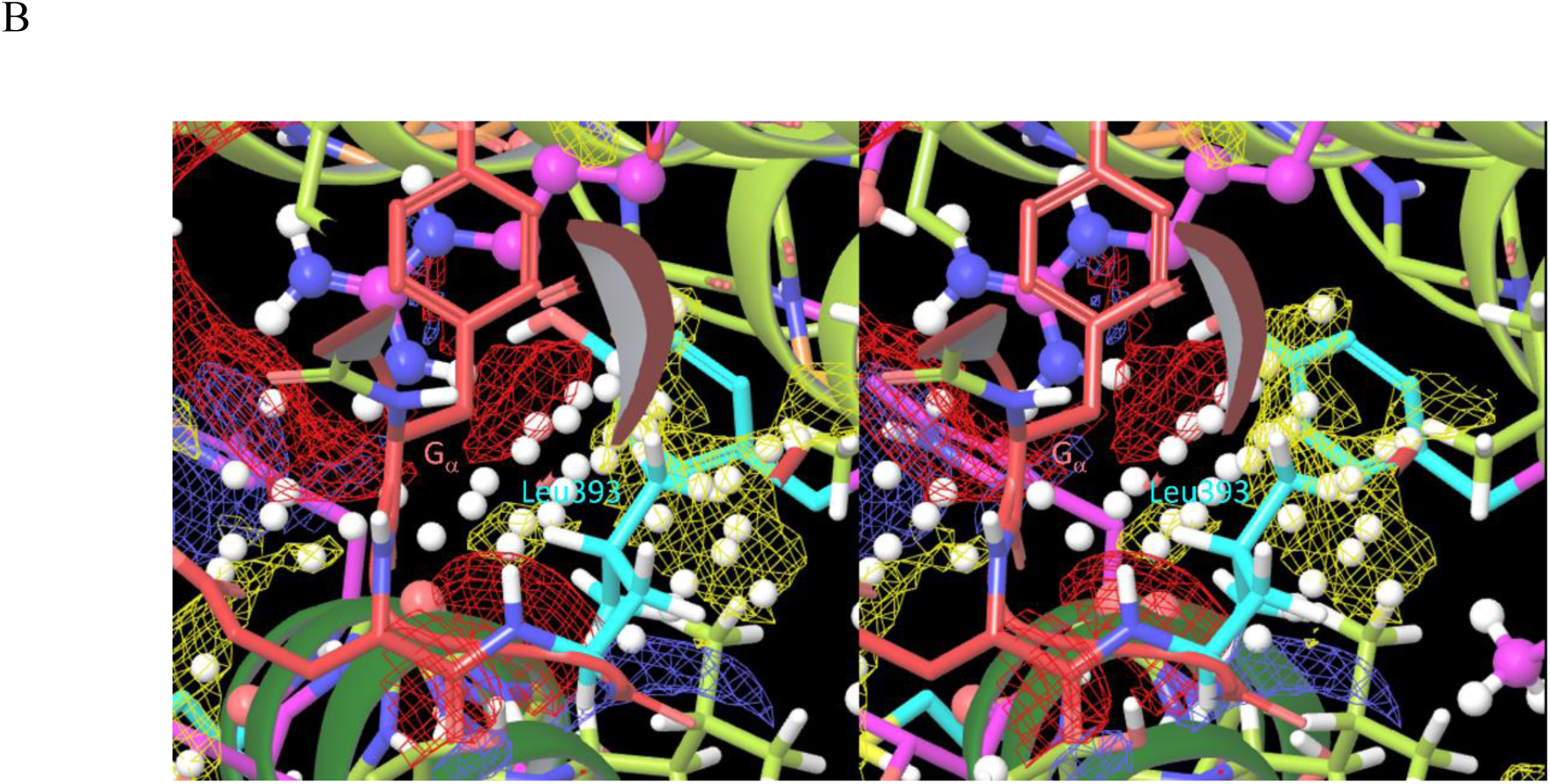
(A) Stereo view of H-bond depleted solvation (yellow contour) within the G_a_ binding site of the b_2_-AR between TMH3, TMH5 and TMH6 is predicted by SiteMap (see Materials and methods). (B) Closeup stereo view of the G_α_ α-helix overlaid on the Sitemap contours in A, in which the side chain of the highly conserved Leu393 (cyan) on the G_α_ helix (coral) projects into the yellow Sitemap contour shown in A, which desolvates this region and slows k_off_ of the G-protein.

A Na^+^ ion is present at this position in the protease activated receptor 1 (3VW7), which may likewise be present but unresolved in the deactivated β_2_-AR (2RH1) (noting that the first three residues listed above are highly conserved throughout the class A superfamily^23,36,37^). Rearrangement of these residues in the activated β_2_-AR (3SN6) is consistent with repositioning or expulsion of this ion during receptor activation. A water filled cavity exists between TMH2, TMH6, and TMH7 in the deactivated receptor (2RH1) that putatively opens at the IC surface, and is stabilized by the side chains of Asp79 (TMH2), Asn318 (TMH7), Asn322 (TMH7), Trp286 (TMH6), Ser319 (TMH7), and Tyr326 (TMH7) (noting that internal H-bond depleted solvation is maintained within the Goldilocks zone of unfavorability via water-receptor H-bonds) (Figure 18).

**Figure 18.**
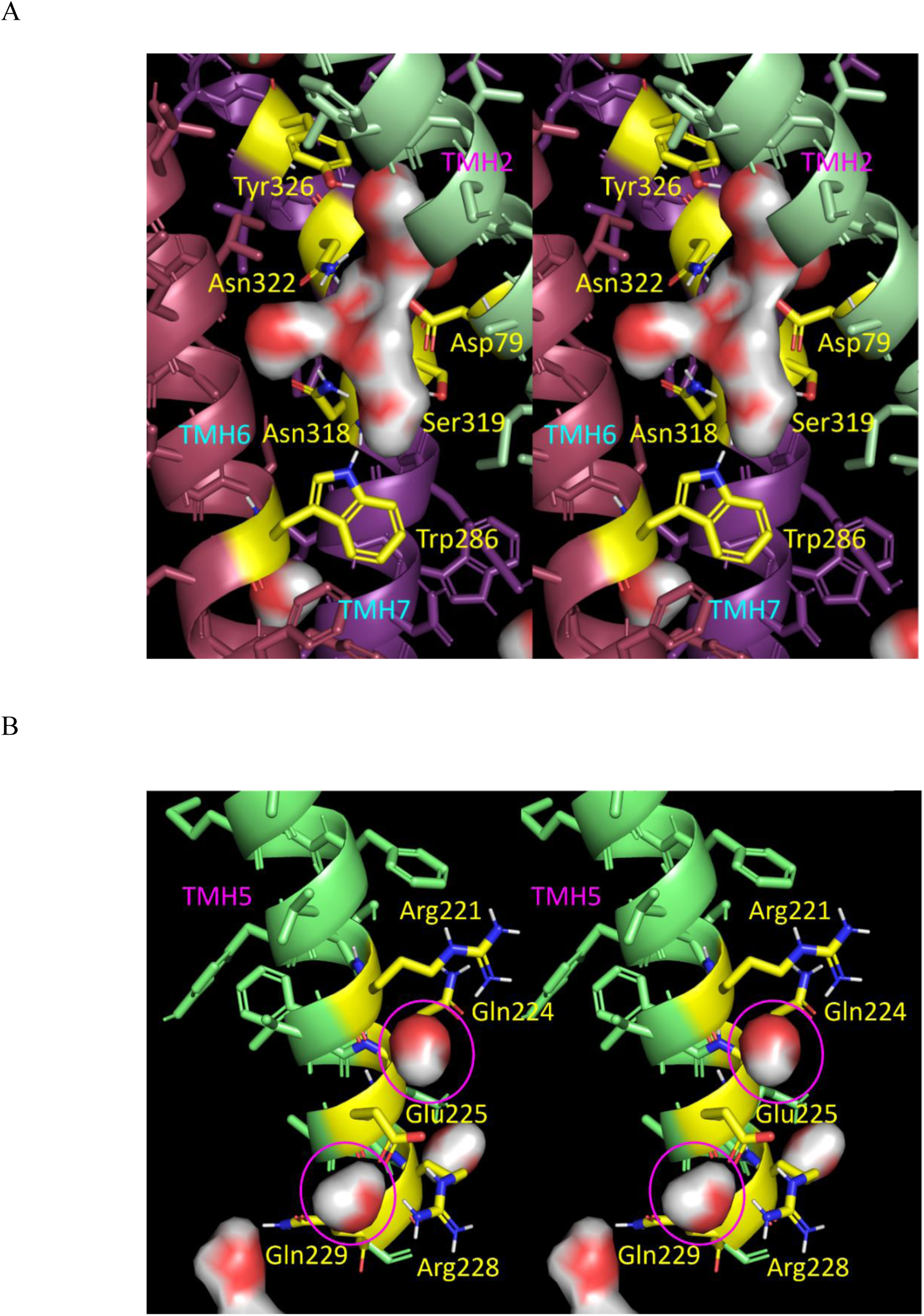
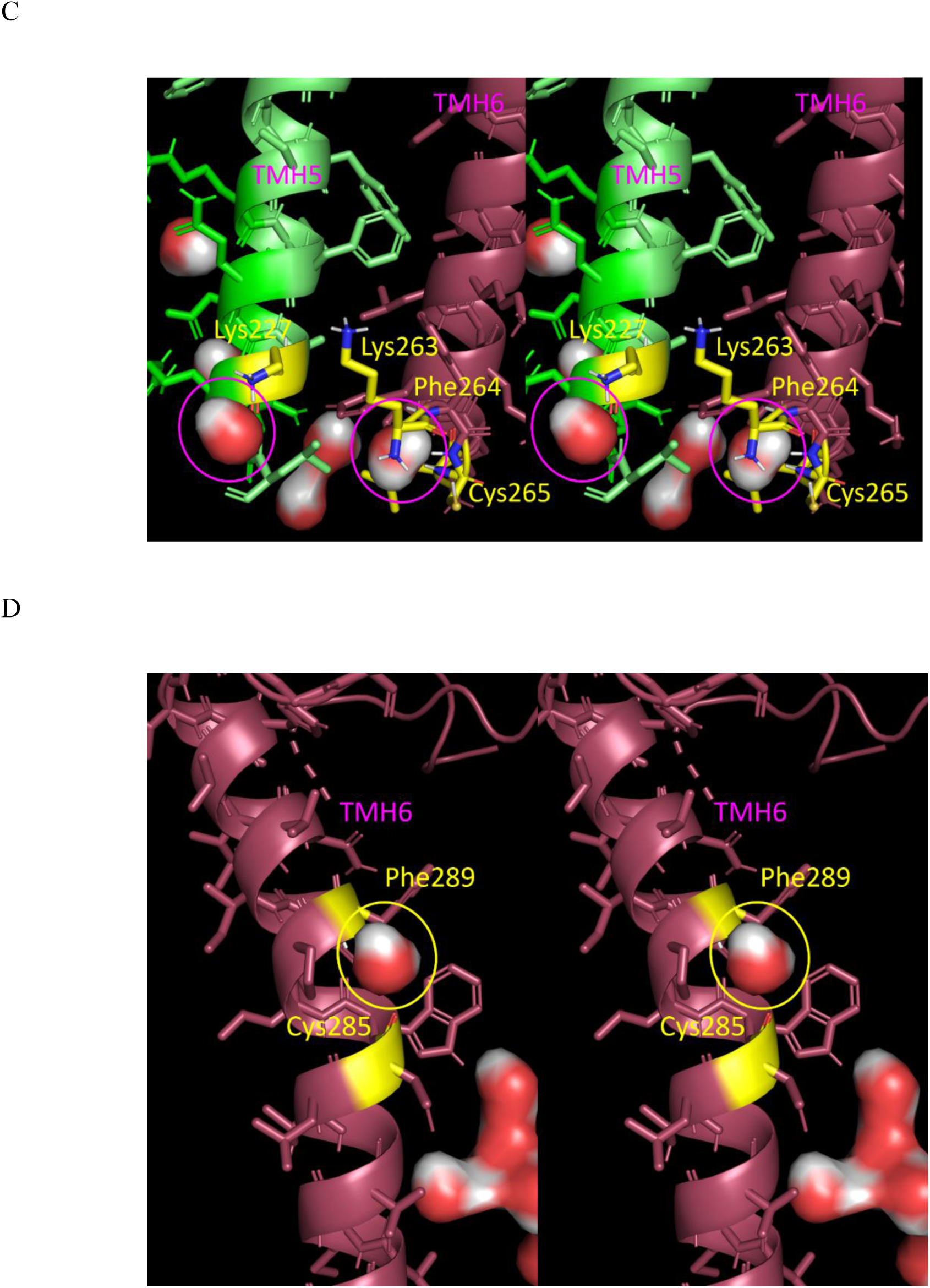
Stereo views of the high energy internal solvation present in the <u>deactivated</u> receptor (2RH1) that both powers activating TMH rearrangements and provides free volume for TMH pivoting and other rearrangements. (A) The central water channel residing between TMH2, TMH6, and TMH7 putatively containing a displaceable Na^+^ ion (not shown) that is stabilized by the labeled side chains is occupied by TMH7 in the activated receptor. (B) Water cavities (magenta circles) adjacent to TMH5 that are stabilized by the labeled side chains and occupied by TMH5 in the activated receptor. (C) Two water cavities adjacent to TMH5 and TMH6 (magenta circles) that are stabilized by the labeled side chains and occupied by TMH6 in the activated receptor. (D) A water cavity (yellow circle) adjacent to TMH6 that is stabilized by backbone groups of Phe289 and Cys285 and occupied by TMH7 in the activated receptor.

First order TMH pivoting is kickstarted by agonist association and H-bonding to Asn312 (TMH7). TMH7 is pulled toward the agonist (Figure 10B) and into the central water channel (Figure 19A), which is powered directly by desolvation of this channel. TMH5 and TMH6 are likewise directly powered by desolvation of internal cavities (Figures 19B-D).

**Figure 19.**
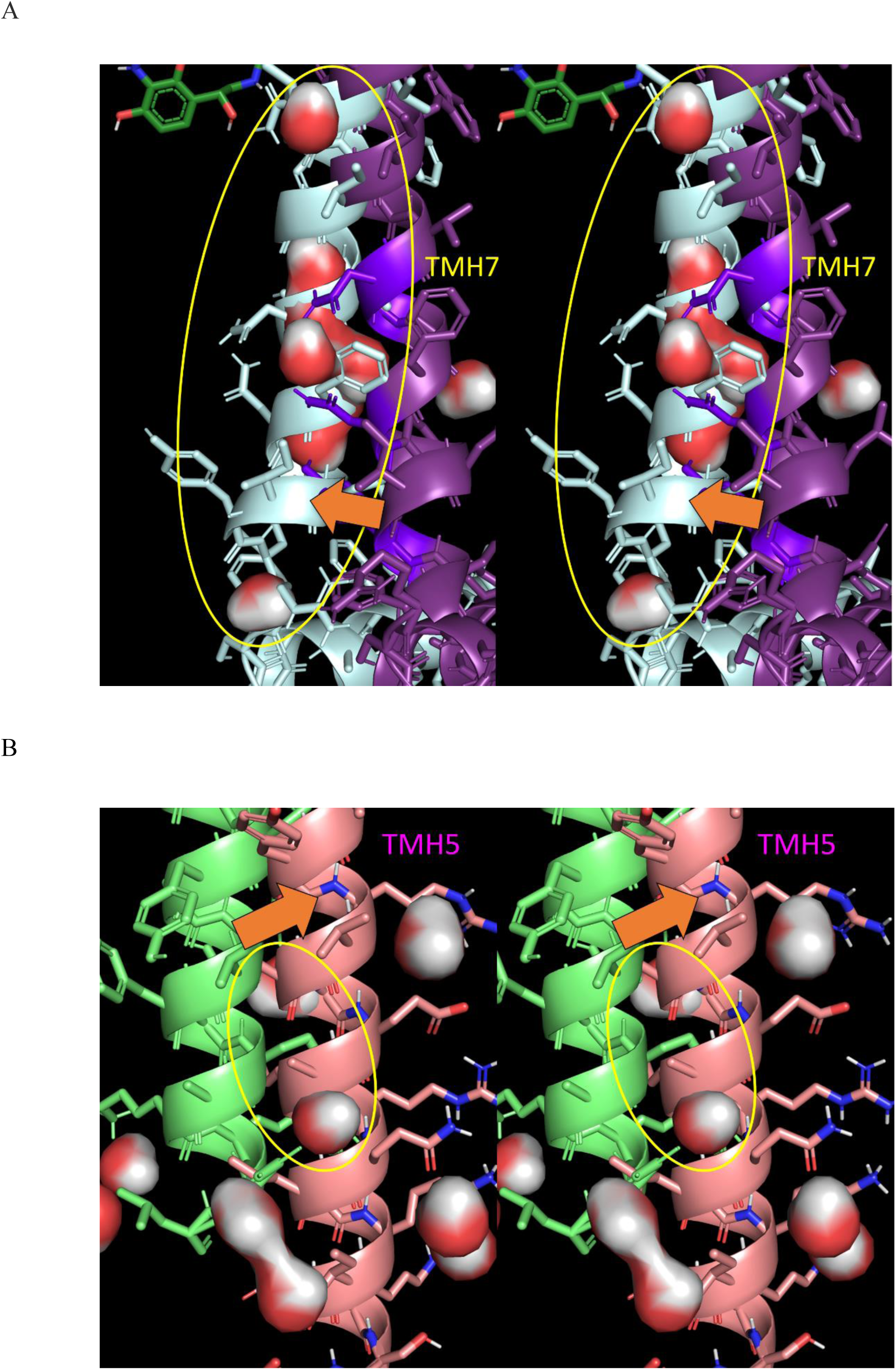
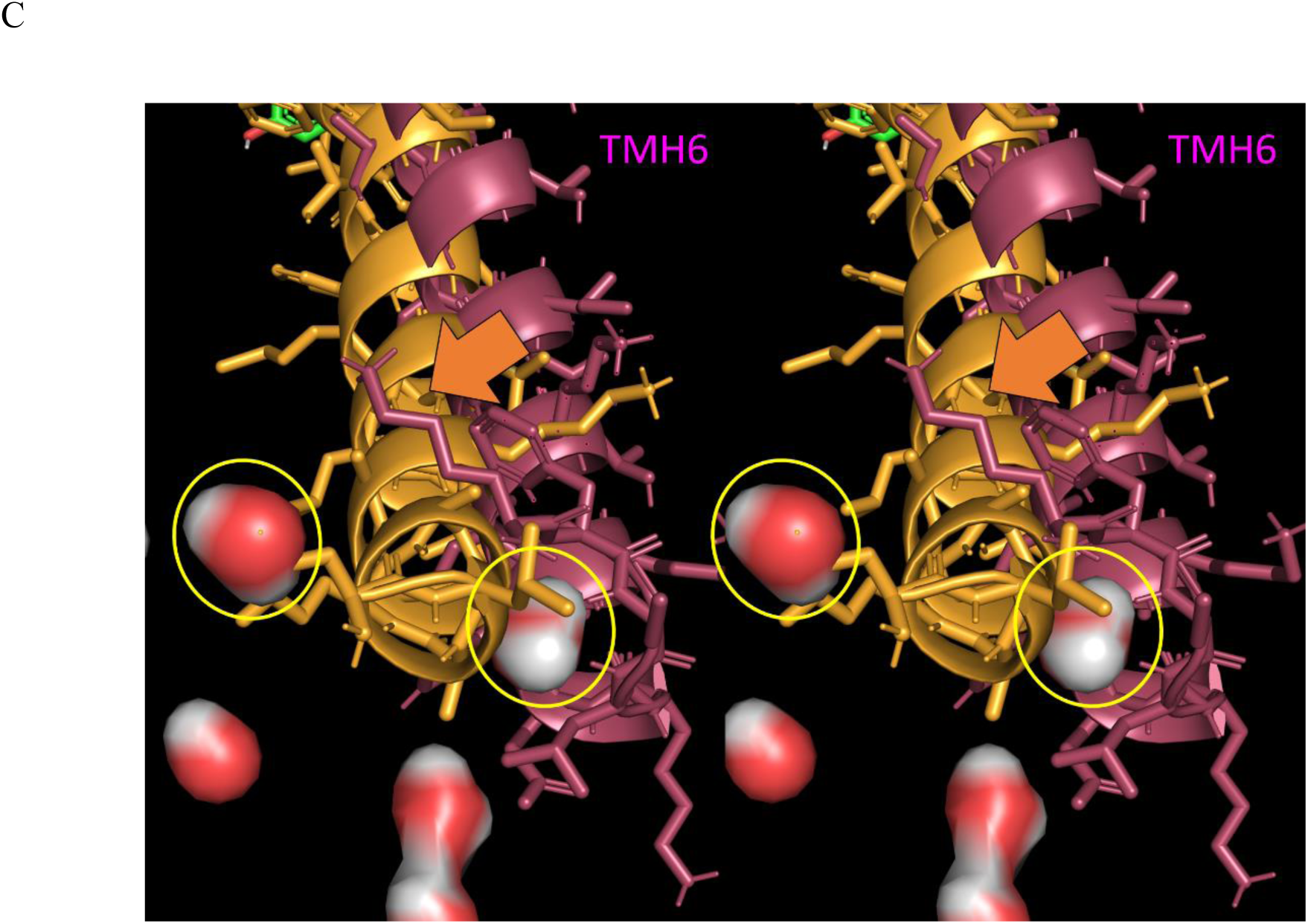
Stereo views of TMH pivoting from the deactivated (2RH1) to the activated state (3SN6) of the receptor. The TMHs pivot into internal water channels/cavities. (A) TMH7 (purple deactivated/light blue activated) pivots into the central water channel shown in Figure 17A. (B) TMH5 (green deactivated/salmon activated) pivots into the two water cavities shown in Figure 17B. (C) TMH6 (tan activated/brown deactivated) pivots into the water cavities shown in Figures 17C and D.

The aforementioned first order rearrangements are followed by second order TMH pivoting (Figure 20). Water is trapped within the orthosteric pocket in the agonist (Figure 21A) and antagonist (Figure 21B) bound states (noting that the k_on_ of the agonist and antagonist present in the two structures is necessarily slowed by this water).

**Figure 20.**
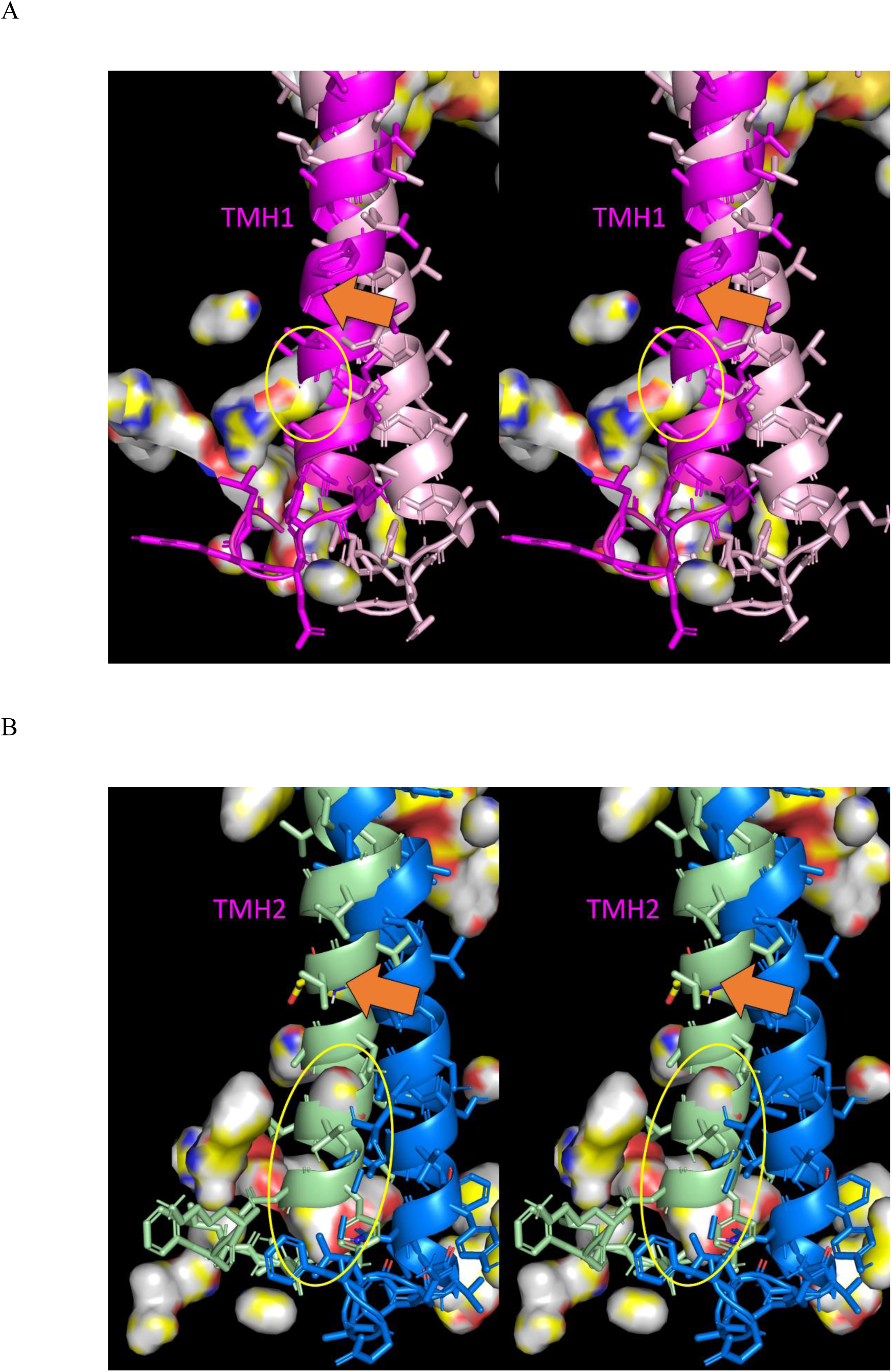
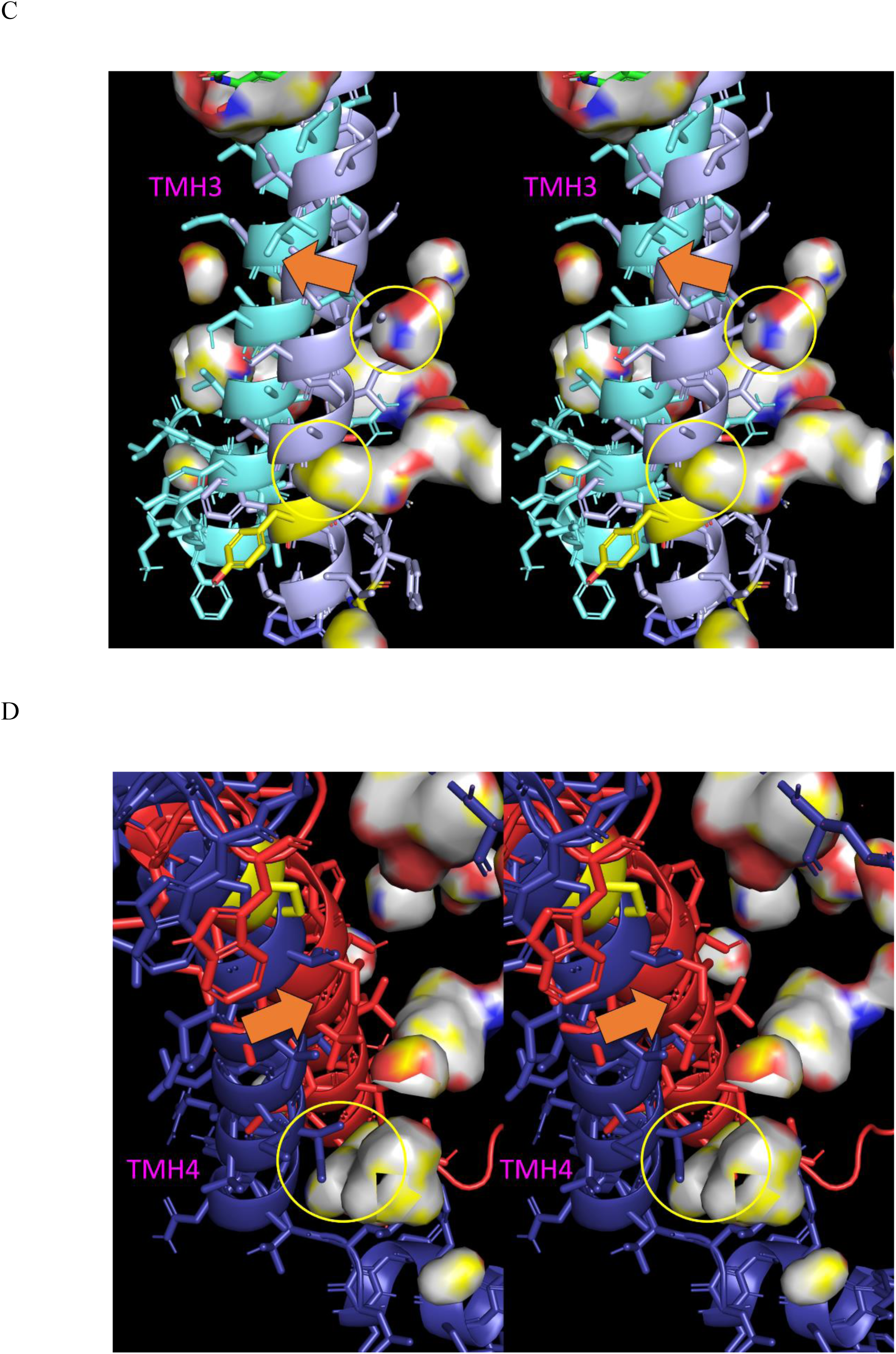
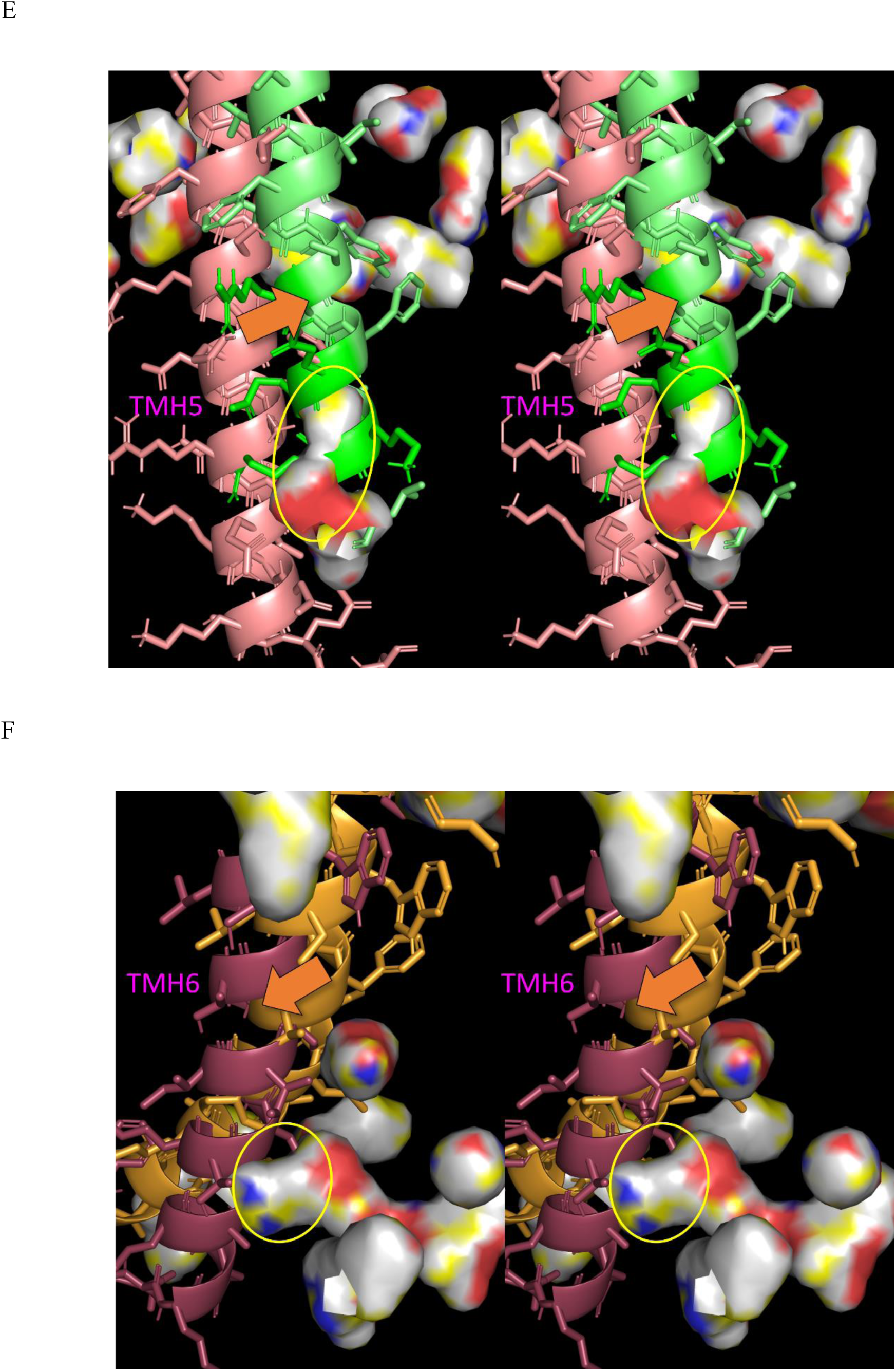
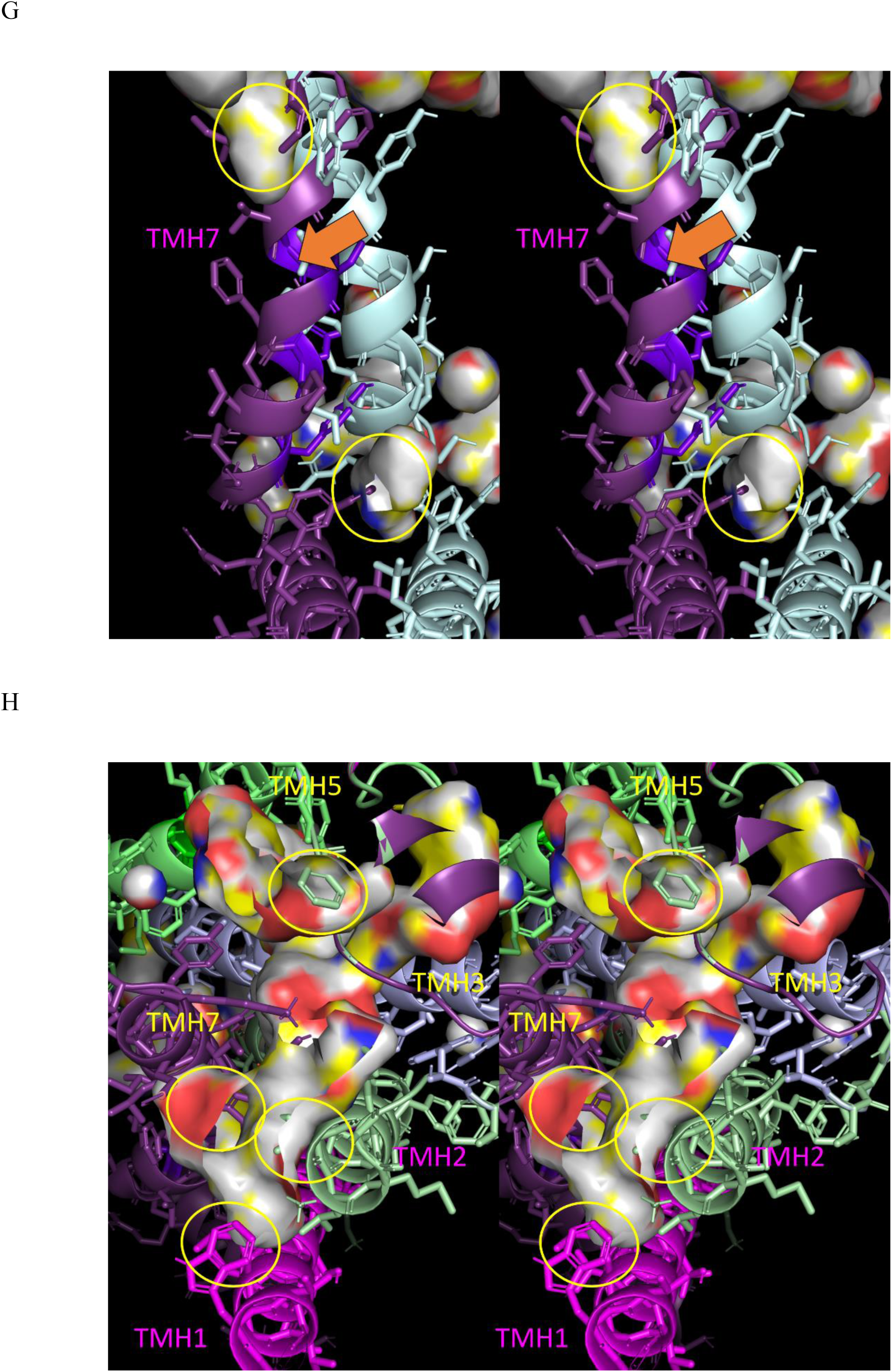
Stereo views of reverse TMH pivoting from the activated (3SN6) to the deactivated (2RH1) state (orange filled arrows). The TMHs pivot into internal water channels/cavities (yellow circles) present in the activated receptor. (A) TMH1 (activated pink/deactivated magenta). (B) TMH2 (activated blue/deactivated light green). (C) TMH3 (activated violet/deactivated cyan). (D) TMH4 (activated dark blue/deactivated red). (E) TMH5 (activated salmon/deactivated green). (F) TMH6 (activated tan/deactivated brown). (G) TMH7 (activated light blue/deactivated purple). (H). Stereo view of TMH pivoting into internal water channels/cavities (yellow circles) on the EC surface (color coding the same as in A-G).

**Figure 21.**
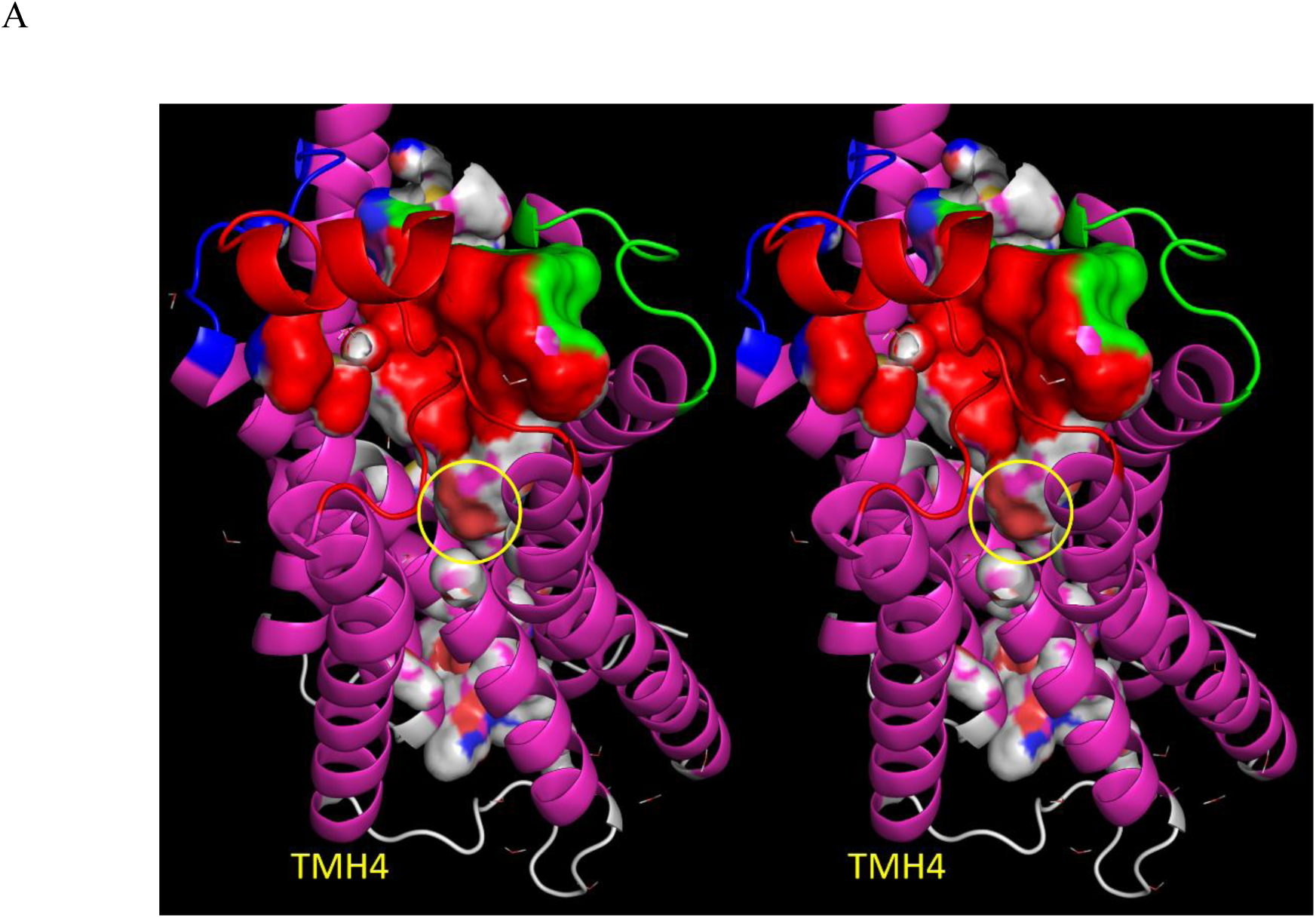
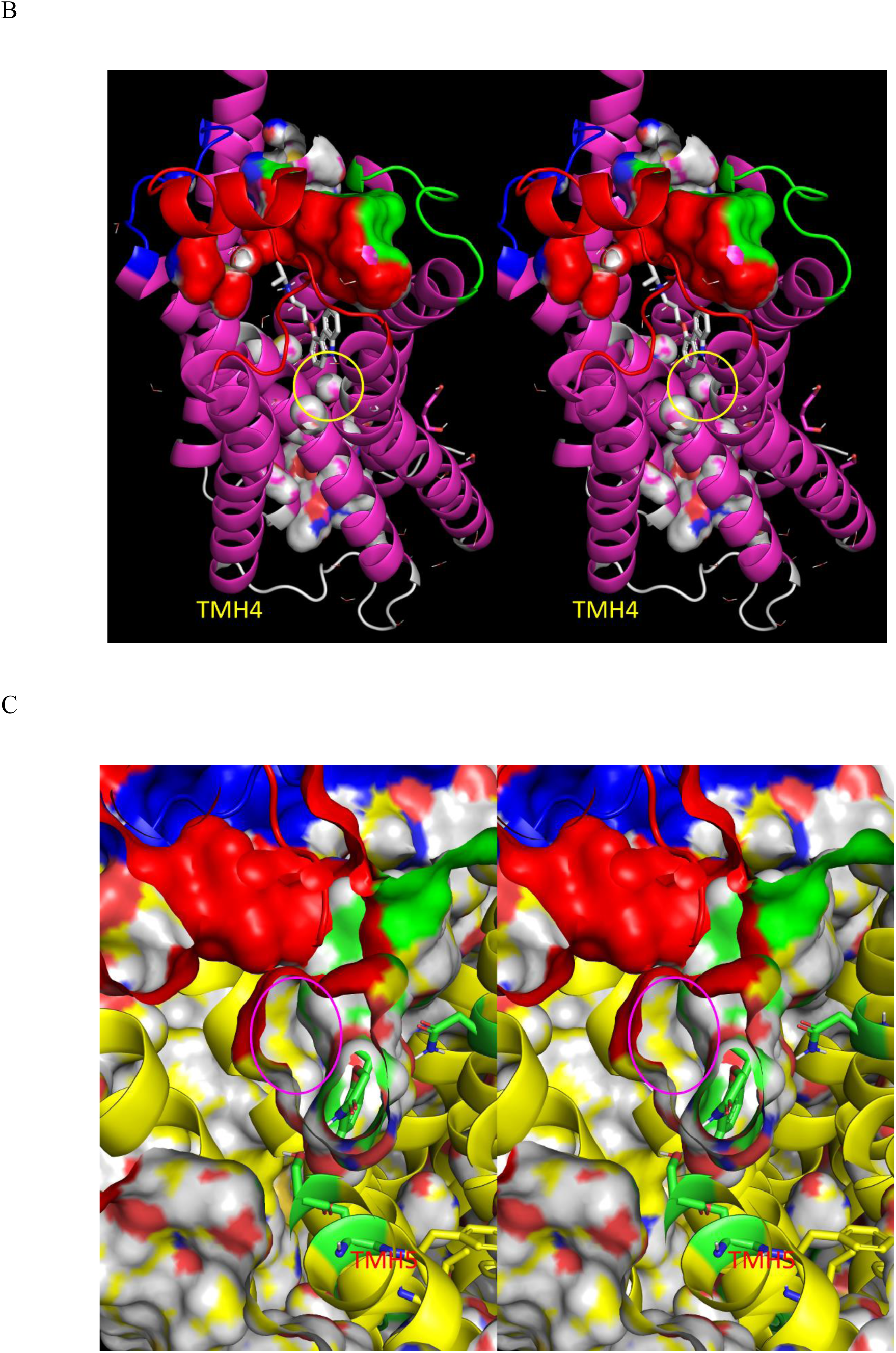
(A) Stereo view of the antagonist (white sticks) bound within the orthosteric pocket of the deactivated receptor (2RH1), showing a water molecule (yellow circle) trapped beneath the antagonist. (B) Same as A, except with the orthosteric pocket surface omitted. (C) Stero view of the agonist (green sticks) bound within the orthosteric pocket of the activated receptor (3SN6), showing an internal water cavity (magenta circle) adjacent to the agonist.

#### The role of helical curvature in GPCR activation

Folding minimizes water-water H-bond losses relative to bulk solvent in aqueous globular proteins and losses in intra- and inter-protein H-bonds in membrane proteins, including the intra-helical H-bonds (IHHBs) of TMHs. Three sets of IHHB partners exist between two adjacent turns of *α*-helices (helical dyads), except in the presence of proline, in which case two IHHB partners are absent (such tropimers are permanently curved to varying degrees/directions, depending on their environment). Tropimeric dyads containing IHHBs that are bilaterally uniform in length are linear (Figure 22A), whereas those containing one or two unilaterally shortened or lengthened IHHBs are curved (Figure 22B) (noting that curvature in <u>isolated</u> helices has been reported by others^38,39^).

**Figure 22.**
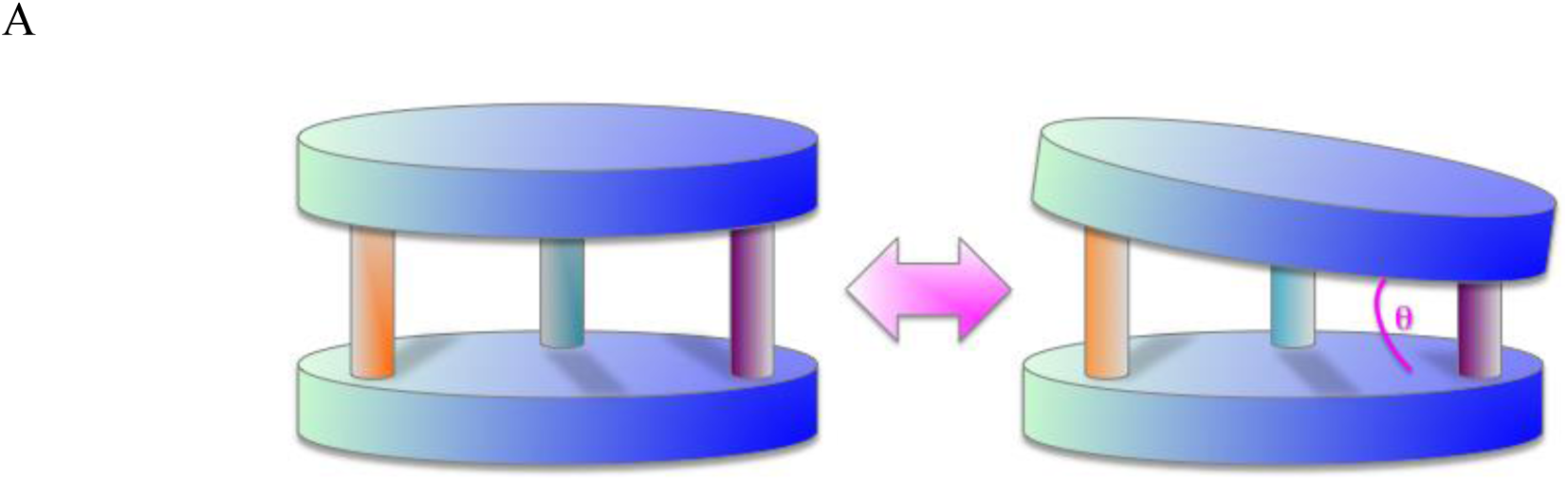
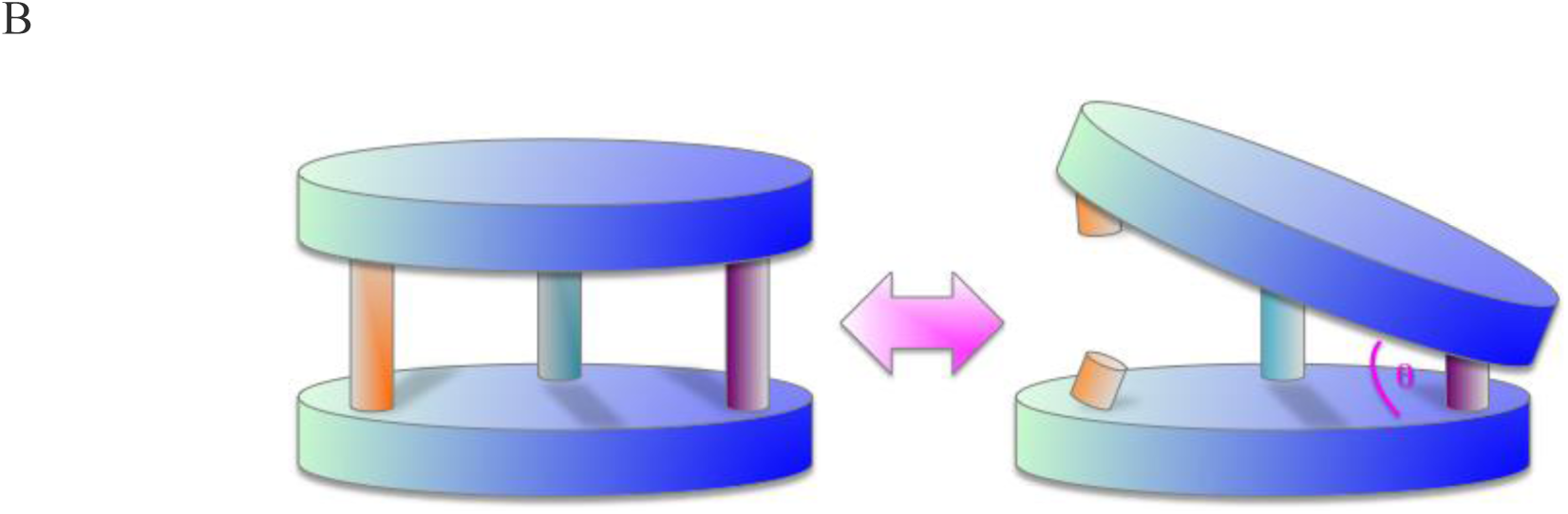
Schematic drawing of a putative tropimer, consisting of two contiguous helical turns (denoted by disks), the intra-helical C=O**^…^**HN H-bond lengths of which (denoted by cylinders) vary asymmetrically across the three sets of intra-helical partners. Tropimers curve in the direction of the shortest helical H-bonds. (A) The angle of attack (θ) depends on the degree of helical H-bond lengthening. (B) Complete loss of intra-helical H-bonding between one or more partners of a tropimer results in the largest θ.

The helical circumference is enlarged within certain curved tropimers (reported by others as helical bulges^40–42)^. Tropimer directionality (referenced to the helix axis direction preceding the tropimer) and angle of curvature θ vary depending on the location(s) of lengthened/disrupted, versus fully H-bonded IHHB lengths, curving toward the shortest H-bond(s). Tropimers are individually or consecutively positioned along certain TMHs (e.g., TMH6 in GPCRs and S6 in VGCCs) (Table 4 and Figure 23).

**Table 4.** Tropimer positions within the activated and deactivated states of the b_2_-AR, all of which are switchable.

| Tropimer | TMH | Disrupted IHHBs | Contains Pro |
| --- | --- | --- | --- |
| 1 | TMH6 | Phe290 O (activated) | No |
| 2 | TMH6 | Phe289 NH, Cys285 O, Leu284 (activated) | Yes |
| 3 | TMH6 | Trp286 NH, Phe282 O, Thr281 O (activated) | No |
| 4 | TMH6 | Ile278 O (activated) | No |
| 5 | TMH6 | Thr274 NH, Ala271 O, Lys270 O (activated) | No |
| 6 | TMH6 | Leu266 O, His269 NH (activated) | No |
| 7 | TMH7 | Ser319 O (activated) | Yes |
| 8 | TMH7 | Leu324 O (activated), Pro323 O (deactivated) | No |

**Figure 23.**
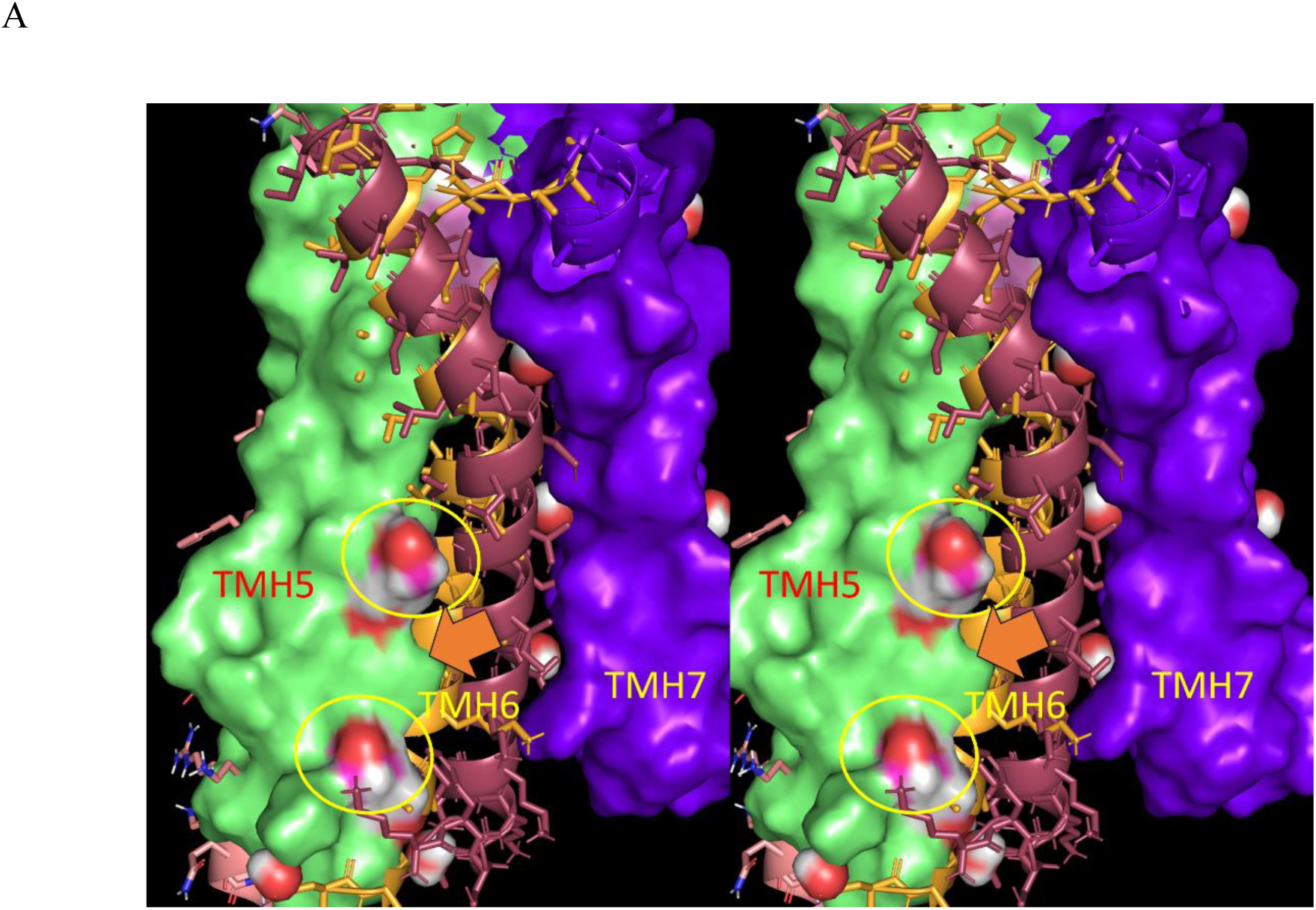
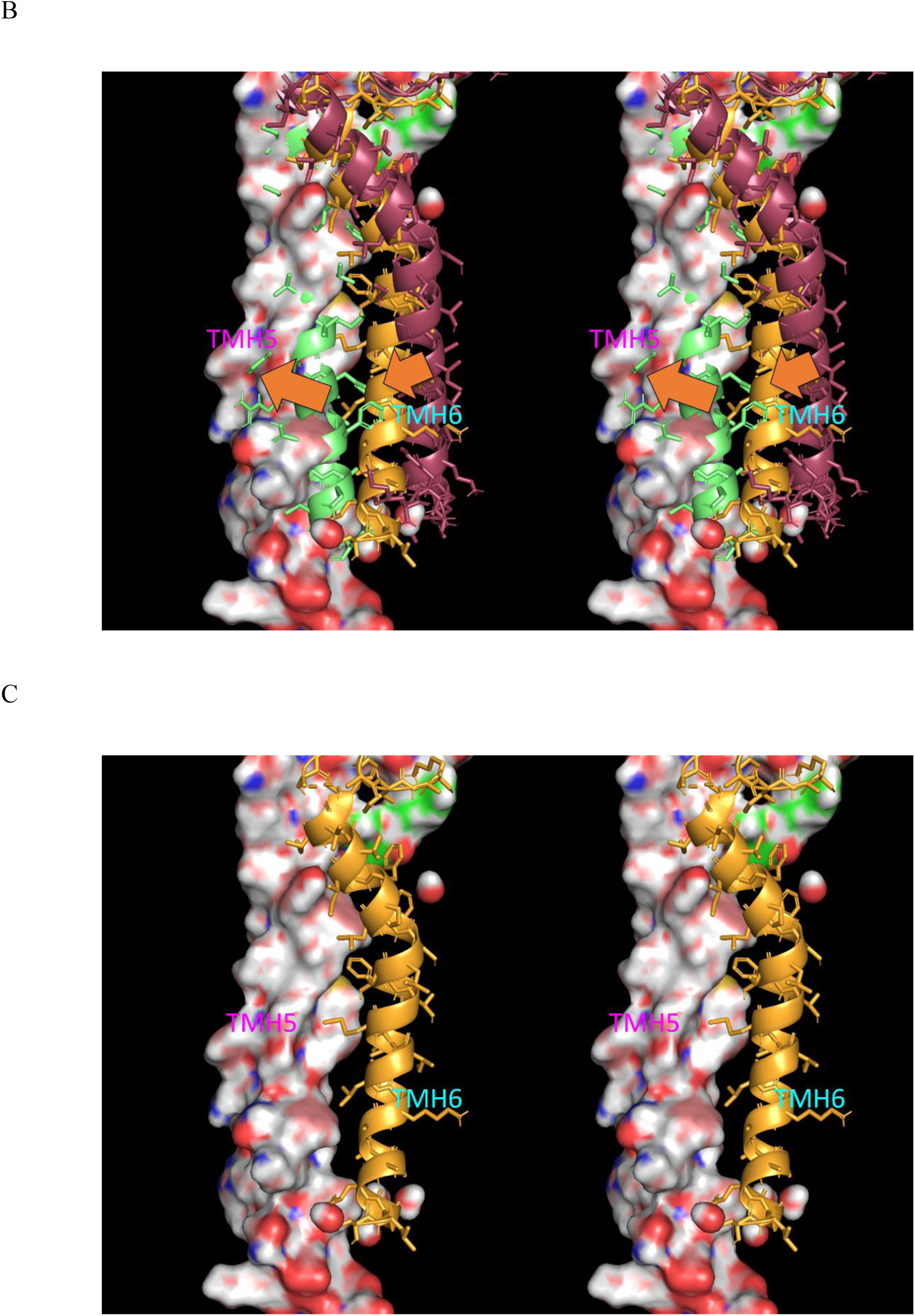
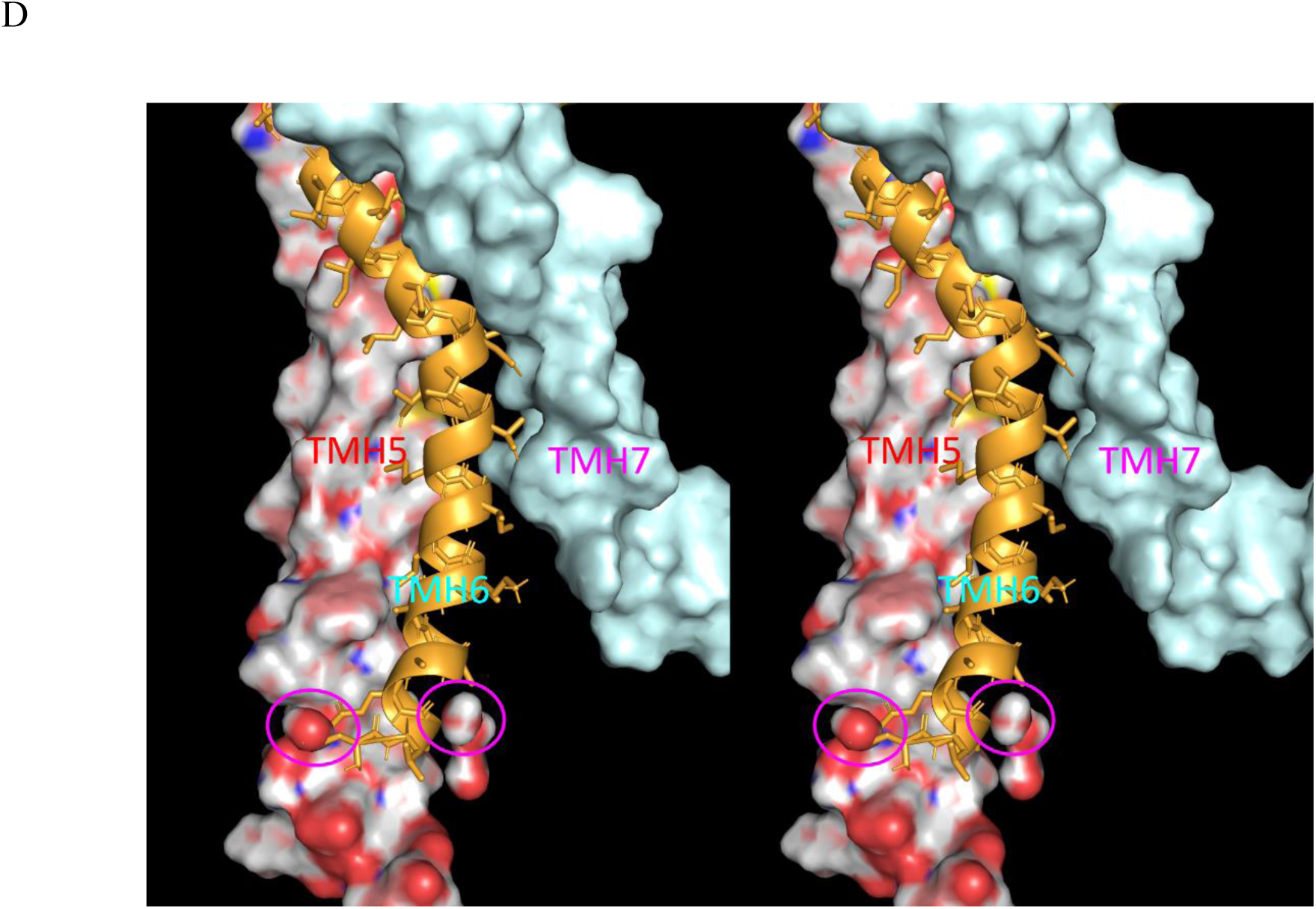
Stereo views of TMH6, showing the contributions to tropimer curvature listed in Table 4. (A) Packing of TMH6 (activated tan/deactivated brown) with TMH5 (green surface) and TMH7 (purple surface). TMH6 pivots (orange solid arrow) into two internal cavities containing 1,4-butanediol, which we assume to be water cavities in the native receptor. (B) Same as A, except showing TMH5 in the activated (salmon surface) and deactivated (green cartoon) states. (C) Same as A, except showing the packing between the activated states of TMH5 and TMH6 (noting that TMH6 pivots with TMH5 so as to maintain packing in the activated state). (D) Same as A, except showing the activated state of TMH7 (blue surface), from which which the IC half of TMH6 has unpacked. TMH6 curvature is powered by desolvation of two internal cavities present in the deactivated state of TMH5 (magenta circles). The desolvation |free energy gain| is necessarily ≥ the cost of IHHB disruption.

We postulate that switchable tropimers contribute to GPCR and VGCC activation and deactivation, as follows:

1. Tropimer curving is conveyed by transient unilateral disruptions of the IHHBs in TMH6 and TMH7 in response to agonist binding in the case of GPCRs, and membrane polarity-induced effects on the S4-S5 linker in the case of VGCCs (described below). The H-bonds of disrupted IHHB partners swap with exo-helical donors or acceptors, including nearby side chains (single or bifurcated), water, or ions on a case-by-case basis. The cost of tropimer curving 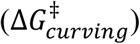 consists of the net loss of H-bond free energy of the swapped intra-helical partners, which we postulate is paid for largely by gains realized from the curving-induced desolvation of internal H-bond depleted water (Figure 24). The lost IHHBs are recovered during exit from the activated state, resulting in tropimer straightening.
2. The free energy difference between the linear and curved tropimeric states is lowered by strategically positioned proline residues containing permanent IHHB disruptions.

**Figure 24.**
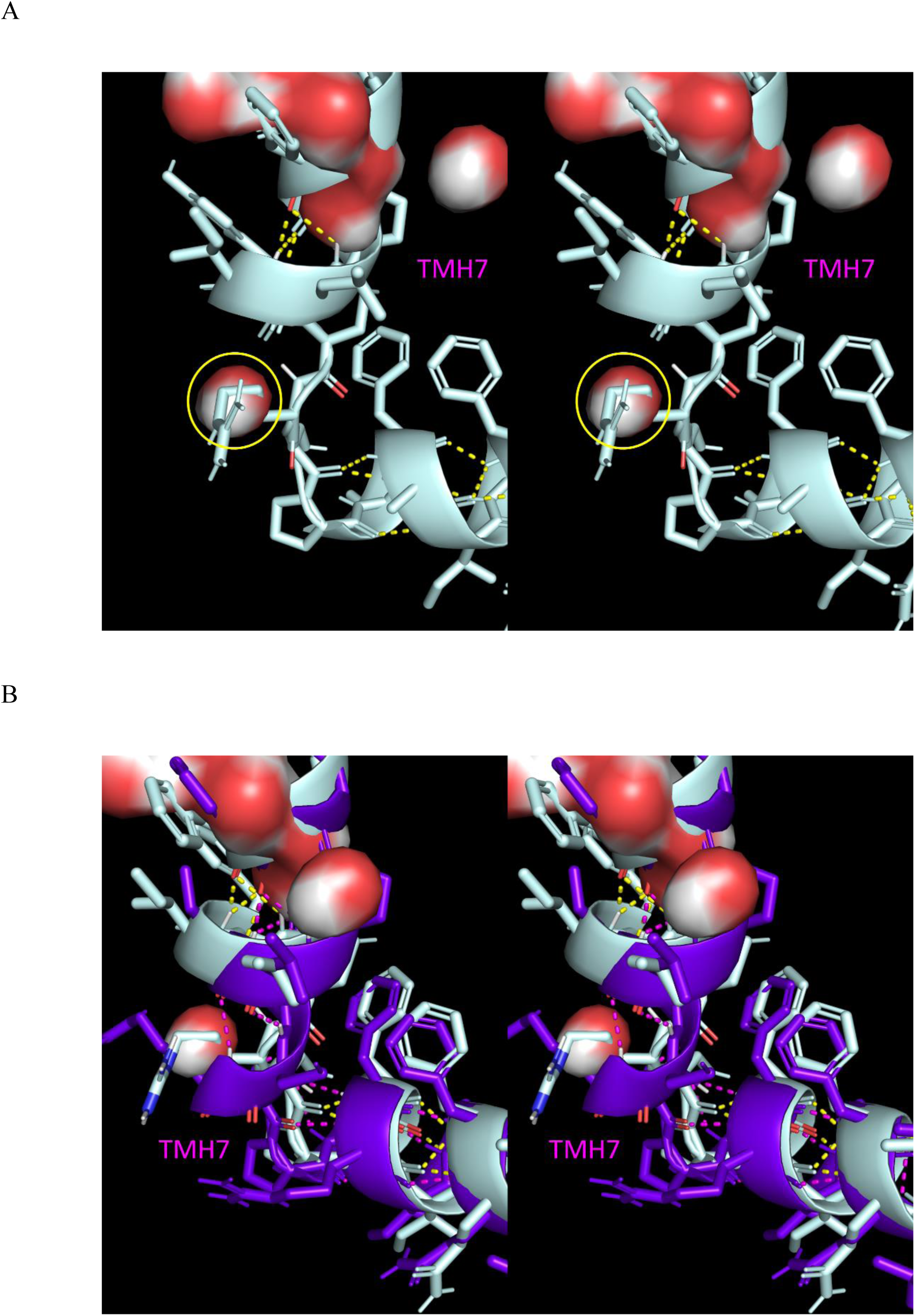
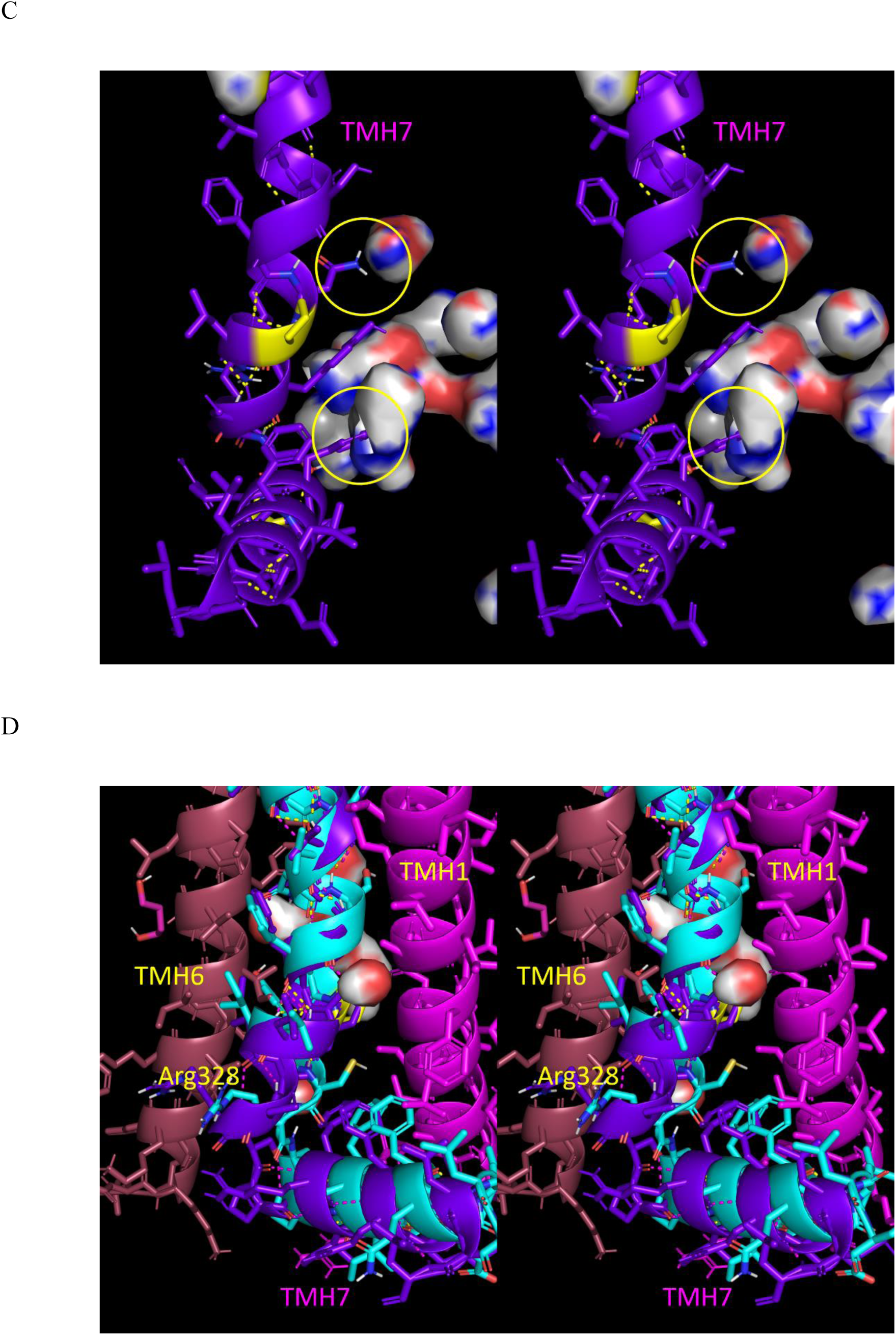
Stereo views of TMH7 showing the contributions to tropimer curvature in the activated state (3SN6) listed in Table 4. (A) The L-shaped tropimer on the IC side of TMH7 in the activated/pivoted position is powered by desolvation of an internal water cavity (yellow circle) by Arg328. (B) Same as A, except with the deactivated TMH7 (purple) overlaid for reference. (C) The L-shaped conformation of TMH7-TMH8 results, in part, from its registry with two water cavities present in the activated state that are desolvated during deactivation. (D) The tropimer in the deactivated/unpivoted position of TMH7 is governed by packing with TMH1 and TMH6. The activated TMH7 (cyan) is overlaid for reference (noting the difference in the Arg328 position).

#### The role of the Asp130-Arg131-Tyr132 (DRY) motif in β_2_-AR activation/deactivation

The aspartate-arginine-tyrosine (DRY) motif residing on the IC end of TMH3 is an oft cited feature in the context of GPCR structure-function. We postulate that Asp130 and Arg131 stabilize the folded state of ICL2 in the deactivated state, which transitions to an extended α-helix-containing structure in the activated state (subserving outward pivoting of TMH4 and transient stabilization in the pivoted state) (Figure 25).

**Figure 25.**
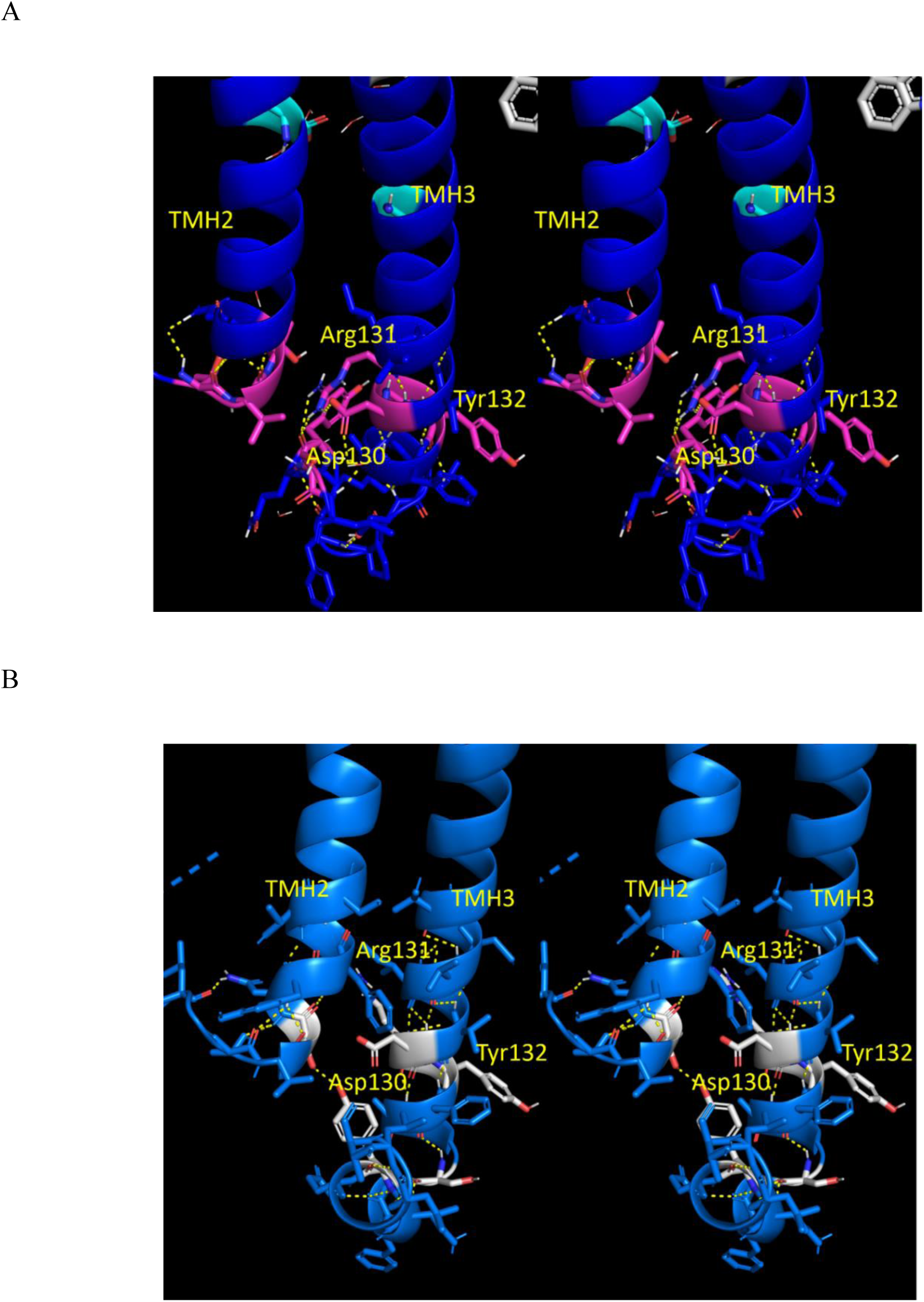
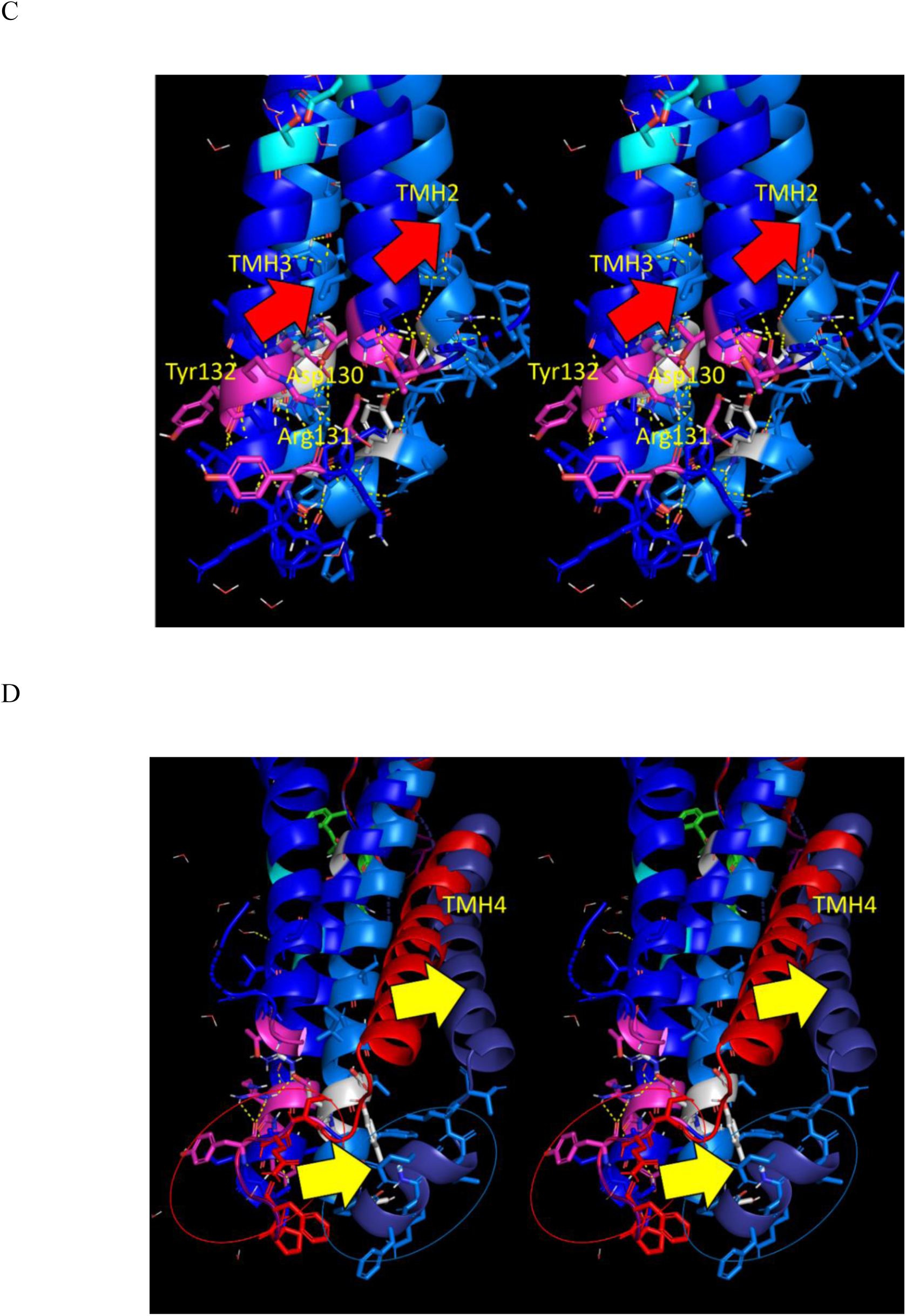
Stereo views of the DRY motif located at the IC end of TMH3 in the β_2_-AR. (A) The deactivated receptor (2RH1) (dark blue with the DRY residues highlighted in magenta). (B) Same as A, except for the activated receptor (DRY residues highlighted in white). (C) Overlay of the activated and deactivated receptor structures (color coding the same as A and B), showing pivoting of TMH2 and TMH3 in the activated structure. (D) Same as C, except with TMH4 included in the deactivated and activated states state (red and purple cartoons, respectively). Pivoting of TMH4 accompanied by the formation of an α-helix in ICL2 is apparent (solid yellow arrows).

#### G-protein binding

As we have demonstrated, the allosteric G-protein binding pocket opens in response to agonist binding to the orthosteric pocket via TMH pivoting and translation, together with tropimer curving within TMH6 and TMH7. Allosteric pocket opening alleviates clashing between the N-terminal helix of the G_α_ subunit of the G-protein with TMH3, TMH4, and TMH6 in the closed allosteric pocket (Figure 26).

**Figure 26.**
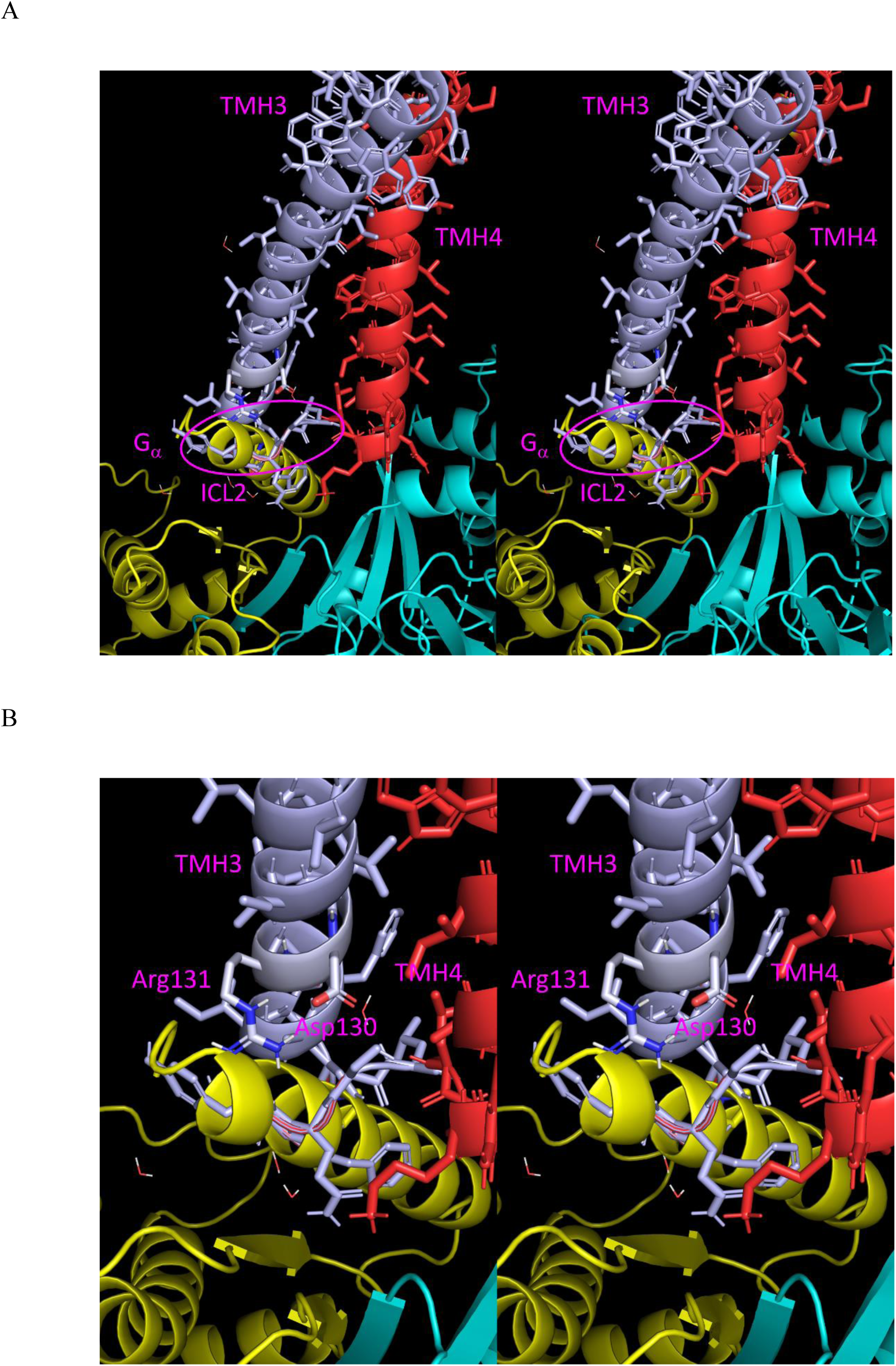
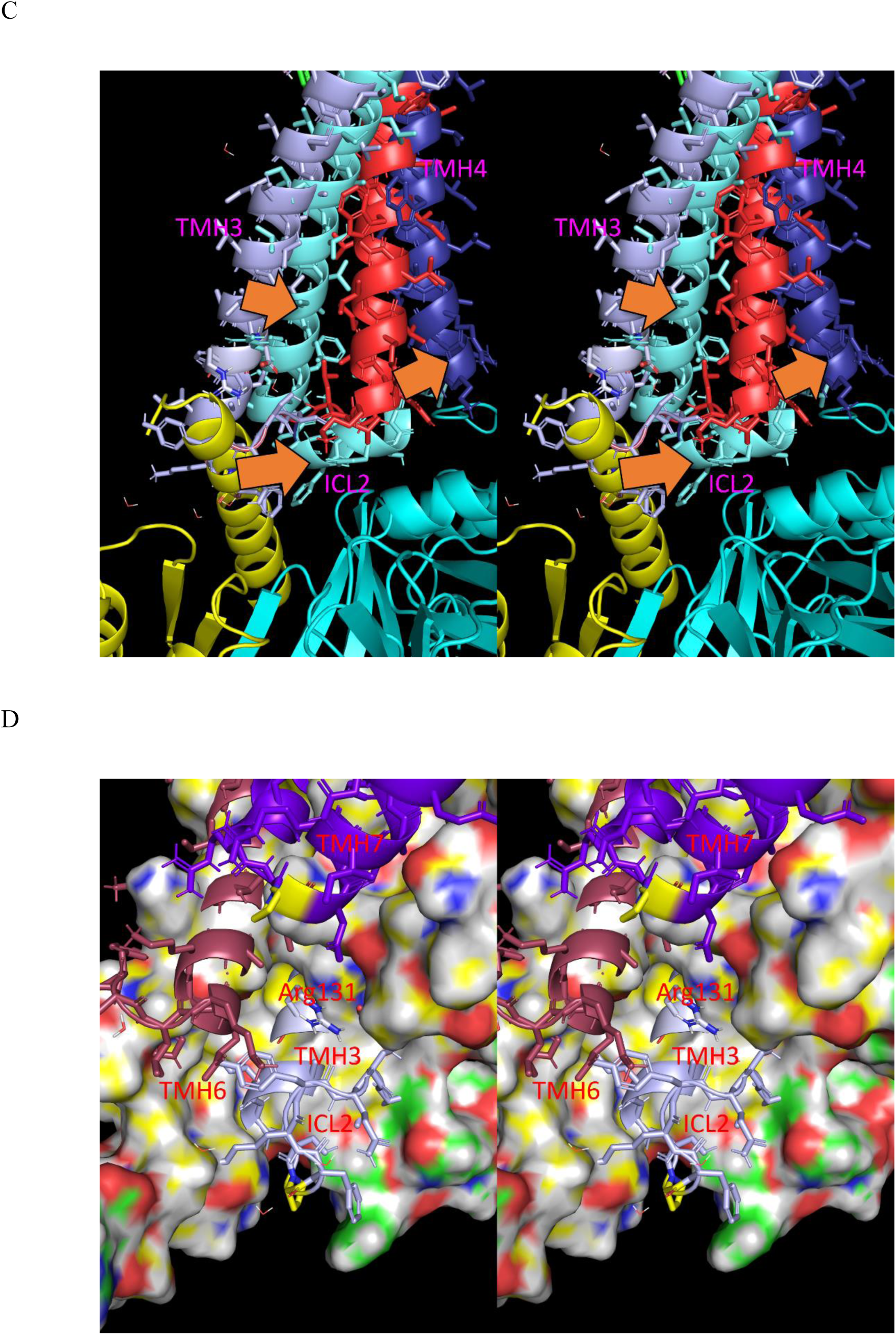
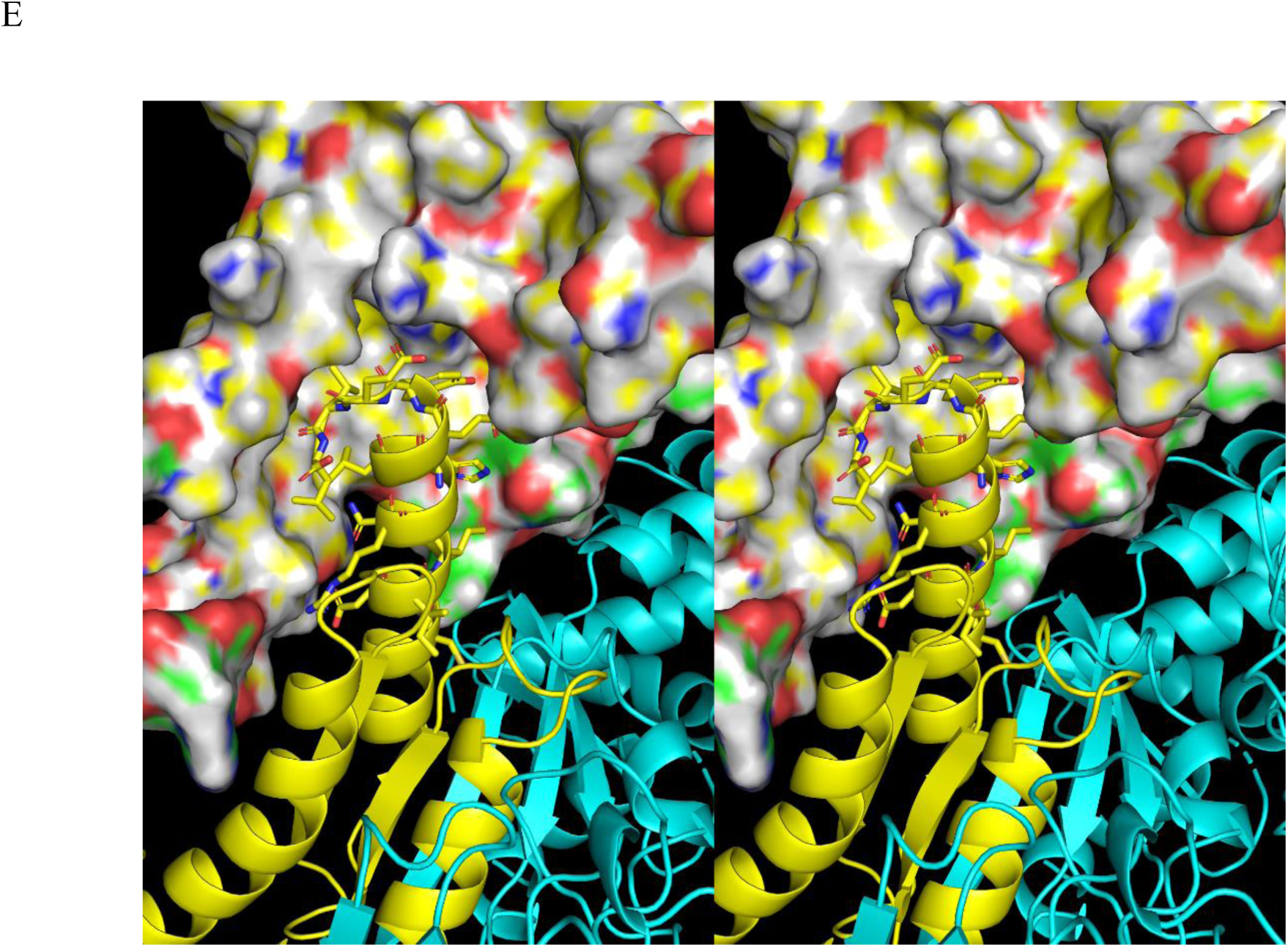
Stereo views of the G-protein interface of the β_2_-AR. (A) G-protein binding is blocked in the deactivated state directly by ICL2. (B) ICL2 is stabilized in its deactivated state by Asp130 and Arg131, which explains the key contribution of these residues to the deactivation mechanism. (C) Pivoting of TMH3 and TMH4 accompanied by rearrangement of ICL2 into an α-helix opens the binding site for the Gα α-helix. (D) Encroachment of TMH3, TMH6, and ICL2 in the G_α_ α-helix pocket located within the deactivated allosteric pocket. (E) The G-protein bound to the activated allosteric pocket, largely by the α-helix of G_α_.

### Putative VGCC structure-function relationships

As for most protein superfamilies, all cationic VGCCs adopt a canonical fold that exhibits significant variation within and between each of the member families. K_v_ channels may consist of homo- or four-fold asymmetric hetero-tetramers, whereas Na_v_ channels consist of four-fold asymmetric single chain hetero-domain proteins, two of which convey VSD-dependent N-type inactivation function. Structural variability among VGCCs is further amplified by differences in loop sizes/behaviors and the frequent presence of cytoplasmic domains and accessory proteins. Each major subunit is comprised of:

1. A voltage-sensing domain (VSD), consisting of four TMHs (S1-S4).
2. One-fourth of the pore domain, consisting of two TMHs (S5 and S6), one diagonally oriented pore helix (P) residing in the extracellular membrane leaflet, one IC S5-S6 loop, one EC S5-pore loop, one EC selectivity filter loop, and one IC S4-S5 linker.
3. Additional domains on a case-by-case basis.

Structural variability within and between the VSD and pore domains and interconnecting loops across the superfamily results in poor structural alignment of the full length proteins, as well as the individual domains thereof. For this reason, apples-to-apples comparison of the activated and deactivated states of VGCCs was limited in this work to Na_v_1.7, for which these states were captured in published cryo-EM structures (6N4Q and 6N4R, respectively)^18^ (Figure 27) (noting that the structures were solved using engineered protein constructs complexed with the spider toxin ProTx2).

**Figure 27.**
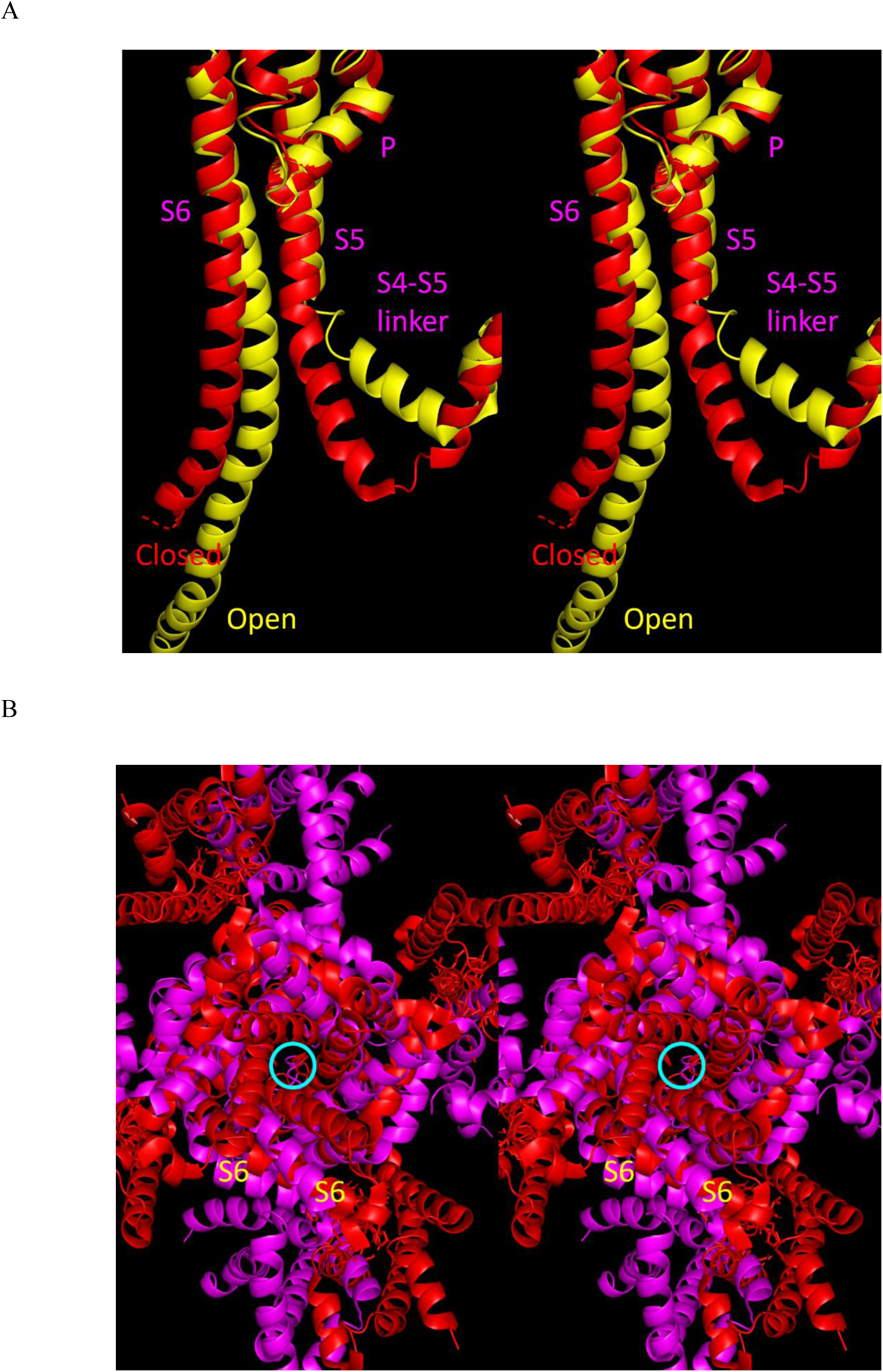
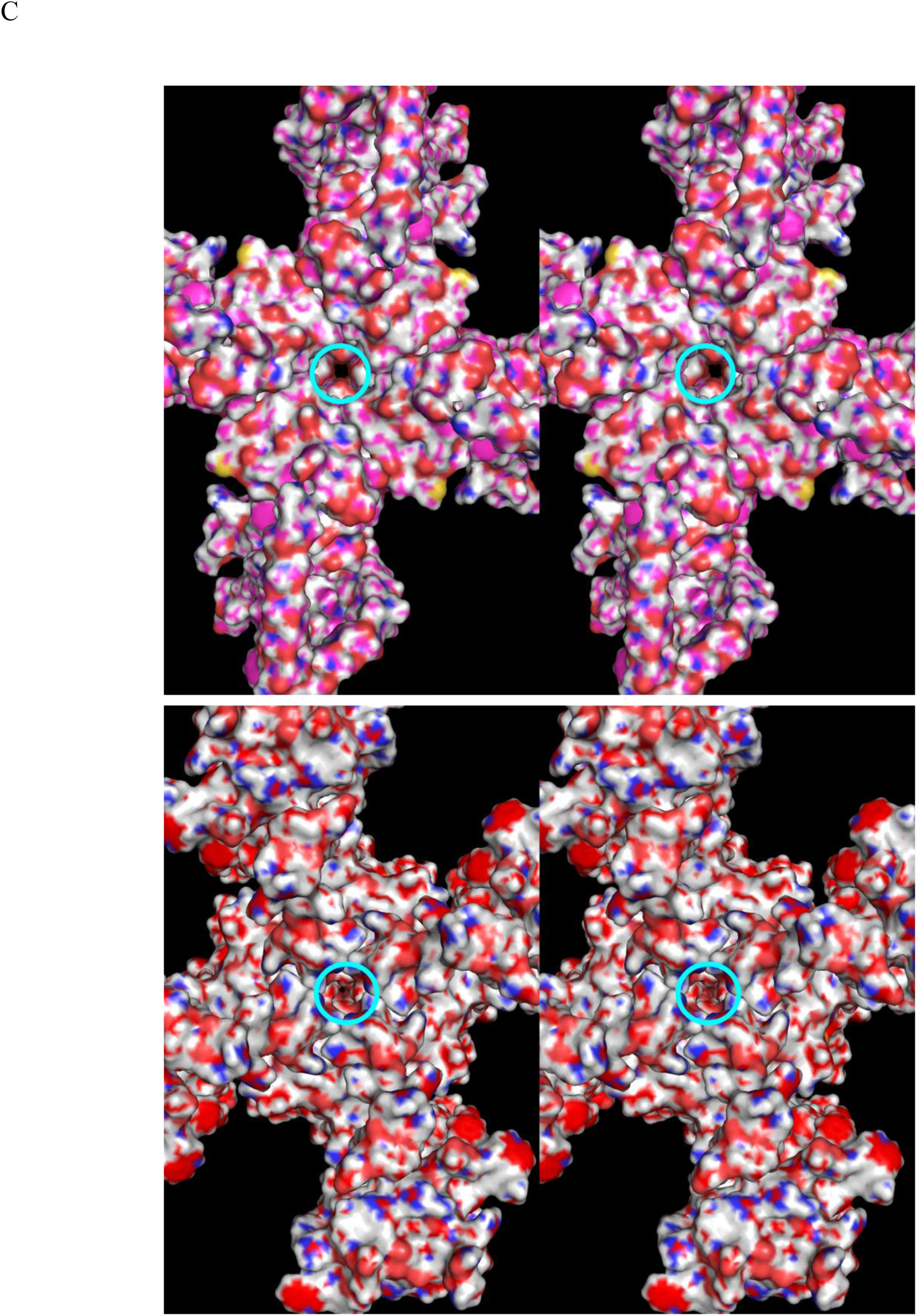
(A) Stereo view of the overlaid pore domains extracted from human the activated/open (yellow cartoon) and deactivated/closed (red cartoon) structures of Na_v_1.7 (6N4Q and 6N4R, respectively). The S4-S5 linker, S5, and S6 shift toward and away from the central pore axis in the closed versus open states, respectively. (B) Stereo view of the overlaid full length open K_v_1.2 (2A79) and closed KCNQ1 (5VMS) structures (the central pore axis is shown as a cyan circle). The relative positions of S6 in the activated and deactivated states are apparent. (C) Same as B, except with the surfaces shown (noting full closure of the KCNQ1 structure).

#### Activation and deactivation of VGCCs are initiated by changes in membrane voltage and membrane dipole potential

We set about to understand the mechanism by which the membrane and peri-channel dipole potentials (Δψ_*m*_(*t*) and Δψ_*d*_(*t*), respectively) are transduced into the activated and deactivated structural states of VGCCs (noting that VGCC <u>inactivation</u> mechanisms are beyond the scope of this work). As determined from the first published VGCC crystal structure, VGCCs are physically blocked (i.e., “gated”) and unblocked by the S6 TMH, the state transitions of which are governed indirectly by the S5 TMH.^7^ In our previous work on the general mechanism of VGCC gating,^43^ we postulated that Δψ_*m*_(*t*) and Δψ_*d*_(*t*) operate in tandem to drive channel activation and deactivation, as follows (noting that activation and deactivation driven solely by Δψ_*m*_(*t*) cannot be ruled out):

1. The channels are closed/deactivated in the resting state, wherein:

a. The pore domain is fully elongated and the peri-channel membrane thickeness increases via lipid chain extension to fully encapsulate the non-polar regions of the protein (thereby circumventing solvent exposure of those regions).
b. Δψ_*d*_(*t*) is maximally unfavorable due to increased ordering of the extended lipid chains (i.e., the potential energy transduced into channel activation is stored within the membrane itself).
2. Relaxation of Δψ_*d*_(*t*) via lipid chain disordering, accompanied by contraction of the pore domain and peri-channel membrane thinning, is opposed by Δψ_*m*_(*t*) (analogous to the brake in a car), thereby circumventing spontaneous channel activation in the resting state (noting that leak currents depend on constitutively active VGCCs).
3. The “brake” is released during depolarization (i.e., reversal of Δψ_*m*_(*t*)), resulting in spontaneous relaxation of Δψ_*d*_(*t*) via channel activation-mediated thinning of the peri-channel membrane due to lipid chain disordering in the contracted form of the pore domain (analogous to the accelerator in a car). Channel activation proceeds from a running start in the “accelerator-brake” model versus from a standstill in the Δψ_*m*_(*t*) model.
4. Channel deactivation and resetting of the high energy state of Δψ_*d*_(*t*) are driven by the reversal of Δψ_*m*_(*t*) toward the repolarized state (serving as the “accelerator” and later the “brake” in the fully repolarized state) (noting that VGCC currents are initially switched off via rapid activation gate-independent N- and C-type inactivation mechanisms).

#### VSD state transitions are powered principally by desolvation of internal cavities

The “brake” on the deactivated → activated state transition and the “accelerator” powering the deactivated → activated state transition (and the activated → inactivated transition in some Na_v_ channels) are situated in the VSDs. We probed the activation/deactivation mechanism using two structures of the activated and deactivated states of Na_v_1.7 (6N4Q and 6N4R, respectively)^18^, which we overlaid about S1 and S2 of the VSD (Figure 28A). S4 contains four Arg residues, the side chains of which face approximately toward the S4-S1 interface (Figure 28B). The positions, types, and numbers of basic S4 residues vary across the VGCC superfamily, and activation is widely attributed to interactions between Δψ_*m*_(*t*) and these residues (which are commonly referred to as “gating charges”).^8^ The charged side chains are widely assumed to translocate with the negative wavefront of Δψ_*m*_(*t*) during the depolarization and repolarization phases of the action potential.

**Figure 28.**
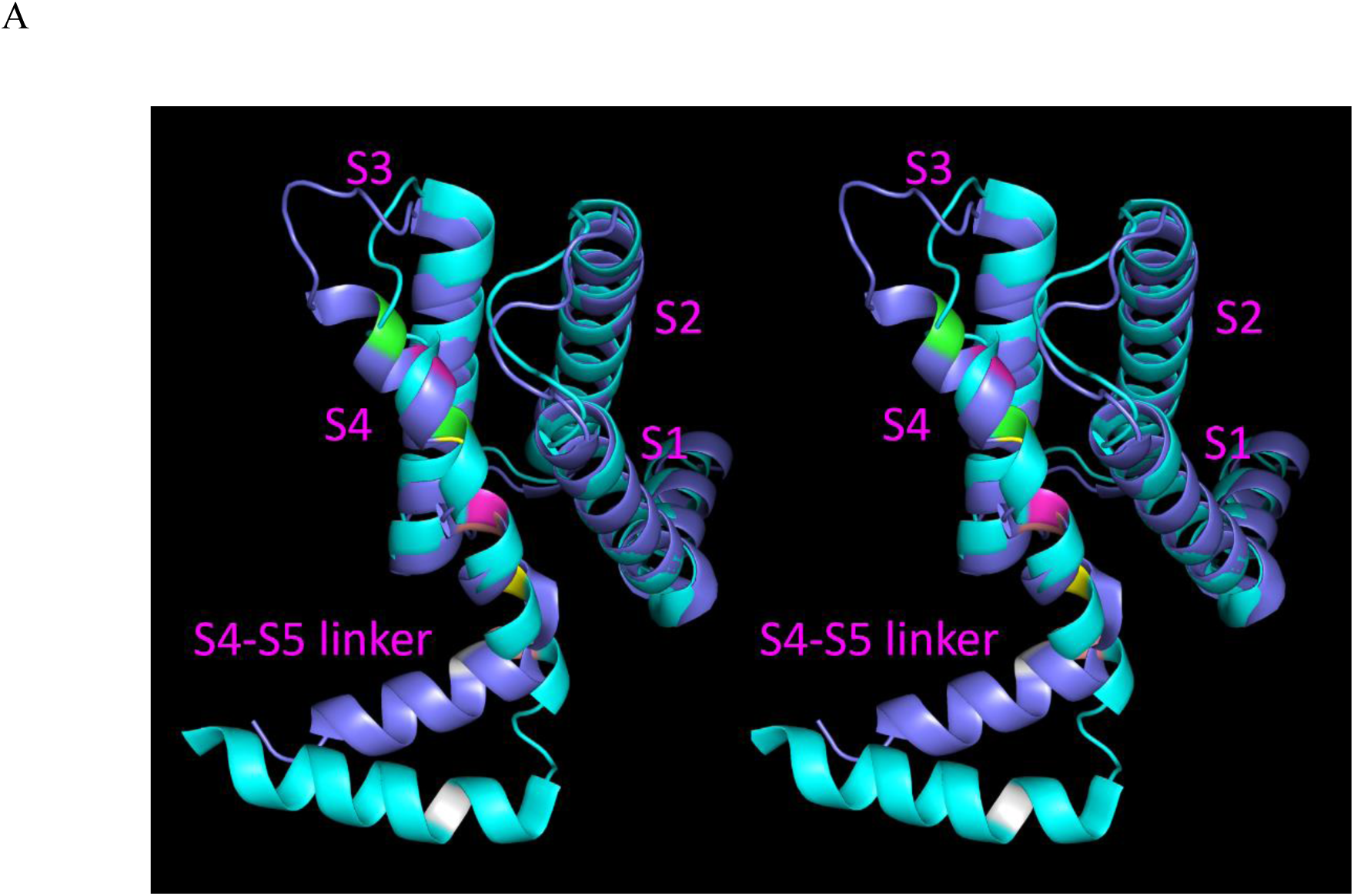
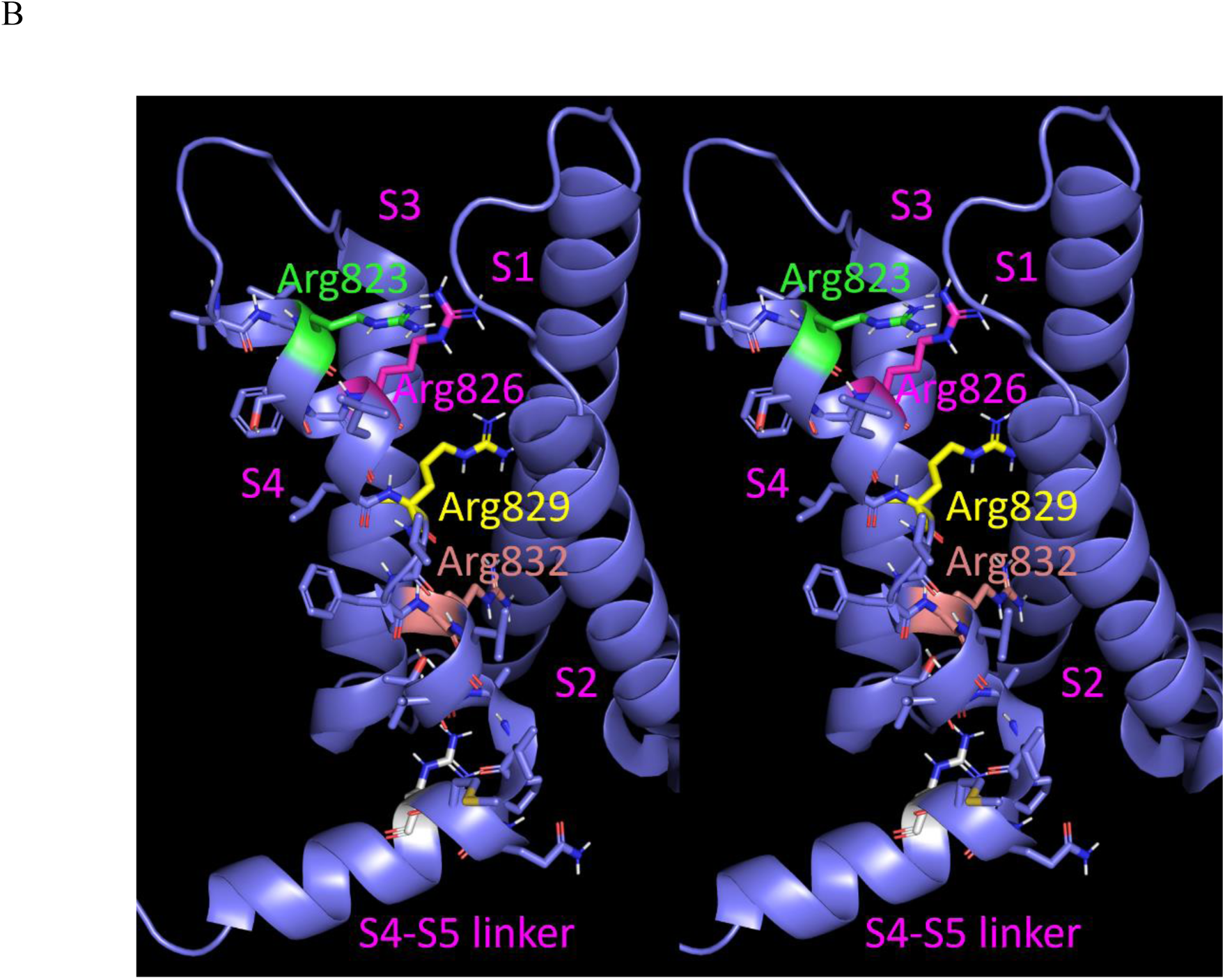
(A) Stereo view of the VSD of the activated (blue cartoon) and deactivated (cyan cartoon) Na_v_1.7 structures (6N4Q and 6N4R, respectively) overlaid about S1 and S2 looking approximately orthogonal to S3. (B) Same as A, except showing the four Arg residues on S4.

However, the Na_v_1.7 structures suggest a very different mechanism, in which S4 undergoes ∼10 Å semi-rigid-body oscillatory translations in the EC and IC directions during activation and deactivation, respectively^18^ (Figure 29).

**Figure 29.**
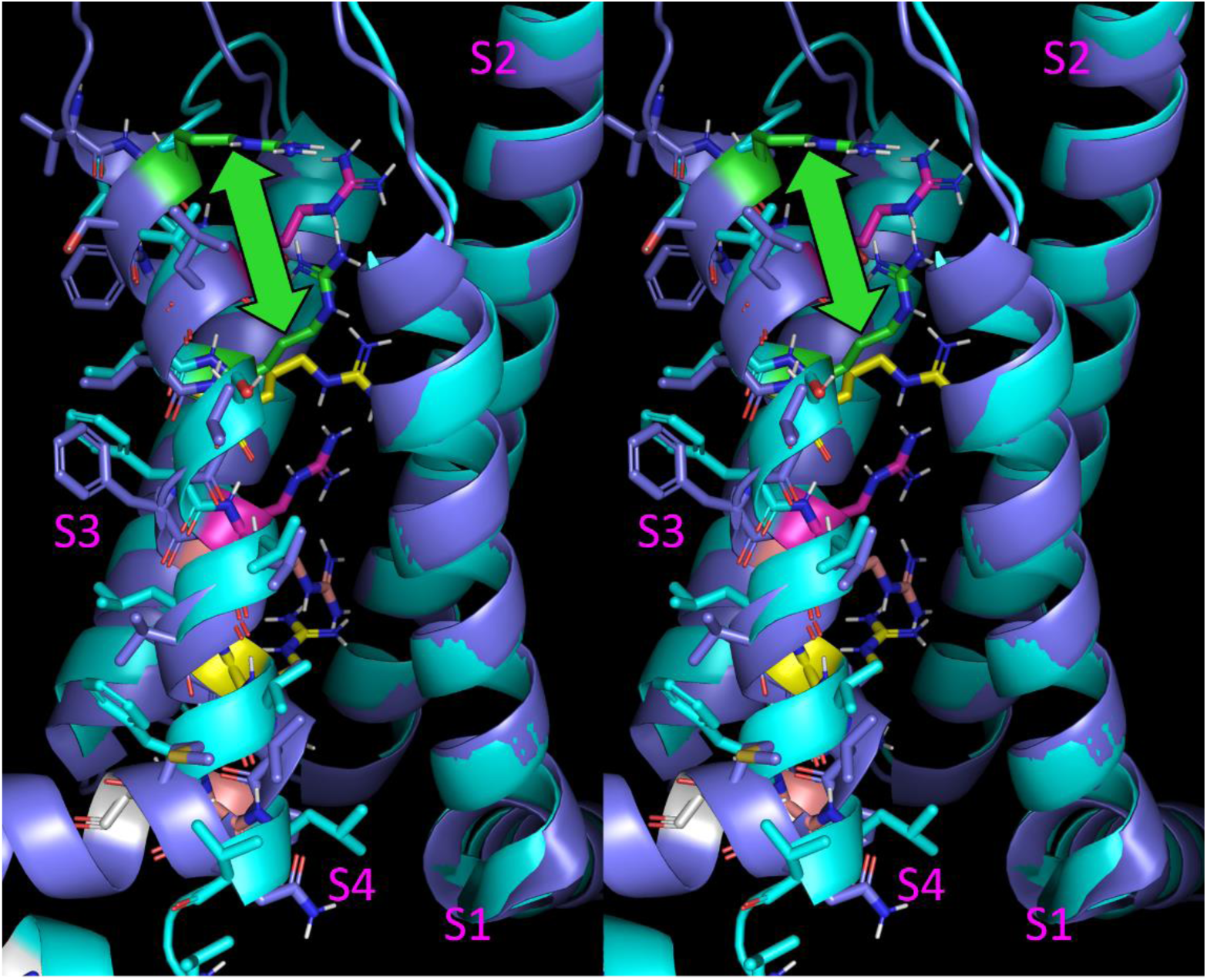
Stereo view of the semi-rigid-body translation of S4 (green double-headed arrow) observed in 6N4Q (blue cartoon) relative to 6N4R (cyan cartoon). The corresponding Arg side chains are color-coded the same in the two structures (numbering provided in Figure 28B).

That the observed translation of S4 is powered solely by electrical interactions between Δψ_*m*_(*t*) and the Arg side chains is unlikely, due to the fact that the positive charges on these residues are heavily screened by solvation, and the desolvation cost of these residues is high (ruling out the possibility that they reside within non-polar environments). Instead, we postulate that:

1. The Arg side chains are well-solvated by internal solvation within the VSD.
2. The configuration of this solvation is dynamically altered by Δψ_*m*_(*t*) (noting that the activated and deactivated states are captured in the structures, despite the absence of this potential and Δψ_*d*_(*t*) in the cryo-EM preps).
3. The Arg side chains (the gating charges) follow the solvation configuration via the observed S4 translations, which serves as both the free volume and power source for the rearrangements, analogous to the proposed rigid-body pivoting described above for the β_2_-AR.

We set about to test this hypothesis by analyzing the internal cavities within the activated and deactivated VSD states captured in 6N4R and 6N4Q, respectively (Figure 30).

**Figure 30.**
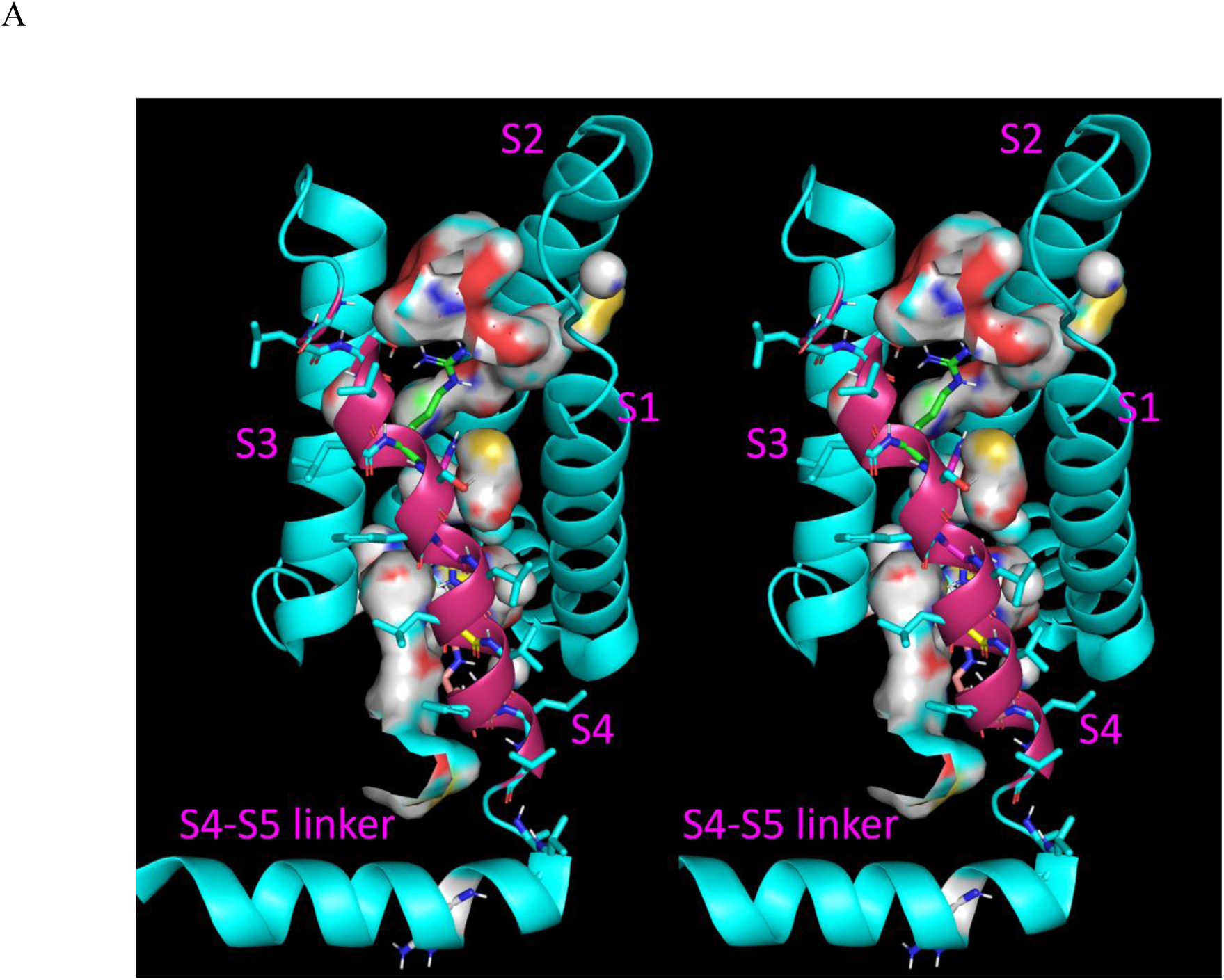
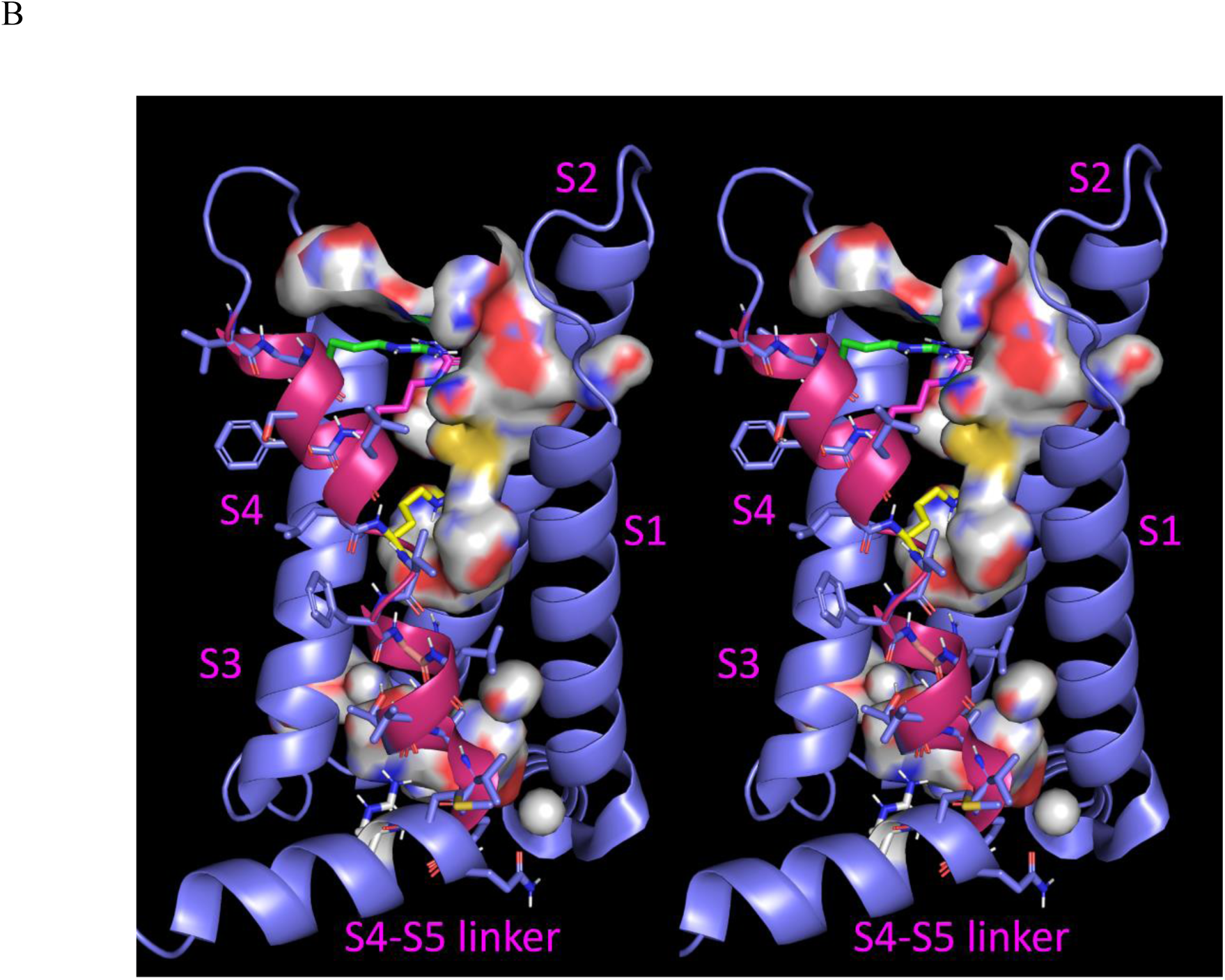
Stereo views of the Na_v_1.7 VSD looking toward S4 (warm pink), showing the internal cavities within the bundle of (A) the deactivated state (6N4R) and (B) the activated state (6N4Q). S4 is isolated from the other TMHs of the VS and pore domains by internal solvation that provides both the power and free volume for its oscillatory translations.

We postulate that the internal solvation proximal to the S4 Arg side chains is promoted to a large degree by these side chains (Figure 31).

**Figure 31.**
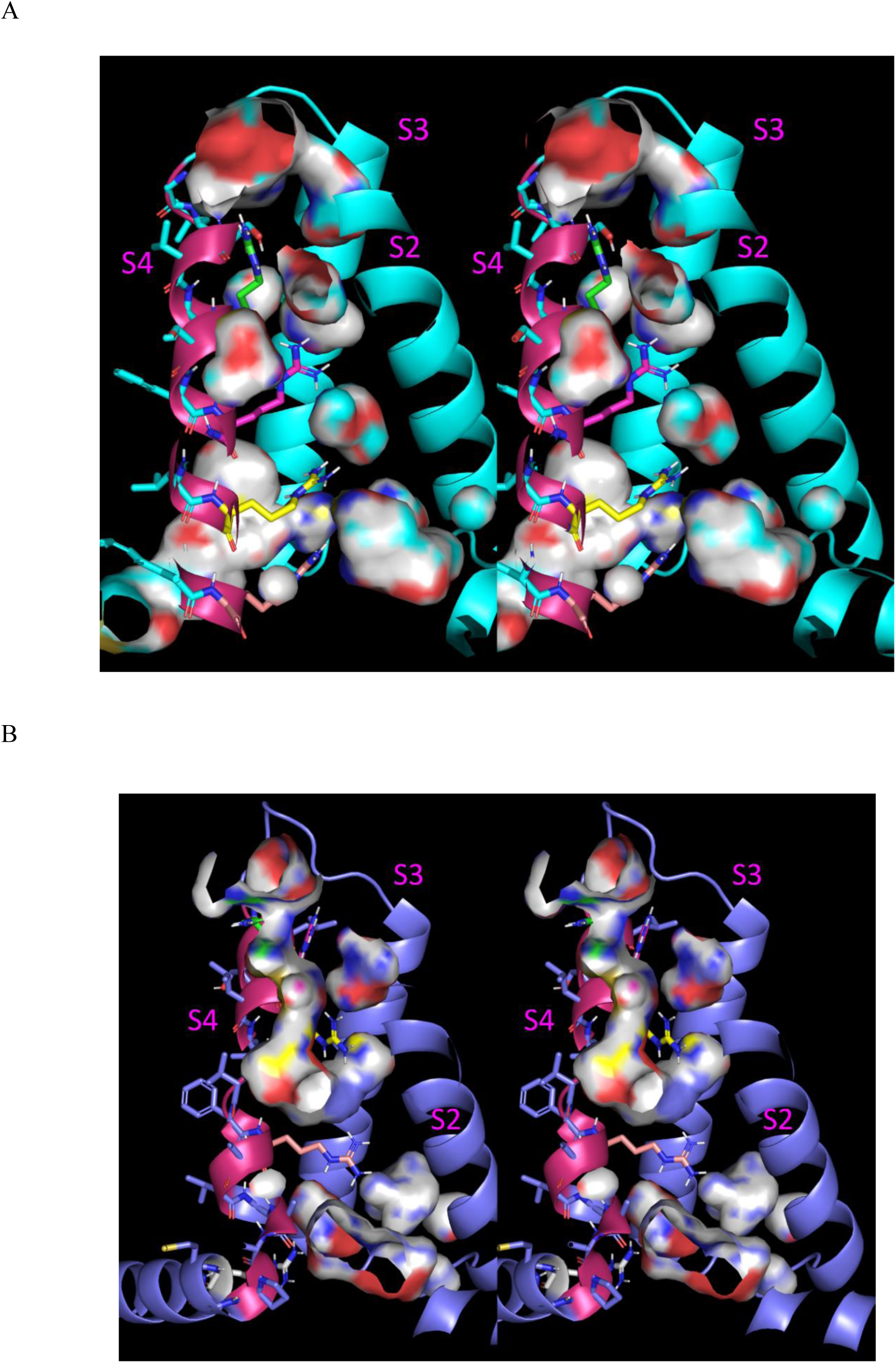
Cutaway stereo views of the close juxtaposition of the Arg side chains on S4 and the internal cavities within the VSD of Na_v_1.7. The guanidinium groups are necessarily H-bonded to the solvation contained within these cavities, which are never desolvated during the S4 translation. (A) The deactivated VSD (6N4R). (B) The activated VSD (6N4Q).

Furthermore, internal cavities present in the activated state are occupied by three Arg side chains in the deactivated state, and internal cavties present in the deactivated state are occupied by all four side chains in the activated state (Figures 32 and 33). These results suggest that bidirectional semi-rigid-body translation of S4 is powered directly by the expulsion of H-bond depleted solvation from the internal cavities of the VSD, rather than Δψ_*m*_(*t*) (which is, in fact, absent in the cryo-EM structure preps). We postulate that Δψ_*m*_(*t*) reversal jumpstarts the observed S4 rearrangement, analogous to the role of agonists in GPCR activation. We further postulate that Δψ_*m*_(*t*) acts on the internal solvation (noting that water is known to be polarizable in the presence of an electric field) rather than the positive charges of the Arg side chains per se, which are heavily screened by solvating water and acidic side chains in some cases (noting that the latter possibility cannot be ruled out).

**Figure 32.**
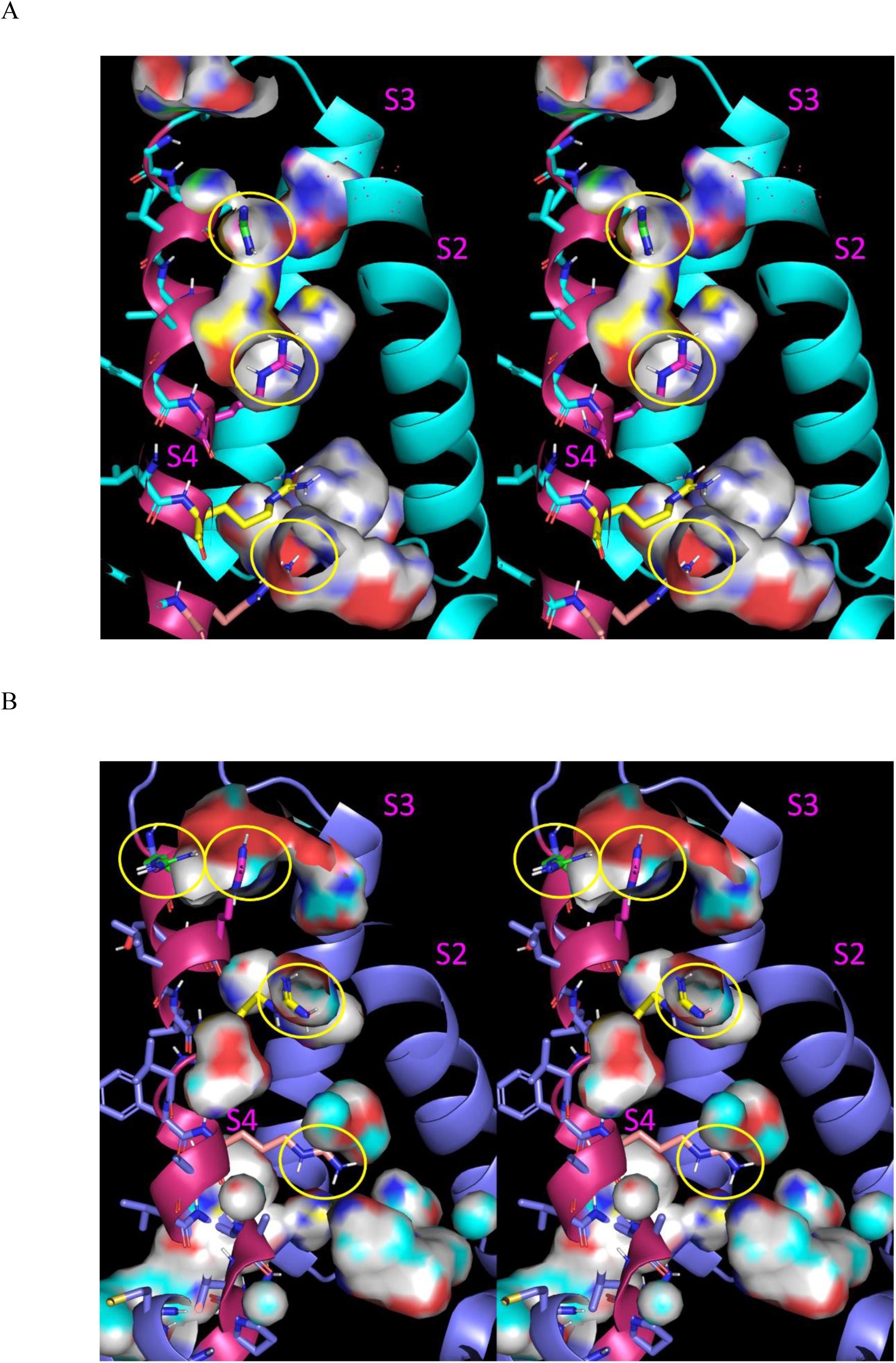
Cutaway stereo views showing the cross-registration between the Arg side chains of S4 in Na_v_1.7 and the internal cavities present in the deactivated and activated states of the VSD. The four Arg positions are circled in yellow. (A) The deactivated state overlaid on the internal cavities present in the activated state. (B) The activated state overlaid on the internal cavities present in the deactivated state.

**Figure 33.**
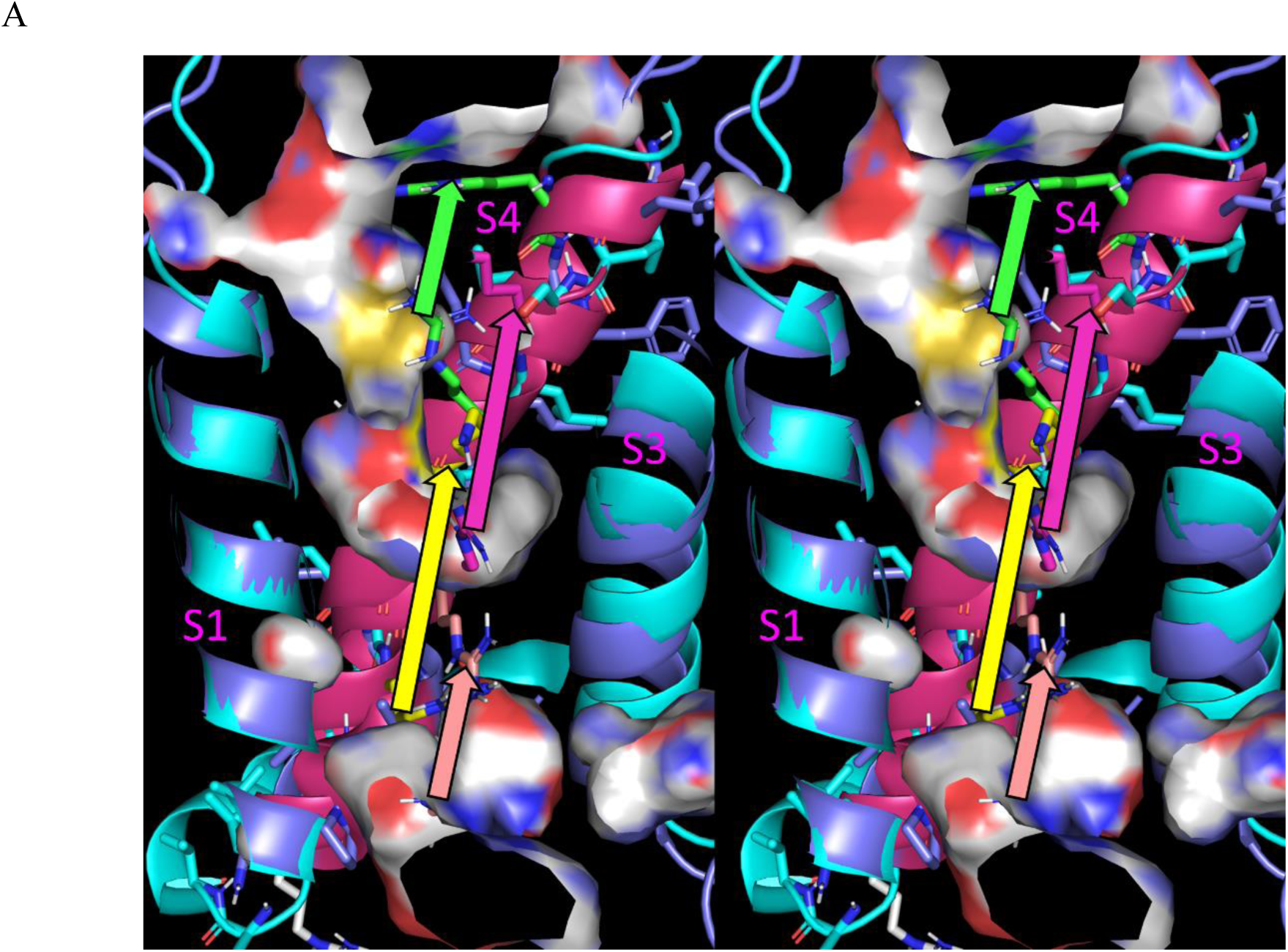
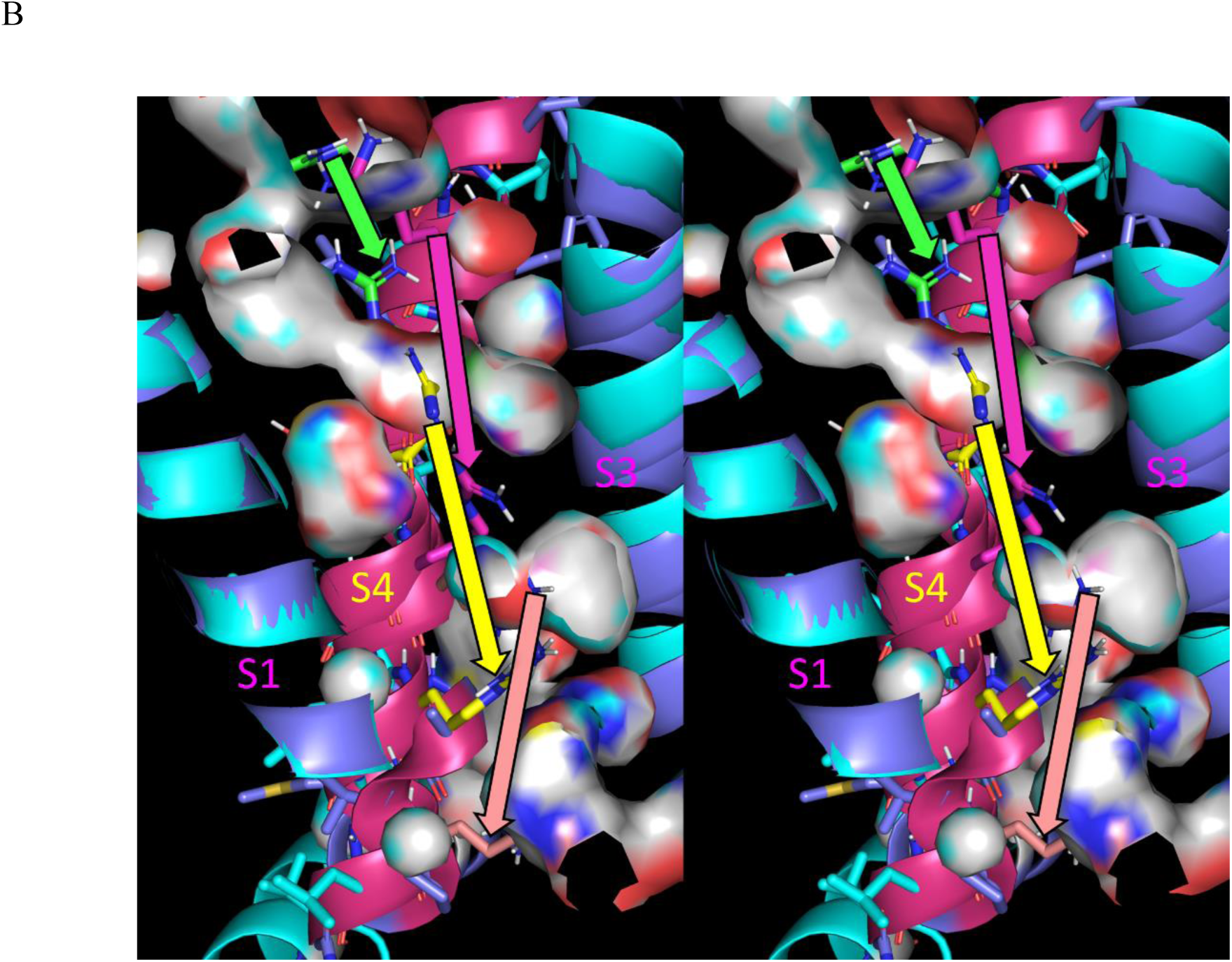
Cutaway stereo views showing bidirectional repositioning of the Arg side chains of S4 into the internal cavities of the opposite overlaid state resulting from semi-rigid-body translations of this TMH orthogonal to the membrane surface in the IC → EC (activation) and EC → IC (deactivation) directions (solid arrows color-coded by the Arg color). We postulate that translation of S4 is powered principally by the expulsion of H-bond depleted solvation from internal cavities by these side chains. (A) The position of S4 in the <u>activated</u> state of the VSD. The Arg side chains occupy internal cavities present in the activated state, except for Arg829 (yellow). (B) The S4 position in the <u>deactivated</u> state of the VSD. All of the Arg side chains, including Arg829, translate into internal cavities present in the deactivated state.

Our proposed VSD activation mechanism is summarized here and in Figure 34:

1. Each VSD cycles between two structural states (VSD_pol_ and VSD_depol_) in the polarized versus depolarized states of Δψ_*m*_(*t*), respectively.
2. As for GPCRs, VSD rearrangements are powered principally by the alternate desolvation and resolvation of two high energy internal solvation configurations in VSD_pol_ and VSD_depol_, corresponding to the braked and unbraked positions of the S4-S5 linker, respectively (noting that the pore domain likewise rearranges between two high energy internal solvation configurations described below). Such rearrangements are kickstarted by Δψ_*m*_(*t*) reversal.
3. The relative free energies of the two solvation configurations 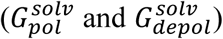 depends solely on Δψ_*m*_(*t*), as follows:

a. In the deactivated/polarized state, the effect of Δψ_*d*_(*t*) acting directly on the pore domain is equal and opposite to that of Δψ_*m*_(*t*) = −90 mV acting directly on the VSDs, which corresponds to the braked position of the S4-S5 linker and 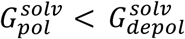.
b. In the activated/depolarized state, the effect of Δψ_*d*_(*t*) on the pore domain is greater than the effect of Δψ_*m*_(*t*) = 110 mV on the VSDs, which corresponds to the unbraked position of the S4-S5 linker and 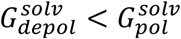.
4. Δψ_*m*_(*t*) may drive the VSD_pol_ ↔VSD_depol_ state transition via direct interaction with the charged side chains on S4 or via indirect polarization of the internal solvation.

**Figure 34.**
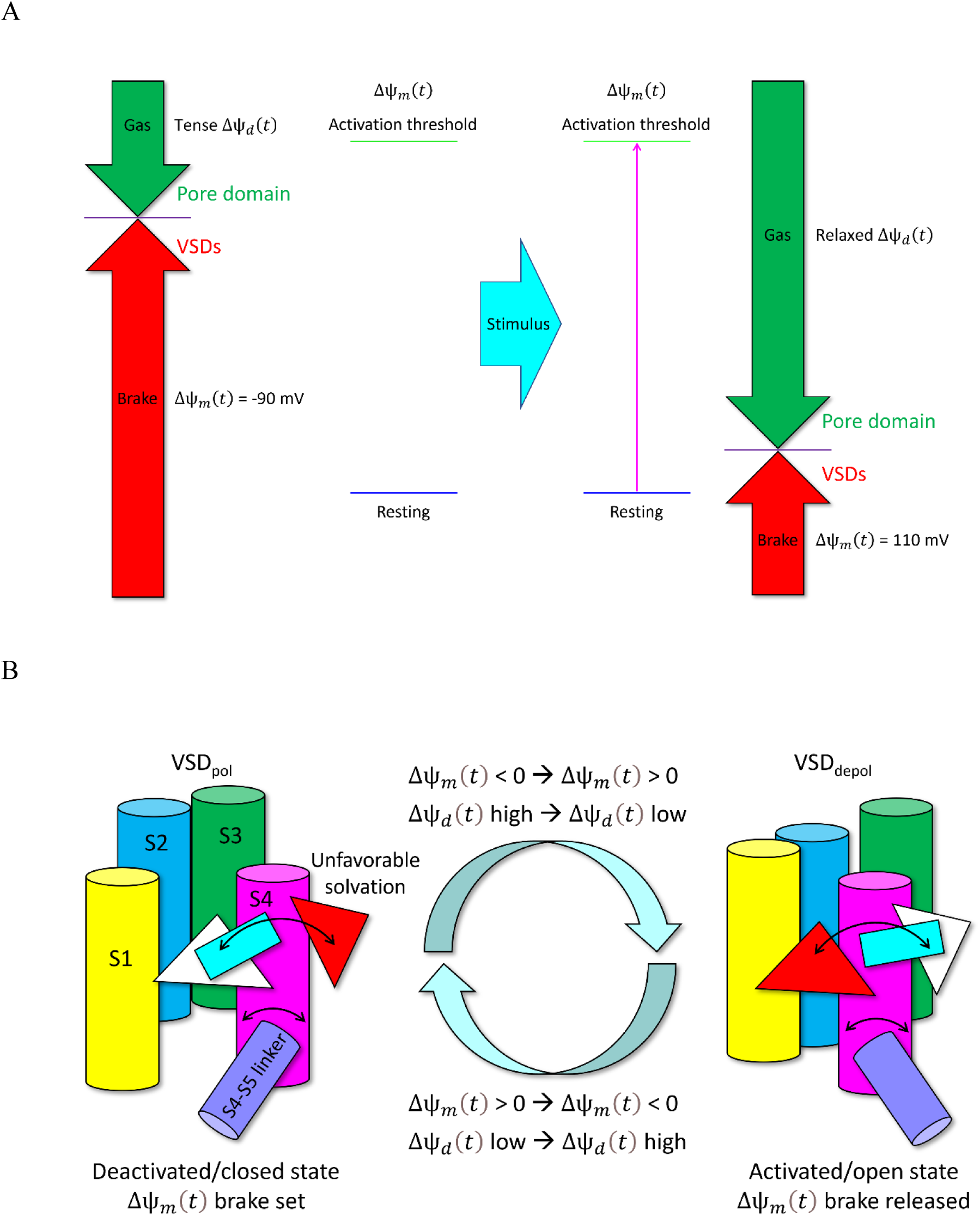
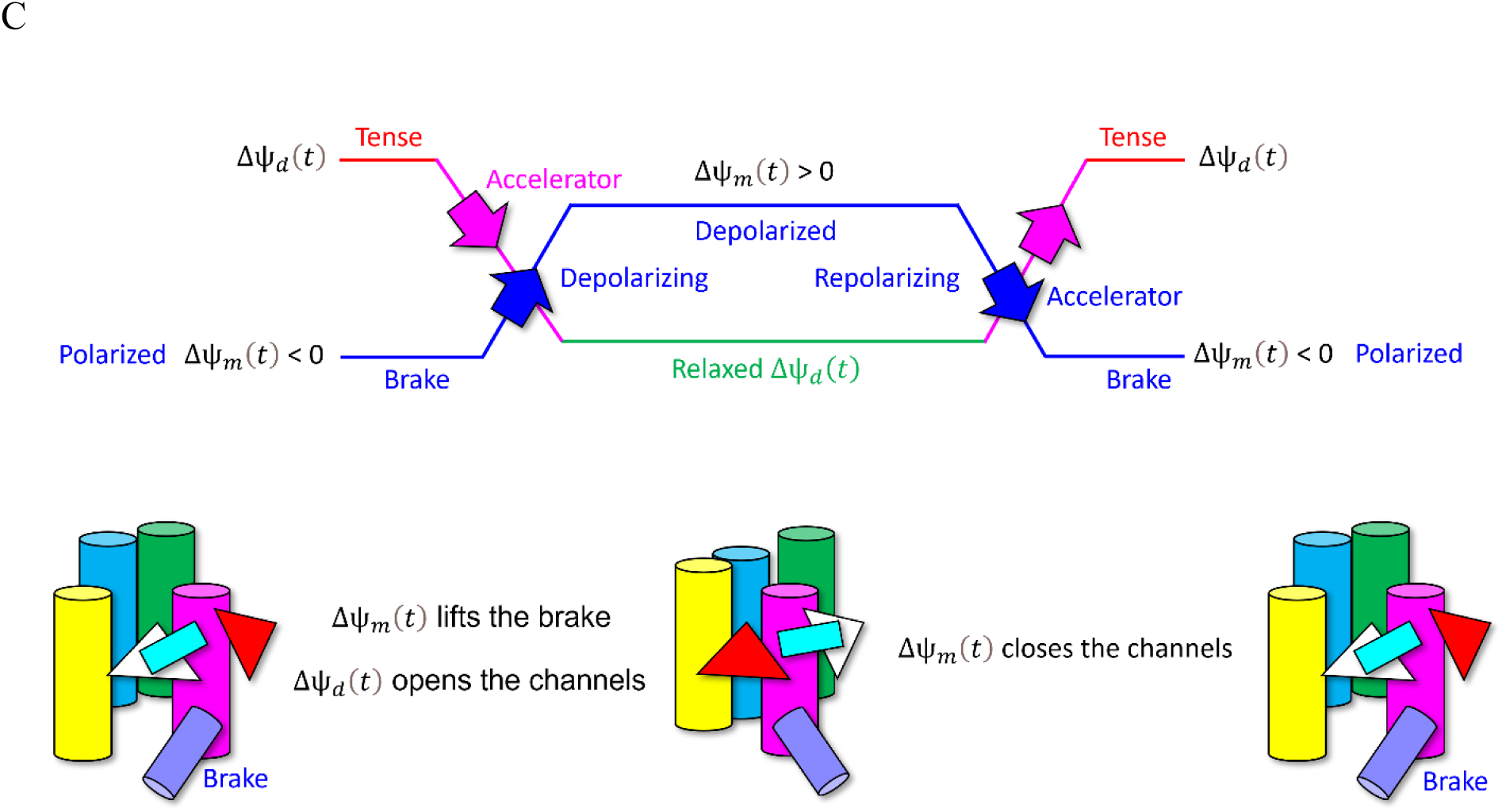
(A) The putative “accelerator-brake” relationship between Δψ_*d*_(*t*) (acting directly on the pore domain) and Δψ_*m*_(*t*) (acting directly on the VSDs) in the resting/polarized state. This relationship reverses during repolarization, where Δψ_*m*_(*t*) serves as the accelerator and Δψ_*d*_(*t*) as the brake. (B) Proposed schema of the three-way interactions between the electric field (Δψ_*m*_(*t*)), dipole potential (Δψ_*m*_(*t*)), and solvation fields within the VSD. Δψ_*m*_(*t*) opposes Δψ_*d*_(*t*) (i.e., acting as the brake) indirectly via its effect on the solvation field within the VSD corresponding to the “braked” position of the S4-S5 linker. The brake is released by Δψ_*m*_(*t*) reversal (i.e., depolarization) in which the Arg-water interactions powering VSD_depol_ are disrupted, resulting in channel opening via Δψ_*d*_(*t*) -powered relaxation (the “accelerator”). Δψ_*m*_(*t*) re-reversal (i.e., repolarization) stabilizes the solvation field within the VSD corresponding to the braked position of the S4-S5 linker (i.e., VSD_pol_), which in turn, restores the braked position of the S4-S5 linker. (C) Same as B, except showing the mapping between the VSD structural states and Δψ_*m*_(*t*) and Δψ_*d*_(*t*).

#### Connecting the dots between VSD and pore domain rearrangements

Next, we set about to understand how S4 repositioning is relayed to the S4-S5 linker and on to S5 and S6 of the pore domain (the latter of which directly opens and closes the ion conduction pathway). An internal cavity present at the base of S4 in the deactivated VSD structure (6N4R) coincides with the S4-S5 linker position in the activated VSD structure (Figure 35), suggesting that this cavity is a key endpoint of S4 translation. This, in turn, promotes spontaneous repositioning of the S4-S5 linker therein (noting that this latter step entails rearrangements within the elbow region between S4 and the linker, analogous to those in elbow region between TMH7 and TMH8 in GPCRs).

**Figure 35.**
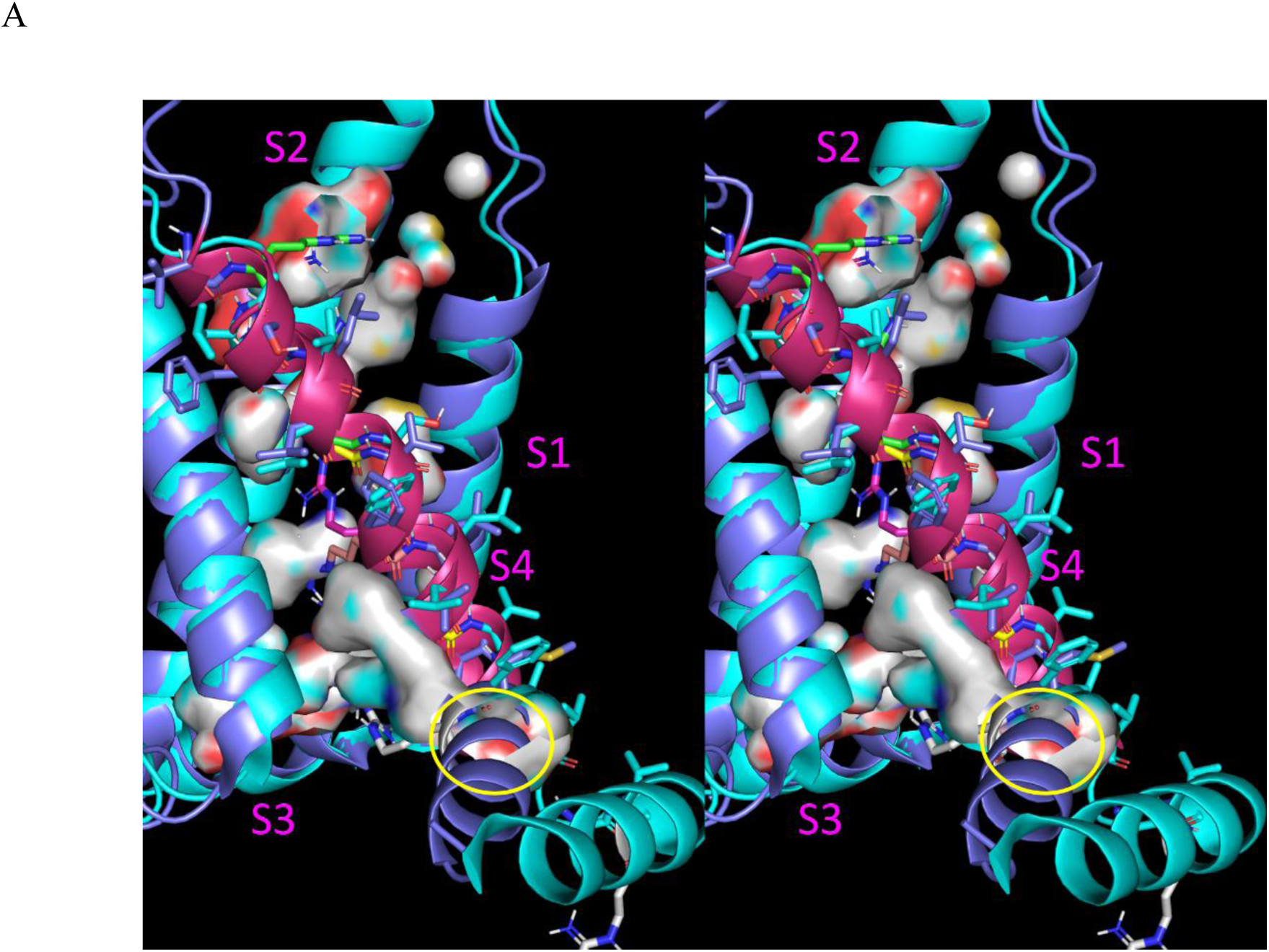
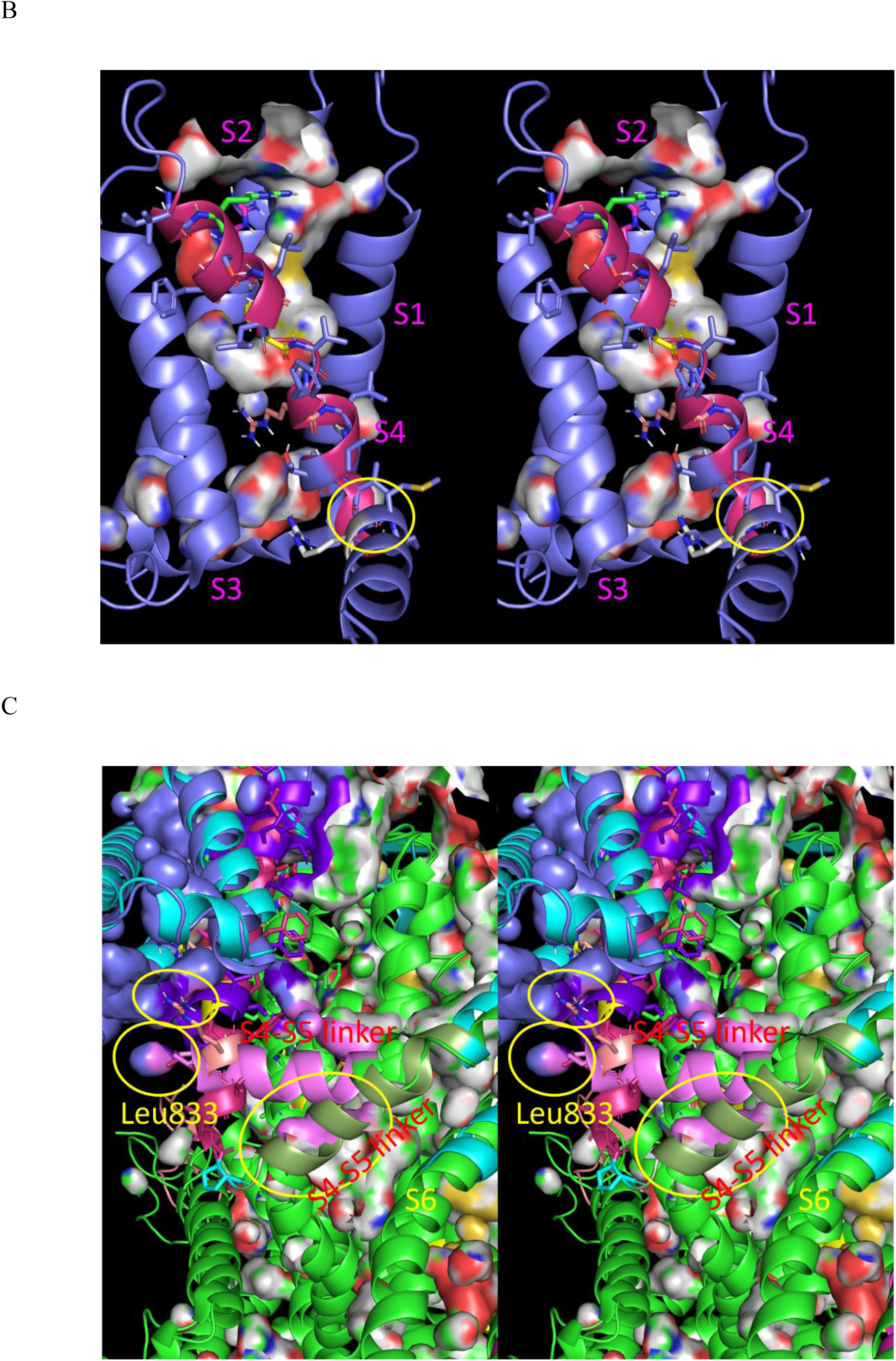
Stereo views of the VSD of Na_v_1.7. (A) Overlay of the activated (blue cartoon) and deactivated (cyan cartoon) states of the VSD. An internal cavity (yellow circle) present in the deactivated state powers repositioning of the S4-S5 linker in the activated state. (B), Same as A, except showing the surface in the activated state, in which this cavity is is fully absent. (C) Expanded view of the VSD and pore domain, including the S4-S5 linker in its deactivated (olive green) and activated (pink) positions. Leu833, Arg832, and the S4-S5 linker of the deactivated structure occupy internal cavities present in the activated structure (yellow circles).

Additional internal cavities residing between the VSD and pore domain, as well between as adjacent subunits of the tetramer, are likewise occupied by the S5 and S6 TMHs (Figures 35-38).

**Figure 36.**
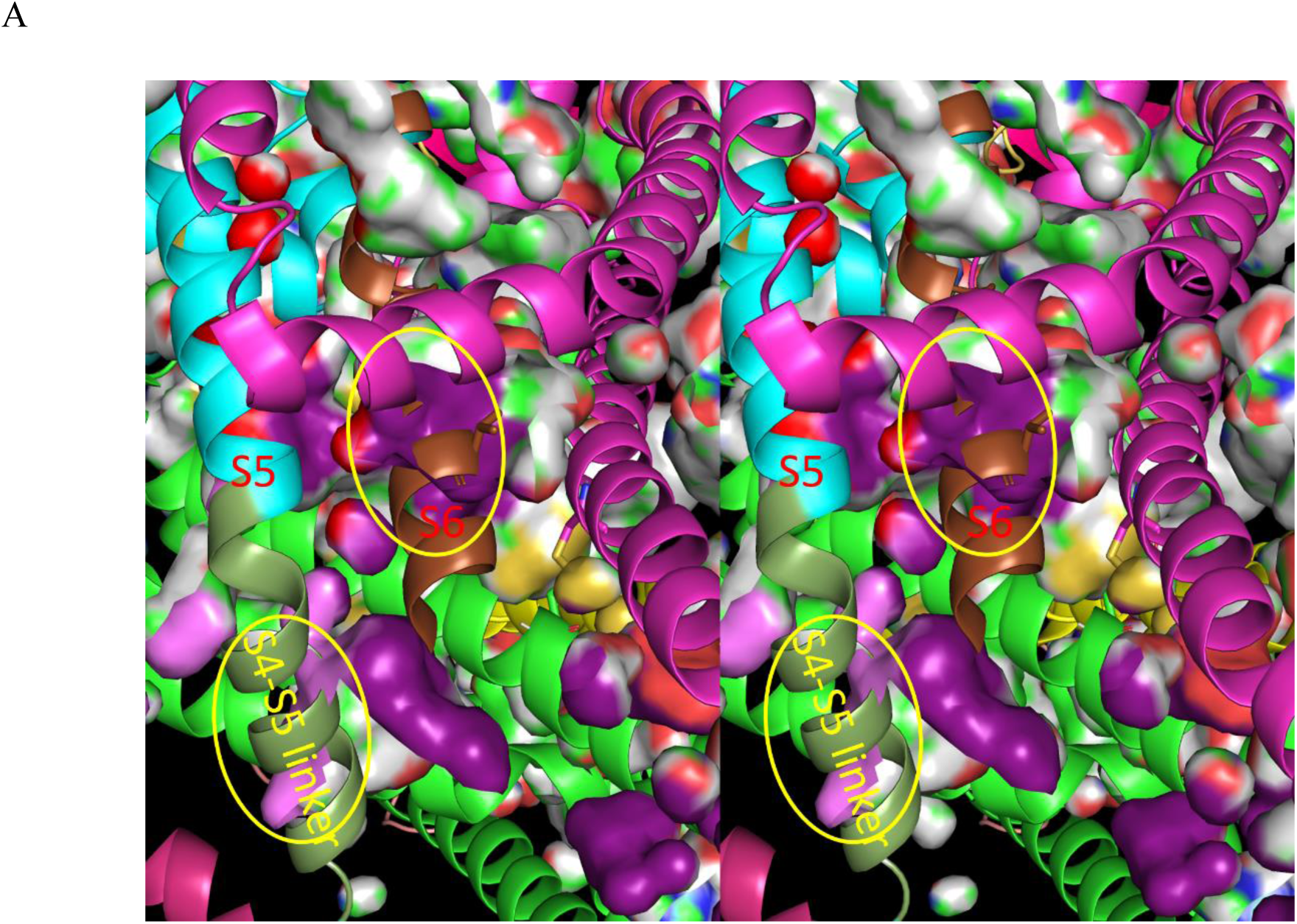
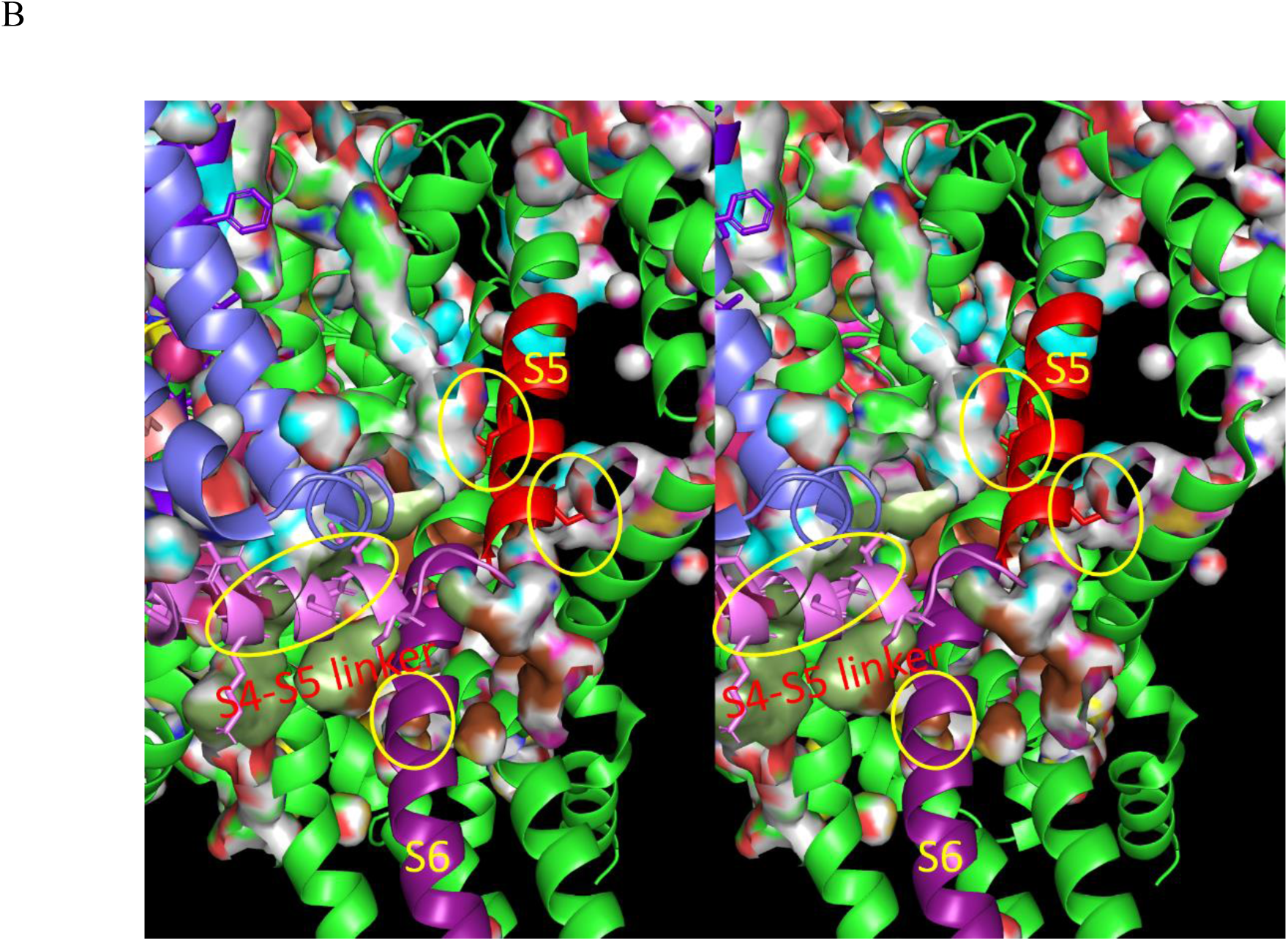
(A) Stereo view of the S4-S5 linker (olive cartoon) + S5 (cyan cartoon) + S6 (brown cartoon) of the deactivated structure (6N4R) overlaid on the internal surfaces present within the activated structure (6N4Q). Both the S4-S5 linker and S6 occupy internal cavities (yellow circles). (B) Stereo view of the S4-S5 linker (pink cartoon) + S5 (red cartoon) + S6 (violet cartoon) of the activated structure (6N4Q) overlaid on the internal surfaces present within the deactivated structure (6N4R). The activated structure occupies four internal cavities in the deactivated structure (yellow circles).

**Figure 37.**
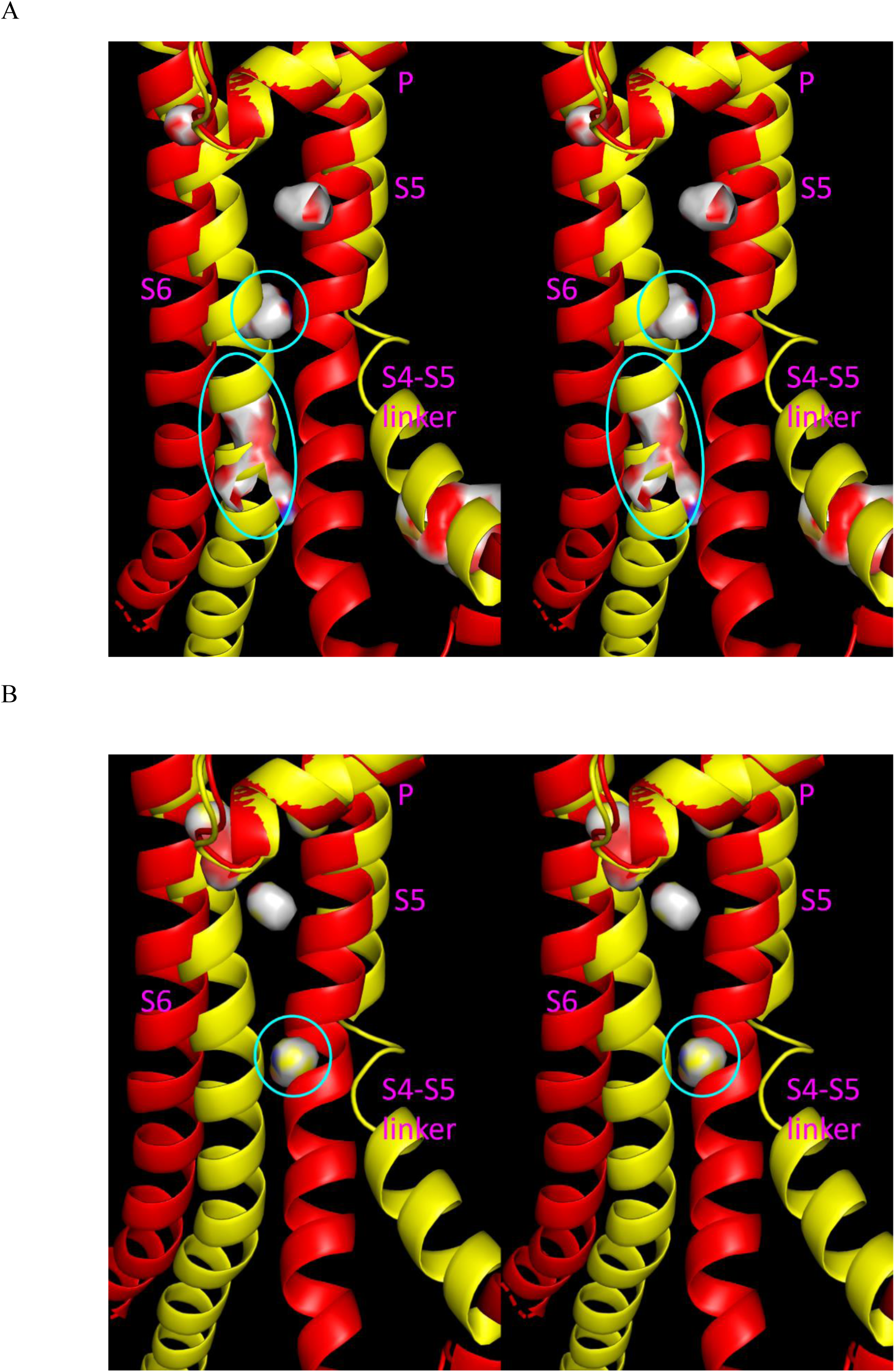
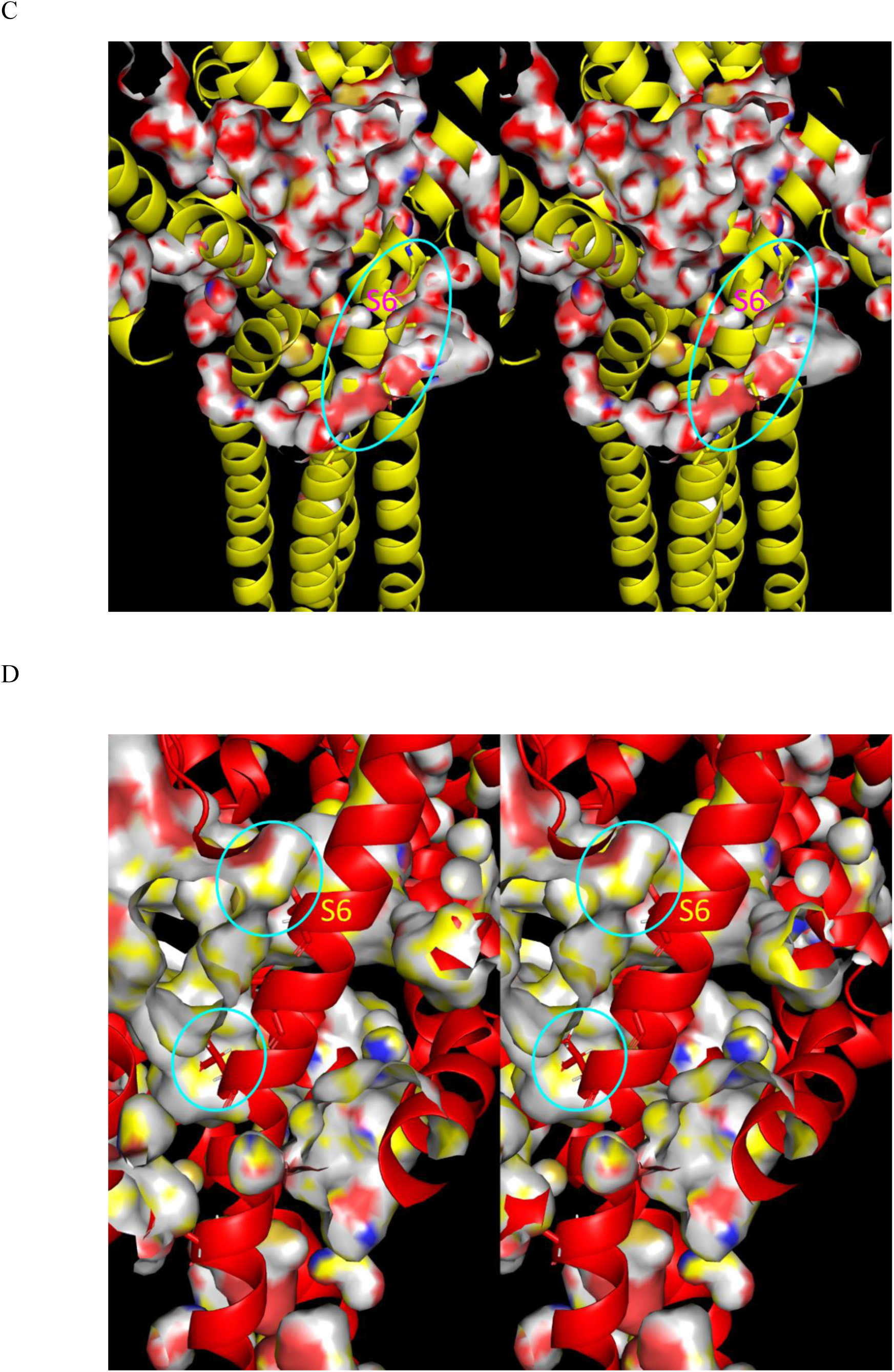
Stereo views of the overlaid activated/open and deactivated/closed states of S5 and S6. (A) The activated S6 occupies an internal cavity present in the deactivated structure. (B) The deactivated S5 occupies an internal cavity present in the activated structure. (C) Same as A, except showing the internal cavities within the entire pore domain. (D) Same as B, except showing the internal cavities within the entire pore domain.

**Figure 38.**
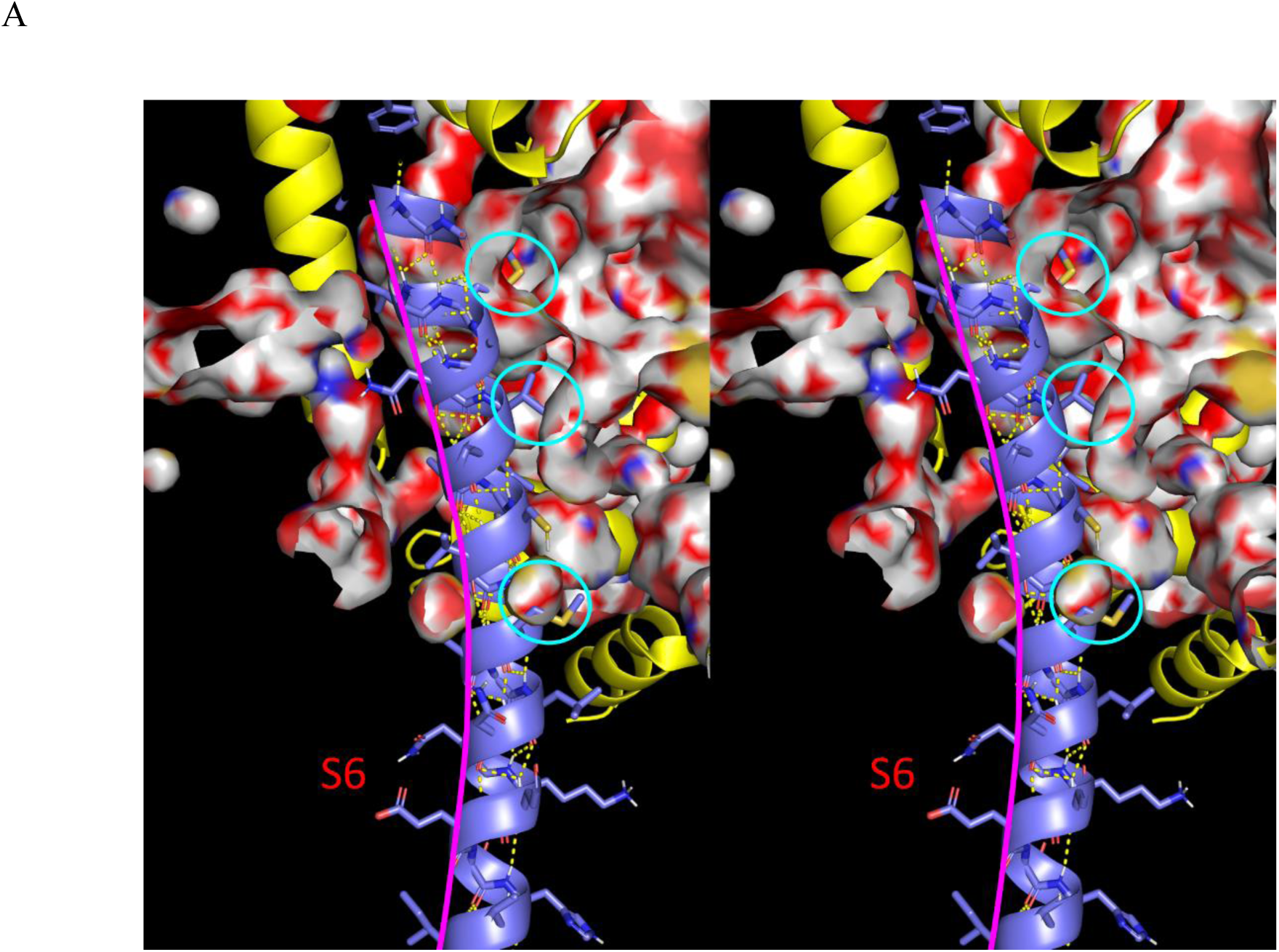
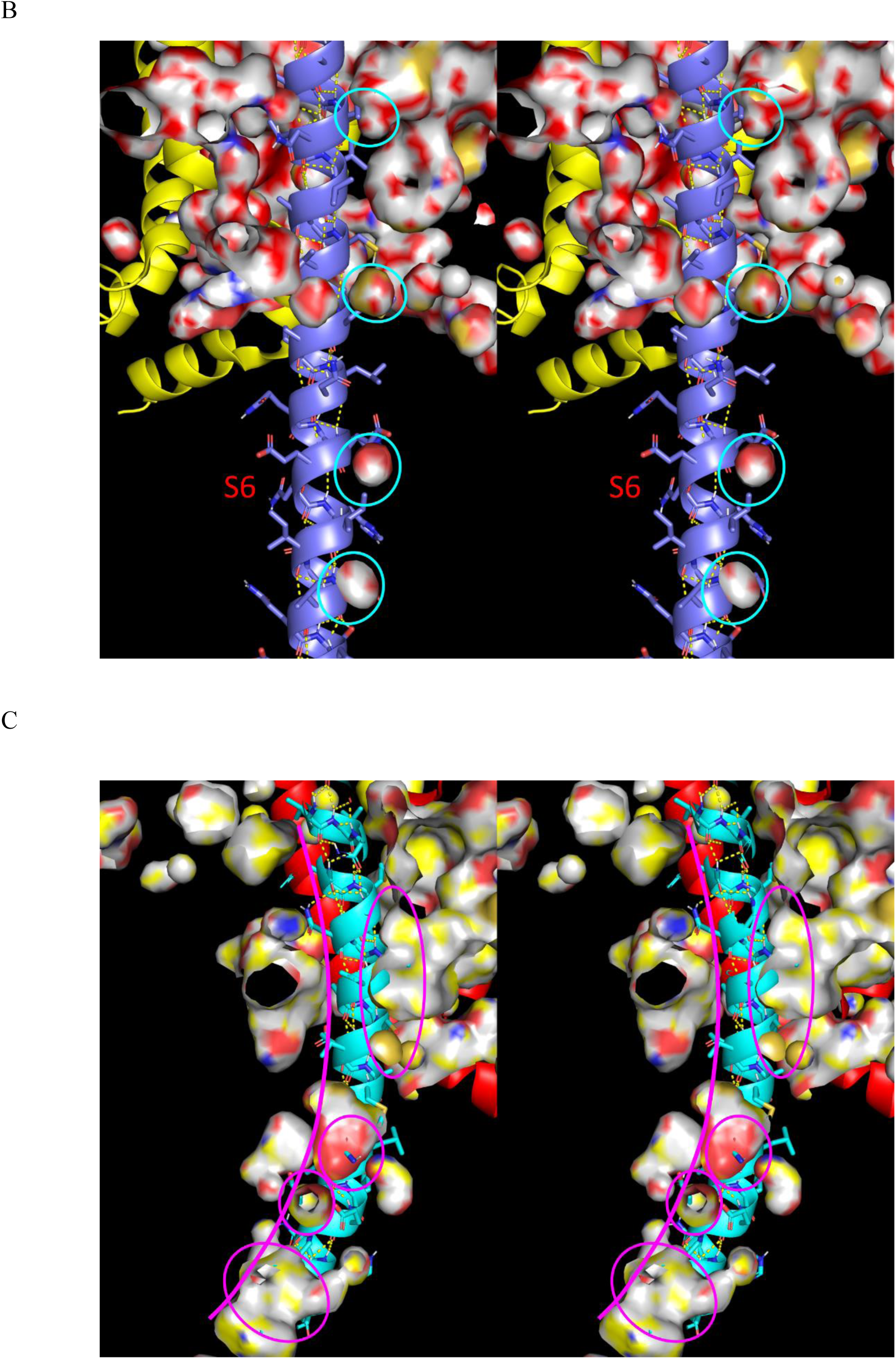
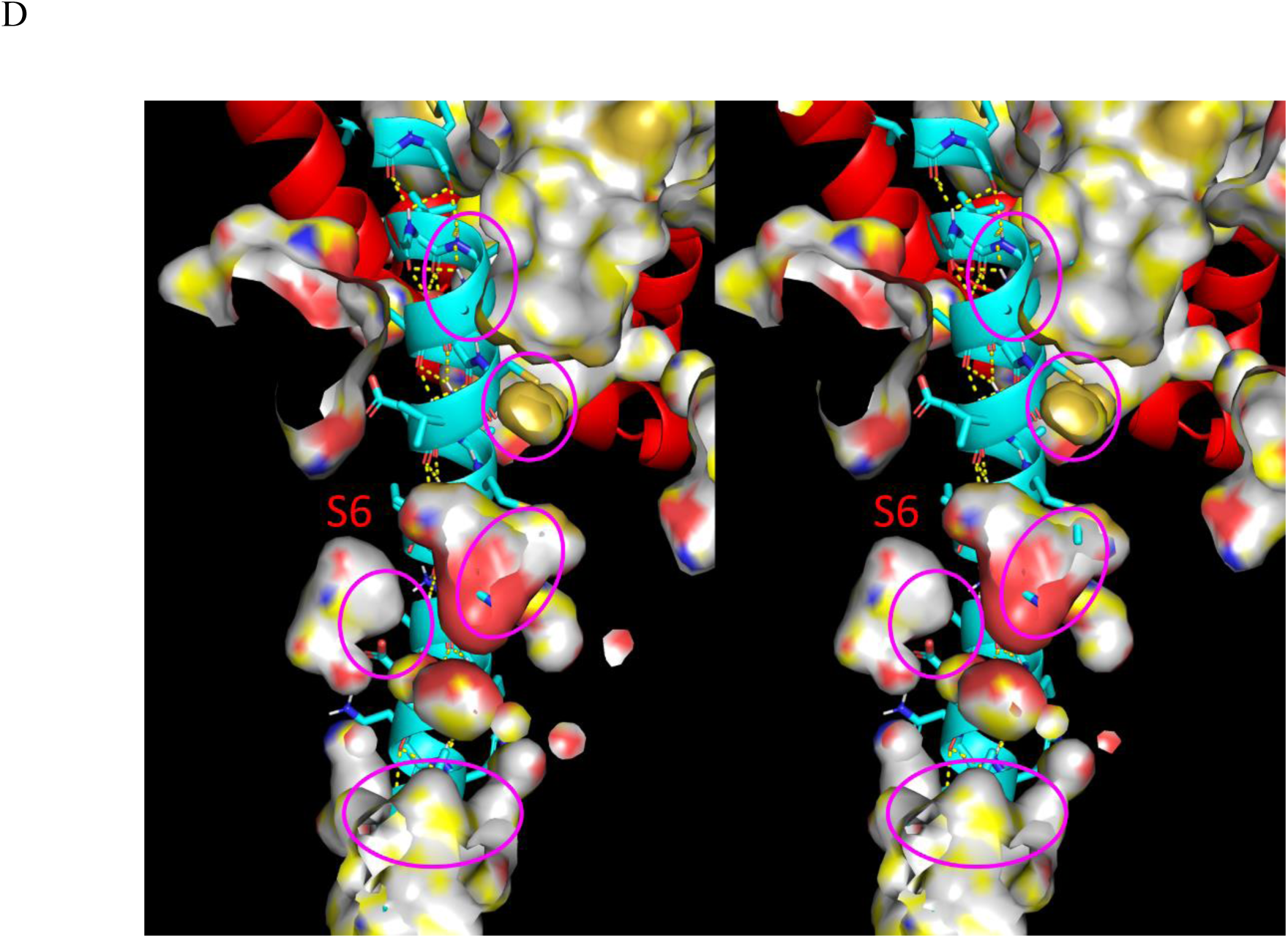
Stereo views of the tropimers within S6 of each state vis-à-vis the internal cavities of the opposite state in the Na_v_1.7 tetramer. The helix clearly follows the internal solvation cavities. The helical curvature within the tropimeric dyads is apparent. (A) S6 of the activated pore domain overlaid on the internal surfaces of the deactivated domain. (B) Same as A, except from a different vantage point, showing two additional internal cavities. (C) S6 of the deactivated pore domain over the internal surfaces of the activated domain. (D) Same as C, except from a different vantage point.

## Discussion

Our findings on GPCRs and VGCCs reported here further reinforces our previous findings on M^pro^ and cereblon,^21,22^ suggesting that:

1. Non-covalent <u>intermolecular</u> rearrangements of biomolecules are powered principally by the non-equilibrium expulsion of H-bond depleted solvation from their external surfaces. Bound states are stabilized on timescales governed by the resolvation cost 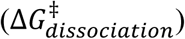 of the dissociated partners (where t_1/2_ = ln(2)/k_off_).
2. Non-covalent <u>intramolecular</u> state transitions (exiting from one state and entering another at rates denoted as k_out_ and k_in_, respectively) are powered principally by the non-equilibrium expulsion of H-bond depleted solvation as occupancy shifts from one set of internal cavities to another, and the vacated cavities resolvate in their wake. Intramolecular states are likewise stabilized on timescales governed by the resolvation cost(s) of the successor state(s) (where t_1/2_ = ln(2)/k_out_). We postulate that the magnitudes of unfavorable solvation free energy within the internal cavities of GPCRs and VGCCs have been optimized evolutionarily for achieving functionally relevant t_1/2_s.
3. Conservation of internal cavities and high energy solvation during intramolecular rearrangements ensures that free volume and power are available for further rearrangements that may operate cyclically (H-bond depleted solvation merely relocates during such rearrangements). Examples include enzymes that cycle between substrate binding/product release and complexation with additional partners;^21,22^ GPCRs that cycle between agonist binding/activation/G-protein binding/activation and agonist and G-protein dissociation/deactivation; VGCCs that cycle between activated/conducting and deactivated/inactivated/non-conducting states as a function of membrane potential state.
4. H-bond depleted solvation serves as the “muscles” of molecular structure, supporting the “weight” of functional states against thermal/entropic decay on a regional basis and powering the functional rearrangements thereof (noting that perturbation energies are limited to transducing local, but not long-range structural effects). Protein substructures thread through internal cavities present in the penultimate state (as illustrated by the S6 of Na_v_1.7 in in Figure 37). Interatomic contacts (van der Waals, electrostatic, solute H-bonds and halogen bonds, π-cation, π-stacking, etc.) are far too weak for this purpose.^20,21,44^
5. Access to H-bond <u>depleted</u> solvation during intra- and intermolecular rearrangements is governed by the desolvation cost of H-bond <u>enriched</u> solvation incurred during the rearrangement (i.e., 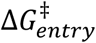). This contribution was not assessed in the present work, which can in principle be approached using our WATMD calculations.^21,22^
6. The dynamic structural behaviors of solute-solvation systems are governed by their <u>mutual</u> interactions, and as such, behave in a highly <u>non-linear</u> fashion. Solutes perturb the H-bond network/”field” of their solvation non-isotropically, which in turn, perturbs solute structure so as to minimize the overall H-bond free energy of the system (analogous to the mutual interactions between space and heavenly bodies giving rise to gravitational attraction). The dynamic structural behaviors of solute-solvation systems (analogous to solar systems/galaxies) “emerge” from the non-linearity of the interactions, which precludes the prediction of such behaviors using computational force-field and molecular descriptor-based approaches that do not properly account for the intricate solute-solvation “dance” governing non-equilibrium structure-free energy relationships.

Unlike our previous studies on M^pro^ and cereblon,^21,22^ we did not calculate the solvation fields of the β_2_-AR and Na_v_1.7 structures, instead inferring the locations of internal solvation solely from the internal cavity surfaces qualitatively mapped with PyMol (default surface settings), together with crystal water molecules present in the deactivated structure (2RH1). Our overall findings suggest that proteins are “hydro-powered”, “water-breathing” entities that rearrange on the basis of internal water cavities that transiently form and decay in response to inter- or extra-molecular signals/perturbations (e.g., Δψ_*m*_(*t*)).^22^ Internal solvation operates within a Goldilocks zone of free energy (where structures crossing the upper threshold do not form, and those crossing the lower threshold do not rearrange) that depends on a dynamic balancing act between H-bond enriched and depleted solvation and other contributions (e.g., Δψ_*m*_(*t*), Δψ_*d*_(*t*)).^22^ Structural state distributions are tipped in one direction or another by perturbations, such as agonist-GPCR binding (H-bonding with TMH3, TMH5, and TMH7), membrane depolarization/repolarization acting directly on the VSD, and putatively the membrane dipole potential acting directly on the pore domain (structural rearrangements are assisted, but not powered by such perturbations).

Our analysis was greatly hampered by the high variability present in the crystal and cryo-EM structures of even closely related GPCRs and Na_v_ and K_v_ family members, necessitating comparison of the structural features and internal cavities of all but the same unperturbed protein captured in its activated and deactivated states—namely, the β_2_-AR and Na_v_1.7 (noting that the internal solvation and key features of the activated and deactivated states of the VS and pore domains of the latter protein are self-consistent, despite the use of a non-native VSD construct and the inclusion of ProTx2). We attempted several overlay models of the β_2_-AR, including the global all-atom alignment and alignment about single and subsets of TMHs. However, only the ECL2 alignment (which putatively serves as the anchor for the induvial helices of the 7-TMH bundle) resulted in a meaningful relationship between the internal solvation in the activated and deactivated states of the receptor. The reciprocal relationship between the qualitatively mapped internal cavities in the activated and deactivated forms of the receptor and VGCC is remarkable, and leads to the conclusion that the shapes, sizes, and positions of these cavities constitute a major objective in protein folding.

## Conclusion

We showed previously that the sterically-accessible non-covalent inter- and intramolecular states and state transitions of <u>globular</u> proteins are powered principally by the H-bond free energy of solvating water.^45^ Here, we have extended these studies to the structures and rearreangements of <u>membrane</u> proteins, exemplified by the activation and deactivation of class A GPCRs (the β_2_-AR) and VGCCs (Na_v_1.7). Our findings explain the activation and deactivation mechanisms of the β_2_-AR and the means by which they are powered by desolvation and resolvation of internal cavities that guide the underlying rearrangements. The TMHs thread through and desolvate these cavities during activation, which are subsequently vacated and resolvated during deactivation. The solvation free energy is alternately tipped toward lower energy H-bond depleted water in the activated and deactivated forms in the presence versus absence of concurrent H-bonds between bound agonists and specific polar side chains on TMH3, TMH5, and TMH7. Our findings also explain the activating and deactivating rearrangements of the VSD and pore domain of Na_v_1.7, which are likewise powered and guided by internal cavities that are alternately occupied and vacated in each state. The solvation free energy is likewise alternately tipped toward lower energy H-bond depleted water in the polarized and depolarized states of the channel (corresponding to the down and up positions of S4). Furthermore, certain TMHs in both GPCRs and VGCCs curve via unilateral disruption of IHHBs within contiguous helical dyads (referred to in this work as “tropimers”). Threading of the TMHs through the internal water cavities of GPCRs and VGCCs is achieved via both pivoting and helical curvature. High specificity of such rearrangements is enforced by evolutionarily determined internal cavity (and virtual internal cavity) positions generated during protein folding.

## Notes

### Competing Interest Statement

The authors have declared no competing interest.

### Summary of Updates

Minor corrections to the text.

